# Enhancer grammar of liver cell types and hepatocyte zonation states

**DOI:** 10.1101/2022.12.08.519575

**Authors:** Carmen Bravo González-Blas, Irina Matetovici, Hanne Hillen, Ibrahim Ihsan Taskiran, Roel Vandepoel, Valerie Christiaens, Leticia Sansores-García, Elisabeth Verboven, Gert Hulselmans, Suresh Poovathingal, Jonas Demeulemeester, Nikoleta Psatha, David Mauduit, Georg Halder, Stein Aerts

## Abstract

Cell type identity is encoded by gene regulatory networks (GRN), in which transcription factors (TFs) bind to enhancers to regulate target gene expression. In the mammalian liver, lineage TFs have been characterized for the main cell types, including hepatocytes. Hepatocytes cover a relatively broad cellular state space, as they differ significantly in their metabolic state, and function, depending on their position with respect to the central or portal vein in a liver lobule. It is unclear whether this spatially defined cellular state space, called zonation, is also governed by a well-defined gene regulatory code. To address this challenge, we have mapped enhancer-GRNs across liver cell types at high resolution, using a combination of single cell multiomics, spatial omics, GRN inference, and deep learning. We found that cell state changes in transcription and chromatin accessibility in hepatocytes, liver sinusoidal endothelial cells and hepatic stellate cells depend on zonation. Enhancer-GRN mapping suggests that zonation states in hepatocytes are driven by the repressors Tcf7l1 and Tbx3, that modulate the core hepatocyte GRN, controlled by Hnf4a, Cebpa, Hnf1a, Onecut1 and Foxa1, among others. To investigate how these TFs cooperate with cell type TFs, we performed an *in vivo* massively parallel reporter assay on 12,000 hepatocyte enhancers and used these data to train a hierarchical deep learning model (called DeepLiver) that exploits both enhancer accessibility and activity. DeepLiver confirms Cebpa, Onecut, Foxa1, Hnf1a and Hnf4a as drivers of enhancer specificity in hepatocytes; Tcf7l1/2 and Tbx3 as regulators of the zonation state; and Hnf4a, Hnf1a, AP-1 and Ets as activators. Finally, taking advantage of *in silico* mutagenesis predictions from DeepLiver and enhancer assays, we confirmed that the destruction of Tcf7l1/2 or Tbx3 motifs in zonated enhancers abrogates their zonation bias. Our study provides a multi-modal understanding of the regulatory code underlying hepatocyte identity and their zonation state, that can be exploited to engineer enhancers with specific activity levels and zonation patterns.

## Introduction

Gene Regulatory Networks (GRNs), in which transcription factors (TFs) bind to genomic enhancers and promoters to regulate target gene expression, underlie cell identity. The advent of single cell technologies has provided unprecedented tools to probe mechanisms of cell type specific gene regulatory networks^1–3^. For example, it is now possible to measure gene expression (scRNA-seq) and accessible chromatin (scATAC-seq) in the same cells.

Single-cell data have also challenged the definition of cellular identity and cell types. Historically, cells were classified based on their phenotype (e.g. function, shape, location, and interactions), but single cell technologies have enabled a deeper characterization of their physiology and molecular components, such as proteins, transcripts, and chromatin. Single-cell approaches enable a precise, albeit sparse, description of individual cells in contrast to bulk analyses where many cells are pooled together that can mask signals from rare subpopulations. Single cell methods thus generate an unbiased view of the entire cell state space of a tissue^4, 5^.

While the historic definition of cell types is now considered artificial – requiring an arbitrary threshold to classify each cell on a discrete cell type category –, several studies now define a cell type as a (continuous) set of cell states, which can be aligned with the range of cellular phenotypes resulting from the interaction of a cell type with its microenvironment^6^. Indeed, certain cell types can be affected by their spatial location in the tissue, as in the mammalian liver. The liver is composed of hexagonal structures called liver lobules, which have a portal triad in each of its vertices (including a portal vein, an artery, and a bile duct each) and are pierced in the middle by a central vein (Fig 1a). Blood flows into each lobule from the portal vein and artery and moves through the sinusoids along sheets of hepatocytes towards the central vein, where it drains and leaves the liver. The flow along the lobular axis from the portal vein to the central vein, creates a gradient of nutrients, oxygen, hormones, and morphogens that results in a highly variable environmental axis^7^. Hepatocyte function varies depending on the position along this porto-central axis, as they are exposed to different microenvironments, a phenomenon known as zonation^7^. For example, periportal hepatocytes are more involved in gluconeogenesis and beta-oxidation, while pericentral hepatocytes are more involved in glycolysis, lipogenesis, and detoxification^7^. Previous single-cell and spatial transcriptomics studies have shown that not only hepatocyte function, but also transcriptome, varies along this axis^8^. Yet, whether and how these variable states are encoded in the genome and how zonation interacts with the GRNs underlying hepatocyte identity is largely unknown.

**Figure 1.**
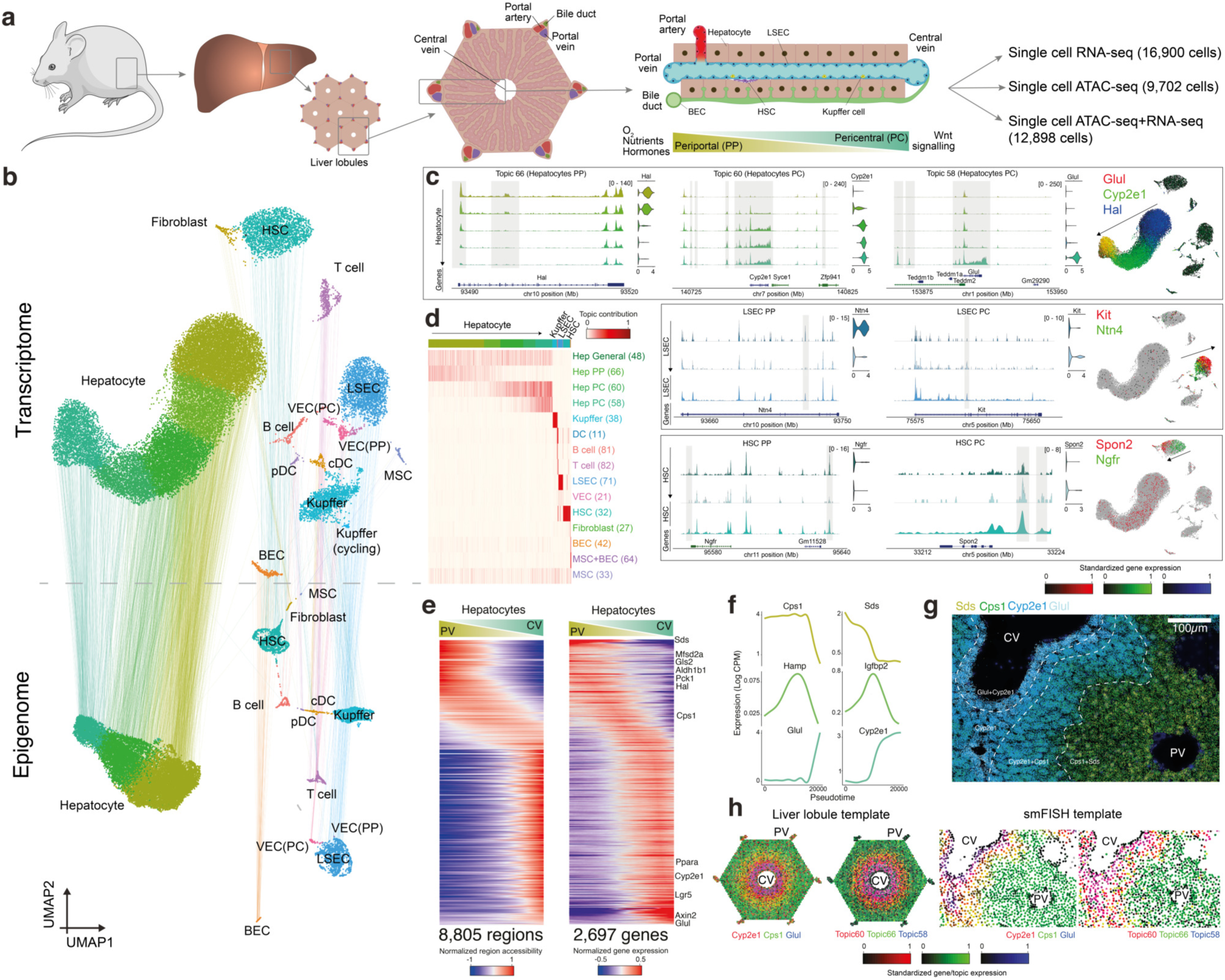
A spatial single cell multiome atlas of the mouse liver. **a.** Mouse liver overview and experimental set-up. The liver is composed of hexagonal structures called liver lobules, in which blood flows from the portal veins and hepatic arteries and drains in the central vein, creating a gradient of oxygen, nutrients, hormones and morphogens (e.g. Wnt). **b.** Transcriptome and epigenome based UMAPs (29,798 and 22,600 cells, respectively). Lines linking the UMAPs connect the transcriptome and the epigenome UMAP positions from the same cell (profiled by single cell multiomics). **c.** Pseudobulk chromatin profiles at different gene loci for hepatocyte, LSEC and HSC zonation state, accompanied by violin plots representing the normalized gene expression of the relevant gene in each class. UMAPs show the gene expression of the relevant genes with RGB encoding. **d.** Cell-topic contribution heatmap. **e.** Normalized region accessibility and gene expression zonation heatmaps. Cells are ordered by pseudotime (from periportal to pericentral) and regions and genes affected by zonation are shown (8,805 regions and 2,697 genes). **f.** GAM fitted gene expression profiles for selected genes along the zonation pseudotime. **g.** Liver section image showing smFISH profiles for Glul, Cyp2e1, Cps1 and Sds. **h.** ScoMAP liver lobule and smFISH colored by gene expression and topic contribution using RGB encoding.

A challenge to infer GRNs from single cell data is to predict whether a regulatory region is active, as chromatin accessibility is necessary but not sufficient for activity^9^. Thus, a potentially large fraction of accessible regions, as captured by scATAC-seq, may not represent functional enhancers. In Drosophila, different studies have reported that ∼70-90% of accessible regions are active enhancers, when their accessibility is cell type specific^10–12^, but in mammals this ratio drops to ∼30%^13^. Nevertheless, new experimental approaches, such as Massively Parallel Reporter Assays (MPRAs), have been established to directly test and identify active enhancers^14–18^. On the computational side, we and others have recently shown the power of convolutional neural networks (CNN) to model, predict, and interpret an enhancer accessibility or activity from its sequence^12, 19–21^. In addition, methods for inferring enhancer-driven GRNs (eGRNs) that integrate enhancer sequence, chromatin accessibility and gene expression^22–24^ found that the accuracy to predict active enhancers increases by ∼10% compared to using accessibility alone^25^. Nevertheless, challenges remain in the identification and design of state-specific enhancers and their validation *in vivo* in mammals, most notably the identification of features or mutations that drive cell type specificity and activity level and their experimental validation in specific cell states.

In this work, we generated a spatial single cell multiome atlas from the mouse liver and used these data to infer cell-type and cell-state specific eGRNs. We then augmented this data set with enhancer activity data measured by an *in vivo* MPRA on 12,000 hepatocyte enhancers. Next, we built a hierarchical deep learning model, called DeepLiver, to predict enhancer accessibility, activity, and zonation state from the enhancer sequence. Finally, we use DeepLiver to modulate enhancer activity and zonation and validate our predictions using a novel cell-state specific MPRA. Our results can be explored as a resource on SCope^26^ (http://scope.aertslab.org/#/Bravo_et_al_Liver) and the UCSC Genome Browser^27^ (https://genome.ucsc.edu/s/cbravo/Bravo_et_al_Liver).

## Results

### Single cell multiomics and spatial transcriptomics recapitulate liver cell types and reveal zoned cell states

To characterize liver cell types and states at the transcriptome and chromatin level, we performed two 10x single cell multiomics (ATAC+RNA) experiments in the wild type mouse liver, resulting in a total of 12,898 high quality cells (see *Methods*). To improve resolution, we additionally performed four 10x single nuclei RNA-seq (snRNA-seq) and two 10x single nuclei ATAC-seq (snATAC-seq) experiments, which provided 16,900 single cell transcriptomes and 9,702 single cell ATAC profiles, respectively (Fig 1a). We first analyzed the scATAC-seq and scRNA-seq data independently, using VSN-Pipelines^28^ for the transcriptome data; and pycisTopic^24^ for the chromatin accessibility data (see *Methods*). While VSN-Pipelines implements Scanpy^29^, pycisTopic relies on a topic modelling algorithm called Latent Dirichlet Allocation^30^, which clusters cells and regions into regulatory topics simultaneously, without predefining discrete cell clusters (which is not trivial in dynamic or continuous populations). From the snRNA-seq data we obtained 5,863 UMIs and 2,377 expressed genes per cell on average. The snATAC-seq data yielded 486,888 accessible regions that were grouped into 82 topics (see *Methods*) with a mean of 12,083 unique fragments and 7,241 accessible regions per cell, a median TSS enrichment of 16.1, and a fraction of reads in peaks (FRIP) of 66%. Importantly, we observed similar overall quality between the multiome experiments and the independent assays (Fig S1-3). Both the snRNA-seq and the snATAC-seq data distinguished the same cell populations, corresponding to 14 different cell types (Fig 1b), including hepatocytes, hepatic stellate cells (HSC), liver sinusoidal endothelial cells (LSEC), biliary epithelial cells (BEC), Kupffer cells, periportal and pericentral vascular endothelial cells (VEC), mesothelial cells (MSC), fibroblasts, and other immune cells (e.g. T cell, B cell and plasmacytoid/ conventional dendritic cells). In addition, we found a significant correlation between the average chromatin accessibility around the genes and gene expression (p-value < 2.2x10^-16^, Fig S4).

In all animals, we found a unidirectional gradient within hepatocytes, corroborated both by gene expression and region accessibility (Fig 1c-f). In addition, the LSEC and HSC clusters are also represented as a gradient, reflected by distinct chromatin accessibility topics and gene expression (Fig 1c,d). To identify whether this gradient represents spatial variation along the porto-central axis, we performed single molecule Fluorescence *In Situ* Hybridization (smFISH) with a panel of 100 selected genes across cell types and cell states in the liver (Fig 1f, Fig S5, Table S1). In hepatocytes, we identified three major zones that agree with the three regulatory topics found across the porto-central axis in hepatocytes; a *Glul+* zone that comprises the hepatocytes surrounding the central vein (Topic 58), a *Cyp2e1+* zone that includes pericentral and mid-lobular hepatocytes (Topic 60) and a *Cps1+* zone that contains mid-lobular to periportal hepatocytes (Topic 66). We additionally identified a mid-lobular area where *Cyp2e1* and *Cps1* are co-expressed, also reflected by the overlap of topic 60 (pericentrally intermediate) and topic 66 (periportal) cell contributions. For each gene and region, we fitted a Generalized Additive Model (GAM) across the pseudospatially ordered cells (from periportal to pericentral hepatocytes, see *Methods*) and found 2,697 genes (out of 6,823 genes expressed in hepatocytes with at least 3 UMI counts in 10 cells) and 8,805 regions in hepatocytes (out of 14,005 shared hepatocyte regions) whose respective expression and accessibility varies significantly along the porto-central axis (Fig 1g, adjusted p-value < 0.01, see *Methods*). This integrated spatial and single-cell analysis also confirmed that *Ntn4+* LSECs and *Ngfr+* HSC were located periportally; while *Kit+* LSECs and *Spon2+* HSC were located pericentrally, as previously reported^31, 32^ (Fig S6). We also identified 220 and 275 genes and 281 and 475 regions that vary along the porto-central axis in HSC and LSECs, respectively (adjusted p-value < 0.01, see *Methods*). In addition, BEC clusters could be clearly located in the bile ducts; pericentral and periportal VECs surround the corresponding vessels, together with fibroblasts; and Kupffer cells are preferentially located in periportal and mid zones without a strong zonation pattern, in agreement with recent studies^33^. Other immune cell types (e.g. B cells, T cells) are located across all zones (Fig S5).

To map the whole transcriptome and epigenome into the smFISH spatial map, we implemented a new version of ScoMAP (Single-cell omics Mapping into spatial Axes using Pseudotemporal ordering)^11^, resulting in a simplified template of a liver lobule (Fig 1h, see *Methods*). To generate this map, we ordered the zonated cell types (Hepatocytes, LSECs and HSCs) based on their transcriptome (and/or chromatin accessibility) into one dimension (i.e., pseudotime) that corresponds to the porto-central axis in the liver. This approach is then also used to order the cells in the smFISH template (based on the expression of the selected 100 genes), while we order the liver lobule template cells based on their distance to the lobule center (i.e. central vein). For each cell type, we divide the real and template cells into bins based on their position along the pseudospatial axis. Finally, we projected single cells profiles into the position of the template cells matching their positional bin. Single cells form cell types that are not affected by their spatial location along the porto-central axis (e.g. BECs, Kupffer cells) were randomly mapped onto their annotated counterparts of the template. This approach enabled us to visualize gene expression and region accessibility, as well as regulatory topics in a simplified representation of a liver lobule that shows a good agreement between the hepatocyte transcriptomic and chromatin accessibility zones (Fig 1g,h).

We also identified an interesting batch effect in hepatocytes, related to differences in physiological state between the mice (Fig S7), including circadian rhythm, nutritional status, and hormone levels. For instance, two complementary topics (Topic 17 and Topic 75) represent different phases of the circadian rhythm, with enrichment for positive and negative regulation (adjusted p-value = 10^-19^ and 10^-7^, respectively); and four topics show enrichment for different GO terms related to hormone response and metabolism (Fig S8). Using publicly available scRNA-seq data on the mouse liver during different phases of the circadian rythm^34^, we confirmed that the animals that were fed *ad libitum* mapped to early zeitgeber timepoints, while samples for which food was removed the night before mapped to late zeitgeber timepoints (Fig S9-S10, see *Methods*).

Next, we examined signaling pathways that have been reported to be affected by liver zonation, including Ras and Wnt signaling, hypoxia and pituitary hormone responses^8^. Our analyses revealed that all these pathways change along the porto-central axis (adjusted p-value < 0.05). Particularly, hypoxia and Wnt target genes are upregulated pericentrally and repressed periportally; Ras target genes are upregulated periportally and repressed pericentrally, and pituitary hormone targets are repressed pericentrally. However, hypoxia, pituitary hormones and Ras signaling target gene expression is highly variable between individuals and circadian rhythm phases (adjusted p-value < 0.05); while Wnt signaling targets are consistently expressed across all samples (adjusted p-value = 0.17, Fig S9-S10). In summary, our spatial single cell multiome atlas of the mouse liver reveals that both cell type identity and cell states, such as zonation and circadian rhythm, are congruently encoded at both the transcriptome and chromatin accessibility level.

### A core enhancer-driven gene regulatory network for hepatocytes is modulated by zonated repressor TFs

To identify candidate TFs underlying the different cell types and zonation states in the liver, we performed motif enrichment analysis in the different regulatory topics and differentially accessible regions (DARs) across cell types using pycisTarget^24^ (see *Methods*). As motifs can often be linked to more than one TF (and frequently, to several members of the same family), we pruned the list of annotated TFs by requiring a correlation of TF expression with motif enrichment. This resulted in the identification of Hnf1a, Ppara, Nfi, Hnf4a, Cebpa, Foxa1 and Onecut1 motifs in regions accessible across all hepatocytes, and Tbx3 and Tcf7l1/2 motifs in regions accessible periportally and pericentrally, respectively (Fig 2a). Interestingly, Tbx3 and Tcf7l1 expression profiles are anticorrelated with the accessibility of their potential target regions. Indeed, Tbx3 is expressed only in pericentral hepatocytes, while its candidate target regions are only accessible periportally. Vice-versa, Tcf7l1 is expressed periportally, and its candidate target regions are accessible pericentrally, while Tcf7l2 is expressed in all hepatocytes (Fig 2a). Tcf7l1 is a paralog of Tcf7l2, the Wnt-effector TF that is active pericentrally^35^; this may suggest that Tcf7l1 and Tcf7l2 bind to the same motif: Tcf7l2 pericentrally for activation, and Tcf7l1 periportally for repression^36^. In addition, this approach identified PU.1, Runx, and Irf as candidate TFs for Kupffer cell enhancers^37^; Pax5 and Ebf1 for B cells^38^; Tbx21 for T cells^39^; Meis1, Maf and Gata4 for LSECs^40^; Mef2c and Lhx2 for HSC^41^; and Tead4, Sox9 and Hnf1b for BECs^42^, among others (Fig S11).

**Figure 2.**
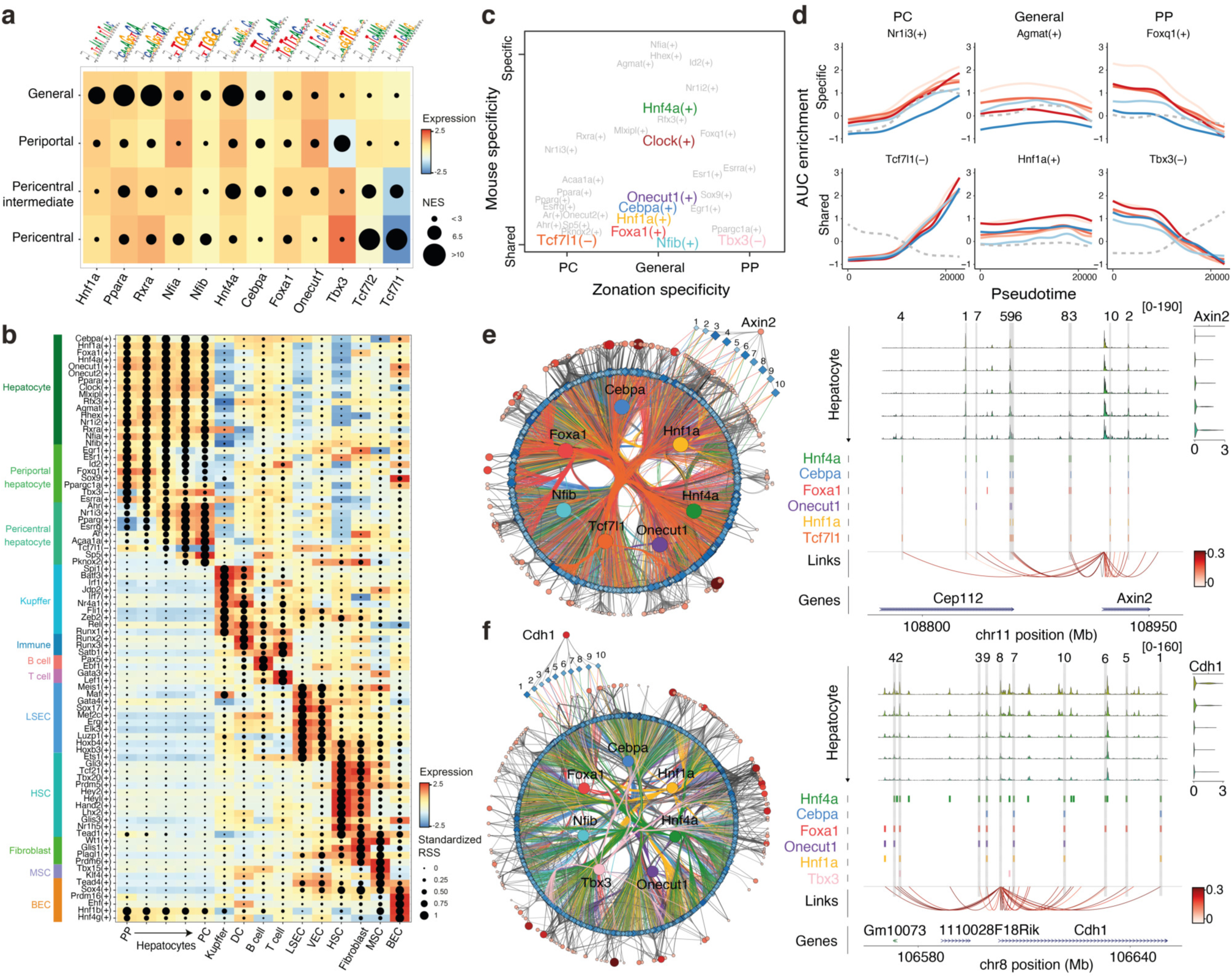
Liver zonation is mediated by repression. **a.** Dot plot showing the highest Normalized Enrichment Score (NES) as size for motifs linked to selected TFs in regions specifically accessible in different hepatocyte classes. Tiles are colored by the expression of the corresponding TF in that hepatocyte class. **b.** SCENIC+ eGRN enrichment dot plot. The gene-based Regulon Specificity Score (RSS) in the cell type is used as size, while color represents the TF expression in the corresponding cell type. **c.** eGRN AUC based PCA plot, where the first PC represents the zonation specificity and the second PC represents whether regulons are shared across mice or specific to certain mice. **d.** GAM fitted eGRN AUC profiles per mouse for selected regulons along the zonation pseudotime. The grey dotted line represents the GAM fitted TF expression profiles. **e.** Pericentral core hepatocyte eGRN, with 265 pericentral markers genes and 1,439 regions targeted by the selected core TFs (with CRM score > 3) and conserved across mice. Pseudobulk accessibility profiles on the *Axin2* locus are shown as example, with regions in the core eGRN highlighted in grey and numbered, TF binding sites, region to gene links colored by SCENIC+ correlation score and gene expression across the zonated hepatocytes classes (from PP to PC). **f.** Periportal core hepatocyte eGRN, with 175 periportal markers genes and 972 regions targeted by the selected core TFs (with CRM score > 3) and conserved across mice. Pseudobulk accessibility profiles on the *Cdh1* locus are shown as example, with regions in the core eGRN highlighted in grey and numbered, TF binding sites, region to gene links colored by SCENIC+ correlation score and gene expression across the zonated hepatocytes classes (from PP to PC).

Next, we examined how the predicted TF binding sites and enhancers are linked to candidate target genes. Following the SCENIC+ pipeline^24^ we compiled *enhancer-GRNs* using as input pycisTopic’s imputed chromatin accessibility, pycisTarget’s TF cistromes (i.e. a TF with its potential target regions) and the gene expression matrix. Using linear correlation and Gradient Boosting Machines, region-gene links (in a space 150kb upstream and downstream of each gene) and TF-gene relationships are inferred. Next, using an enrichment analysis approach, we assess whether the TF-coexpression module significantly overlaps with the genes recovered from the motif/region-based links, and subsequently retain the optimal set of target genes and regions for each TF. A TF with its set of predicted target enhancers and regions is called an enhancer-regulon (*eRegulon*).

This analysis revealed 180 regulons, including Spi1, Jdp2, Runx1 in Kupffer cells; Ebf1 and Pax5 in B cells; Gata3 and Lef1 in T cells; Gata4, Meis1 and Maf in LSECs; Lhx2 and Tead1 in HSC; Wt1 in fibroblasts; and Tead4, Dox4 and Hnf1b in BECs, among others, and in agreement with literature^37–42^. We find general hepatocyte-specific regulons for Cebpa, Hnf1a, Hnf4a, Onecut1, Foxa1 and Nfib^43–45^, and zonation-associated regulons such as Esr1 and Sox9 periportally^46, 47^ and Pparg and Ar pericentrally^48, 49^. Importantly, SCENIC+ identifies Tbx3 and Tcf7l1 as transcriptional repressors in pericentral and periportal hepatocytes, targeting 193 and 520 regions and 77 and 119 genes, respectively (Fig 2b, Fig S12). In other words, the chromatin regions where accessibility is negatively associated with Tbx3 expression are located nearby genes that are anti-expressed with Tbx3 (same for Tcf7l1). Furthermore, we validated the SCENIC+ eGRNs using publicly available Hi-C data^50^ and TF ChIP-seq^51^, and found that transcriptome prediction using SCENIC+ TFs as features is more accurate compared to random (see *Methods*, p-value < 2.2 x 10^-16^, Fig S13a-e).

As we previously observed transcriptomic and epigenomic differences between the mice depending on their physiological state, we performed Principal Component Analysis (PCA) of the eGRN enrichment scores to classify hepatocyte regulons based on their zonation state and mouse specificity (Fig 2c). This allowed us to identify regulons that depend on nutritional status, such as Agmat^52^ and Mlxipl^53^; on hormone levels, such as Nr1i2, Nr1i3 and Rxra^54^; and on circadian rhythm, such as Clock^55^. Some regulons were affected both by the physiological status of the mice and zonation, including Esrra and Foxq1 (periportal) and Ppara (pericentral). Among the general ‘core’ (shared across all mice) regulons we identified Onecut1, Cebpa, Hnf1a, Foxa1 and Nfib; while Tbx3 and Tcf7l1 are core pericentral and periportal (repressive) regulons, respectively (Fig 2d). These physiological states are not independent of the hepatocyte ‘core’ eGRN. For example, the Hnf4a regulon, with 3,442 target genes and 26,127 target regions, contributes to both the core and the physiological state dependent programs (Fig S13f-i). This cooperativity, or blending, of the hepatocyte GRN with TFs controlling cell state is even stronger for zonation: 94.8% and 89.6% of the target regions of Tbx3 and Tcf7l1, respectively, overlap with the target regions of at least one of the general core TFs (Hnf4a, Hnf1a, Cebpa, Onecut1, Foxa1, Nfib). This suggests that the hepatocyte zonation eGRN is a subset (or a layer) of the general hepatocyte eGRN. For example, among the candidate enhancers near the pericentral hepatocyte gene *Axin2* we find six Tcf7l1 target regions. Among the candidate enhancers near the periportal gene *Cdh1* we identify two Tbx3 target regions. In both cases, these regions are bound by additional core general TFs (Fig 2e-f). In summary, SCENIC+ identified Hnf4a, Hnf1a, Cebpa, Onecut1, Foxa1 and Nfib as core general hepatocyte TFs and the repressors Tbx3 and Tcf7l1 as key repressors of the zonation programs; together with additional networks related to the animal’s physiological state.

### Enhancer sequence determines activity in hepatocytes

The enhancer and GRN predictions we have made thus far were fundamentally based on gene expression and chromatin accessibility. However, chromatin accessibility is not necessarily always associated with enhancer activity^9^. To assess whether the predicted enhancers are active, we performed a Massively Parallel Reporter Assay (MPRA) using a previously published enhancer-barcoding strategy^56^ (Fig 3a). We selected 10,845 candidate regions based on accessibility, cloned them in a pooled fashion (see *Methods*), and transfected this library in the mouse liver (7 replicates) and human HepG2 cells (2 replicates) (see *Methods,* Fig 3a, Fig S14a-d, Table S2). Regions that are accessible in hepatocytes show significantly higher enhancer activity compared to shuffled sequences (Fig 3b,c), and replicate MPRA analyses are strongly correlated (R=0.82-1, Fig S14a). We used the shuffled sequences as background to derive an optimal activity cut-off (see *Methods*), which classified 2,913 enhancers as active in at least one of the two systems (806 only *in vivo,* 921 only in HepG2, 1,186 in both, adjusted p-value < 0.1, see *Methods*) and 4,285 regions as inactive. In other words, 40.5% of ATAC peaks are active by MPRA, in line with other studies^13^. Among the mouse regions that are active *in vivo* in the mouse liver, 64% are distal enhancers and 27% are promoters. In contrast, of the regions active in human HepG2 54% were promoters, suggesting either that distal enhancer activation is more difficult to recapitulate in HepG2 cells, or that distal enhancers are less conserved than promoters between mouse and human (Fig 3d, S14e-f, S15a).

**Figure 3.**
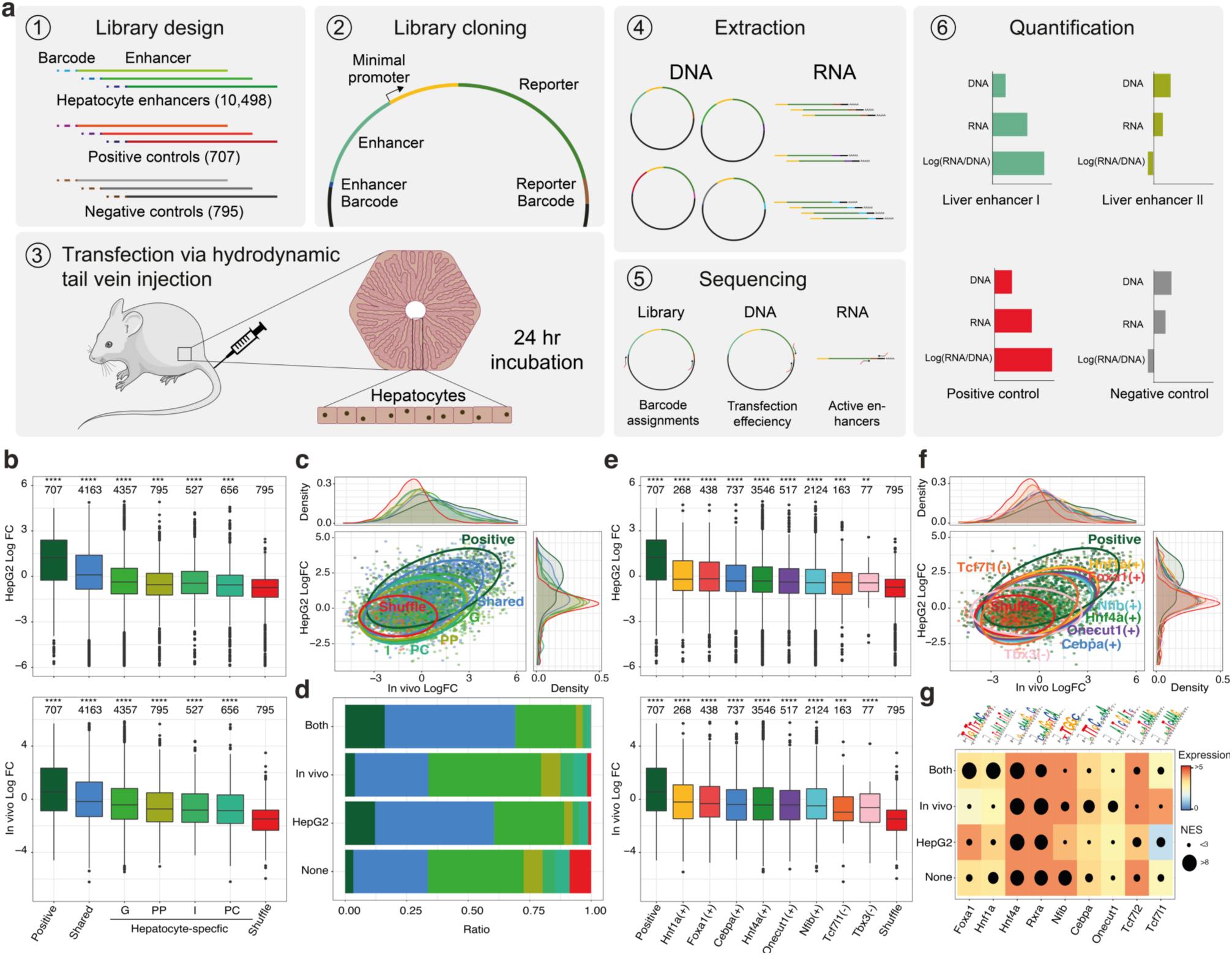
Massively Parallel Reporter Assays (MPRAs) in HepG2 and *in vivo* hepatocytes uncouple enhancer accessibility and activity. a. Schematic workflow of MPRA on the mouse liver. **b.** MPRA Log2 Fold-Change boxplots per enhancer class. The asterisks indicate the significance compared to random (****: p-value <= 0.0001, ***: p-value <= 0.001). G: General, PP: Periportal, I: Intermediate, PC: Pericentral. **c.** Correlation between Log2 Fold-Changes for high confidence enhancers (7,198) in Hepg2 and *in vivo* colored by enhancer type, with data ellipses per group. **d.** Proportion of enhancer classes per high confidence activity class. None: Not active (4,285), *In vivo*: Active only *in vivo* (806), HepG2: Active only in HepG2 (921), Both: Active in HepG2 and *in vivo* (1,186). **e.** MPRA Log2 Fold-Change boxplots per regulon. The asterisks indicate the significance compared to random (****: p-value <= 0.0001, ***: p-value <= 0.001, **: p-value <= 0.01). **f.** Correlation between Log 2Fold-Changes for high quality enhancers (7,198) in Hepg2 and *in vivo* colored by regulon, with data ellipses per group. **g.** Dot plot showing the highest Normalized Enrichment Score (NES) as size for motifs linked to selected TFs in regions in the different enhancer (MPRA) activity classes colored by the expression of the corresponding TF in HepG2, *in vivo* or as average (Both and None).

The SCENIC+ predicted target regions of Hnf1a, Hnf4a, Foxa1, Cebpa, Onecut1, Nfib and Tcf7l1 are all significantly more active compared to shuffled regions, with 45%, 39%, 43%, 39%, 35%, 32% and 26% of their predicted target regions active, respectively (Fig 3e,f). Tbx3 target regions are more active *in vivo* (20% and 5% of the regions are active *in vivo* and HepG2, respectively). In agreement with this, motif enrichment analysis of active versus inactive regions, followed by a classifier-based feature selection using Random Forest models, identified Hnf1a, Hnf4a, Foxa1, Creb, and AP-1 motifs as determining features in active enhancers (Fig 3g, S15, see *Methods*). This motif-based classifier predicts enhancer activity with AUROC of 0.71 (random AUROC 0.54) and AUPR of 0.44 (random AUPR 0.27, Fig S15). In summary, around 40% of accessible regions in hepatocytes are active, and the enhancer sequence is not only predictive of enhancer accessibility, but also of enhancer activity.

### DeepLiver decodes the grammar of enhancer specificity, activity, and zonation state

To further scrutinize how enhancer logic underlies enhancer activity and zonation, we trained a hierarchical Deep Learning model, named DeepLiver. We first trained a Convolutional Neural Network (CNN) to classify DNA sequences to the liver regulatory topics (called Topic-CNN), using 219,823 annotated regions as input. The weights learned by the Topic-CNN model were then used to initialize two additional CNNs one to predict MPRA activity *in vivo* (MPRA-CNN) and another to predict zonation (Zonation-CNN, using zonated accessibility classes as output variable). This transfer learning strategy overcomes the limited number of regions we have available for activity (4,215) and zonation (4,181 pericentral, 1,372 periportal, 12,122 general) (see Methods). For each model, the best epoch was selected based on their accuracy and loss on the test data (10% of the input data, Fig S16a). The three models resulted in higher AUROC and AUPR compared to a random control classifier, and topics associated with cell types had higher performance than low-contributing topics (Fig S16b-d). To validate DeepLiver predictions, we used a previously published MPRA data set performed on synthetic sequences *in vivo*^57^. These sequences were designed by adding different number of instances and combinations of motifs corresponding to TFs that are relevant to hepatocytes, including Hnf1a, Hnf4a (COUPTF), Cebpa, Onecut1 (HNF6) and Foxa1 (HNF3), among others. DeepLiver predictions correlate well with the experimental measurements (Topic-CNN R=0.63 (Topic 6) and MPRA-CNN R=0.68) (Fig S16e-j). Interestingly, we observed that for some TFs, especially Hnf1a, the number of motif instances correlates with the sequence activity (R=0.58). Sequences with more than two Hnf1a motif instances were predicted as highly active by DeepLiver, in agreement with experimental results (Fig S16h-j). Thus, DeepLiver assigns given DNA sequences to cell types (represented by topics) in the liver and predicts activity and zonation patterns of hepatocyte enhancers (Fig 4a).

**Figure 4.**
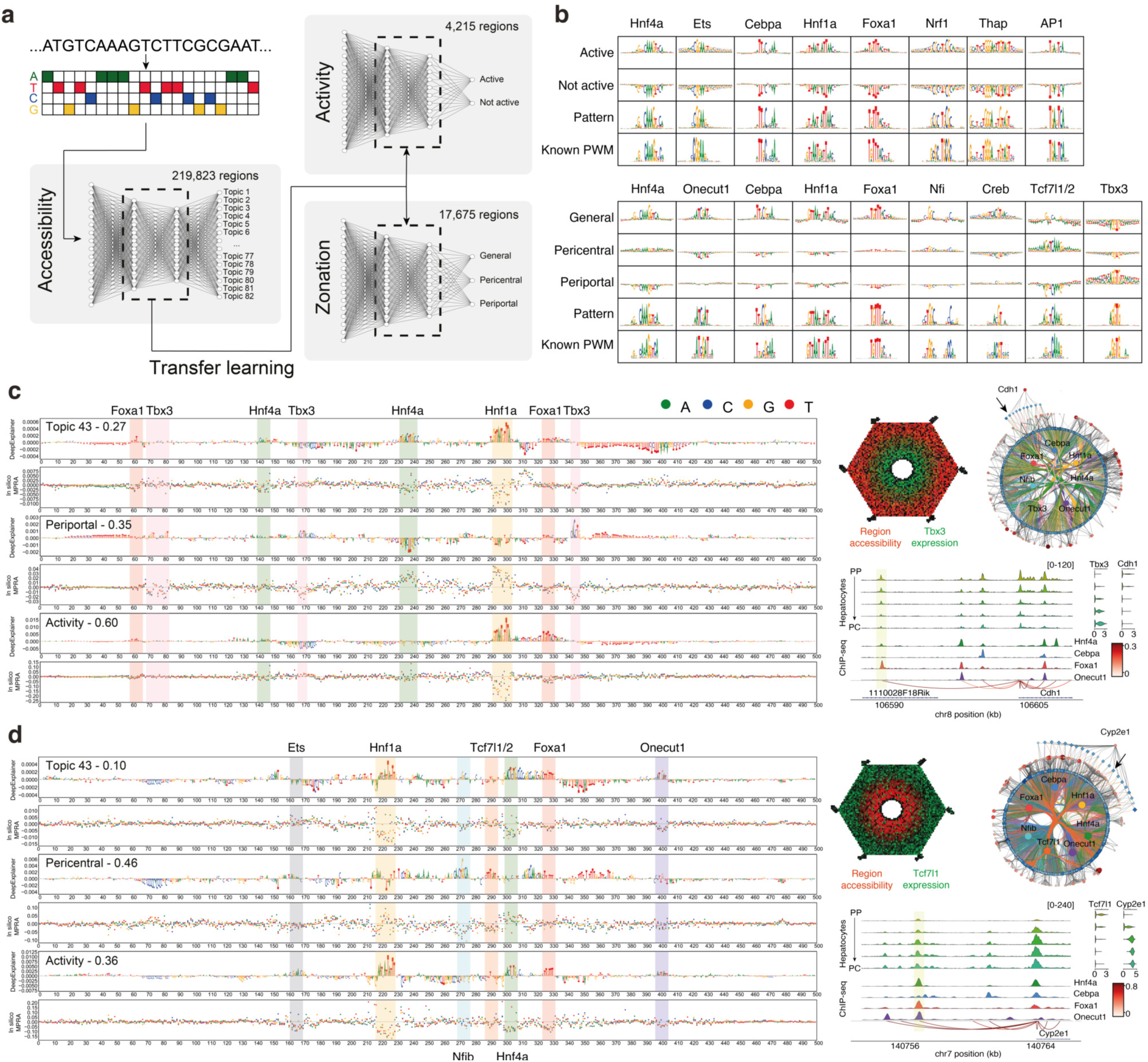
DeepLiver decodes enhancer grammar. **a.** DeepLiver overview. First, a CNN is trained to classify DNA sequences into their corresponding regulatory topic (219,823 sequences). The weights learned in the first model are used to initialize the activity and zonation models. The activity model classifies DNA sequences based on their MPRA activity *in vivo* (using 4,215 high confidence regions), while the zonation model classifies sequences based on their zonation pattern on hepatocytes (pericentral, periportal, or non zonated/general, using 17,675 regions for training). **b.** TF-MoDISco patterns identified in the activity and zonation models, with their contribution score per class and their most similar PWM from the cisTarget motif collection. **c.** DeepExplainer and saturation mutagenesis plots for the accessibility, zonation, and activity models on a Cdh1 periportal enhancer (chr8:106588720-106589220), with motifs highlighted. The accessibility model highlights the nucleotides that make the enhancer accessible on hepatocytes (versus other cell types in the liver); the zonation model, those that contribute to make the enhancer periportal and the activity model, those that have a role in their activity. On the right, the liver lobule template is colored by region accessibility (red) and Tbx3 expression (green) and the region is indicated by an arrow in the periportal core eGRN. Below pseudobulk accessibility profiles, ChIP-seq coverage (for Hnf4a, Cebpa, Foxa1 and Onecut1), SCENIC+ region to gene links colored by correlation score and Cdh1 and Tbx3 expression across the zonated hepatocytes classes (from PP to PC) are shown. The region is highlighted in yellow. **d.** DeepExplainer and saturation mutagenesis plots for the accessibility, zonation, and activity models on a Cyp2e1 pericentral enhancer (chr7:140756424-140756924). The accessibility model highlights the nucleotides that make the enhancer accessible on hepatocytes (versus other cell types in the liver); the zonation model, those that contribute to make the enhancer pericentral and the activity model, those that have a role in their activity. On the right, the liver lobule template is colored by region accessibility (red) and Tcf7l1 expression (green) and the region is indicated by an arrow in the pericentral core eGRN. Below pseudobulk accessibility profiles, ChIP-seq coverage (for Hnf4a, Cebpa, Foxa1 and Onecut1), SCENIC+ region to gene links colored by correlation score and Cyp2e1 and Tcf7l1 expression across the zonated hepatocytes classes (from PP to PC) are shown.

Next, we used DeepExplainer^58^ to assess the contribution of each nucleotide in the enhancer classification, and TF-MoDISco^59^ to identify motifs from recurring patterns in the contribution scores (Fig 4b). For the MPRA-CNN, we identified patterns promoting enhancer activity corresponding to Hnf4a, Cebpa, Hnf1a, Foxa1 and AP-1 motifs, and several promoter-related motifs such as Ets, Nrf1 and Thap. From the zonation model, motifs for Hnf4a, Onecut1, Cebpa, Hnf1a, Foxa1, Creb and Nfi are identified as regulators of accessibility across all hepatocytes; whereas Tcf7l1/2 and Tbx3 motifs are associated with pericentral and periportal accessibility, respectively. These results thus agree with the motifs identified by random forest feature selection from the MPRA data (Fig S15), and the hepatocyte (and zonation) regulators identified by SCENIC+ (Fig 2).

We further exploited DeepLiver to explore the effect of sequence variation on enhancer specificity, activity, and zonation. To validate DeepLiver *in silico* mutagenesis (see *Methods*), we compared the predicted effects of mutations with experimental saturation mutagenesis data on six enhancers from earlier studies (three each from *in vivo* and HepG2 studies)^14, 60^. DeepLiver predictions of the effect of enhancer mutations correlate with experimental results (R=0.36-0.75, Fig S17), and the strongest effect occurs when a mutation affects a TF binding site (causing gain or loss of activity). DeepLiver predictions outperform other methods tested by Kircher et al. (2019)^60^ in the HepG2 enhancers. For example, DeepLiver predicts F9 and SORT1 mutagenesis effects with a spearman correlation of 0.64 and 0.36, respectively, while for other methods ranges from -0.04 to 0.52 and -0.28 to 0.29 (Fig S17).

Next, we used TF-MoDISco patterns and SCENIC+ Position Weight Matrices (PWMs) to identify TF binding sites among the hepatocyte sequences (see *Methods).* We identified between 1,235 and 6,991 target regions for Tbx3, Tcf7l1, Foxa1, Hnf1a, Hnf4a, Nfib, Onecut1 and Cebpa, with good overlap with SCENIC+ predicted target regions (17%-70%, Fig S18a-c). Importantly, DeepLiver classifies Tbx3 target regions as periportal, Tcf7l1 and Nfib as pericentral/general, and Hnf4a, Foxa1, Hnf1a, Onecut1 and Cebpa regions as general (Fig S18a). In agreement with results above, Hnf1a target regions are predicted to be more active (median 0.52), while Nfib target regions have low activity scores (median 0.16, Fig S18b). To validate the predicted binding sites, we compared our predictions with previously published ChIP-seq data for Hnf4a, Cebpa, Foxa1 and Onecut1^51^, finding specific signal for the corresponding TF when centering the regions on the predicted binding site (Fig S18d). Finally, we also assessed distances between motif instances in overlapping regions. This showed that Tcf7l1 and Hnf4a often overlap, which is likely due to the similarity between the motifs (GATCAAAG and CAAAGTCA, respectively). On the other hand, Foxa1, Hnf1a, Cebpa, Nfib and Tbx3 are often located close to Hnf4a motifs (Fig S18e,f).

We then used DeepLiver to interpret enhancers in the ‘core’ pericentral and periportal eGRNs from SCENIC+, now at base pair resolution (Fig 4c,d, Supplementary Note 1). For instance, on a Cdh1 enhancer, DeepLiver finds Foxa1, Hnf4a and Hnf1a sites as drivers of enhancer accessibility and activity, while Tbx3 sites (one dimer motif, and two monomers) are predicted to make the enhancer periportal. In agreement, we find Hnf4a and Foxa1 ChIP-seq signal in this region, but no Cebpa nor Onecut1 ChIP-seq signal. Importantly, both accessibility of this enhancer, and Cdh1 gene expression, are anticorrelated with Tbx3 expression (-0.44 and -0.17, respectively). On a pericentral Cyp2e1 enhancer, DeepLiver identifies Hnf1a, Foxa1 and Onecut1 sites that contribute to enhancer accessibility and activity, and an Ets site that contributes to activity but not accessibility, as observed in other enhancers too (Supplementary Note 1). On the other hand, a Nfib site contributes to accessibility (but not activity), and a Tcf7l1/2 site is uniquely found in the zonation model, contributing to make the enhancer pericentral. In agreement, we observed Hnf4a, Foxa1 and Onecut1 ChIP-seq signal in this region. Tcf7l1 expression is anticorrelated with region accessibility and gene expression (-0.32 and -0.40). These observations suggest that Tbx3 and Tcf7l1 may repress these regions. In summary, DeepLiver decodes enhancer accessibility, activity, and zonation at base pair resolution, and can predict variants that modulate enhancer activity and zonation in hepatocytes.

### Computational and experimental validation of zonated repressor TFs

The DeepLiver model provides meaningful interpretations of hepatocyte enhancers and predicts that these enhancers consist of a core hepatocyte code, mixed with zonated repressor TFs, Tbx3 and Tcf7l1, which bias enhancer activity to either pericentral or periportal zones respectively. To test these predictions further, we first performed simulation experiments on the SCENIC+ network, following our previously published perturbation-simulation strategy^24^. Simulating Tbx3 or Tcf7l1 knockdown and overexpression in hepatocytes (see *Methods*), suggests that Tbx3 overexpression and Tcf7l1 knockdown can switch periportal hepatocytes to a pericentral state, while Tbx3 knockdown or Tcf7l1 overexpression can switch pericentral hepatocytes to a periportal state (Fig 5a-c). Simulated knockdown of these TFs results in upregulation of their predicted target genes, while their overexpression results in target gene downregulation. The SCENIC+ eGRN furthermore predicts Tbx3 and Tcf7l1 to directly repress each other. Consequently, the knockdown or overexpression of one of the TFs provokes the upregulation or downregulation of the other, respectively (and downregulation and upregulation of the target genes of the other as well; Fig 5c).

**Figure 5.**
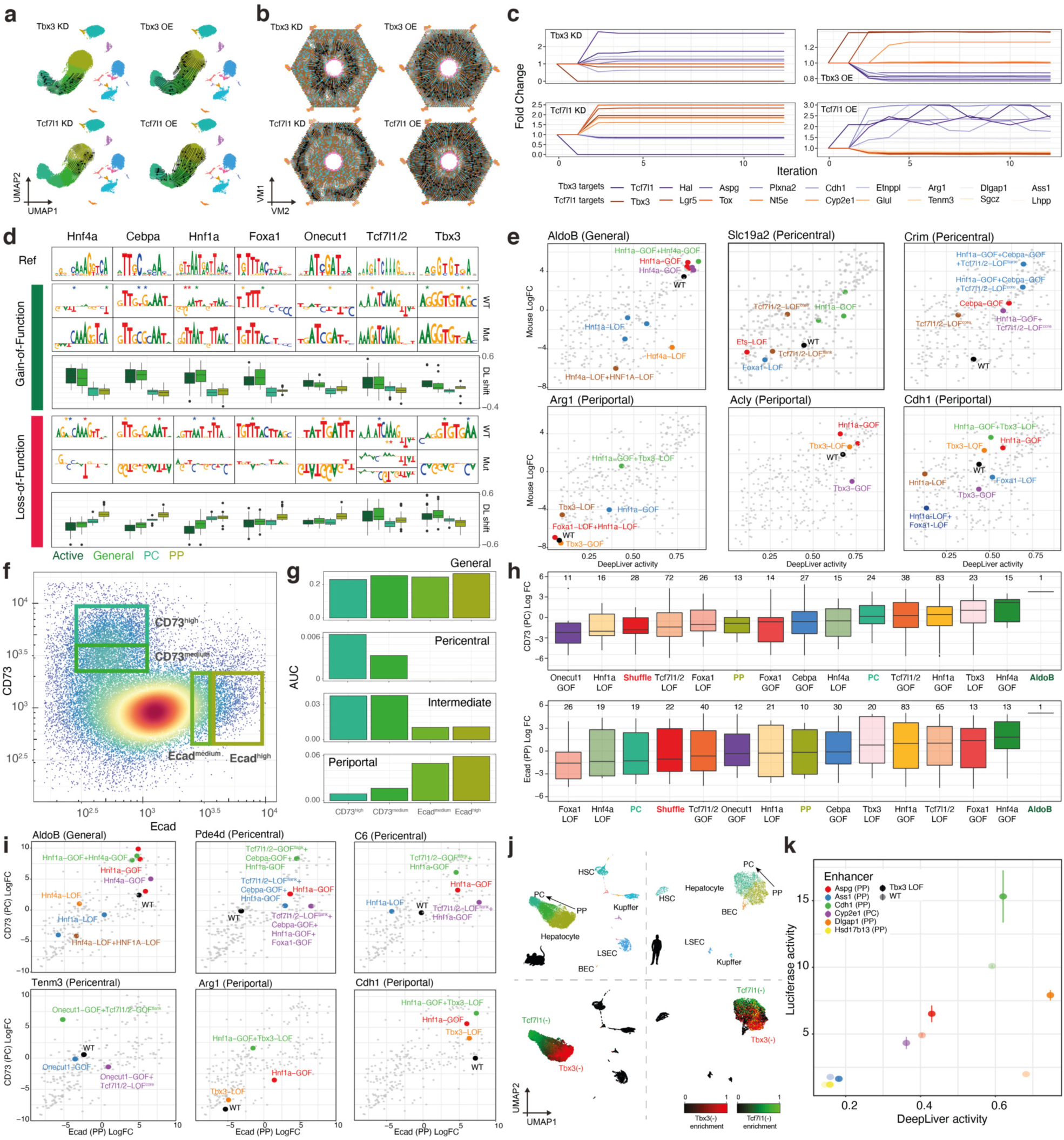
Tcf7l1/2 and Tbx3 modulate enhancer accessibility and activity along the porto-central axis. **a.** Simulated cellular shift on the snRNA-seq UMAP (29,798 cells) after Tbx3 or Tcf7l1 knockdown (KD) or overexpression (OE) represented using arrows. Arrows are shaded based on the distance travelled by each cell after the simulation. **b.** Simulated cellular shift on the liver lobule virtual map (4,498 metacells) after Tbx3 or Tcf7l1 knockdown (KD) or overexpression (OE) represented using arrows. Arrows are shaded based on the distance travelled by each cell after the simulation. **c.** Predicted fold change for selected genes (Tbx3 targets in purple and Tcf7l1 targets in orange) upon simulation of Tbx3 knockdown and Tcf7l1 overexpression in pericentral hepatocytes and Tcf7l1 knockdown and Tbx3 overexpression on periportal hepatocytes. **d.** Overview of DeepLiver-based sequence mutations introduced in the wild-type enhancers to shift activity and zonation patterns. These variants cause the appearance of improved motifs (Gain-Of-Function) or their destruction (Loss-Of-Function). The boxplots below each variant indicate DeepLiver’s predicted shift on activity (Active) or zonation (General, Pericentral or Periportal) scores. **e.** *In vivo* MPRA Log Fold change versus DeepLiver activity score with highlighted sequence variants for each enhancer. **f.** FACS of the selected cells according to the intensities of CD73 and Ecad. The rectangles indicate the selected bins along the porto-central axis. **g.** AUCell enrichment of the core general, pericentral, pericentral-intermediate and periportal regions on the sorted populations. **h.** *In vivo* MPRA Log Fold change on the aggregated CD73 and Ecad populations per variant class. Control and wild-type sequences are highlighted. **i.** CD73 MPRA Log Fold change versus Ecad MPRA Log Fold change with highlighted sequence variants for each enhancer. **j** Mouse and human liver scATAC-seq UMAP (22,600 and 6,366 cells, respectively) colored by cell type (top) and regulon enrichment (bottom). **k.** Luciferase activity in HepG2 versus DeepLiver activity scores for selected enhancers and their variants.

In a second experiment, we introduced specific mutations into a set of hepatocyte enhancers, guided by the DeepLiver model, and then measured their activity using *in vivo* MPRA. We selected 13 periportal and 21 pericentral enhancers, that are predicted to be repressed by Tbx3 (pericentrally) and Tcf7l1 (periportally), both by SCENIC+ and DeepLiver. We introduced GOF (Gain-of-Function) and LOF (Loss-of-Function) mutations affecting Hnf4a, Cebpa, Hnf1a and Foxa1 binding sites, and mutations of Tbx3 and Tcf7l1/2 motifs (Fig 5d, S19a-c), leading to a total of 455 sequences. The activities of these enhancer variants were first tested using bulk MPRA on the mouse liver and human HepG2 cells (see *Methods,* Fig S19d-g, Table S3), where GOF variants of Hnf4a, Hnf1a, Cebpa and Foxa1 indeed result in higher activity, as predicted by DeepLiver (Fig 5e, S19c,f-g). Variants of the predicted binding sites of the zonation TFs Tbx3 and Tcf7l1 also show changes, but these are more difficult to assess from these bulk experiments where periportal and pericentral hepatocytes are pooled (Fig 5e, S19f-g, S20, S21). To solve this problem, we performed MPRA experiments on FAC-sorted hepatocytes, sorted by zone, using pericentral and periportal surface proteins CD73 (encoded by *Nt5e*) and Ecad (encoded by *Cdh1*), respectively; following a previously published protocol^61^ (see *Methods*, Fig 5f, S22a). The sorted cell fractions indeed represent pericentral and periportal hepatocytes, as shown by bulk ATAC-seq profiles on the separate fractions, which agree with the scATAC-seq zonated profiles (Fig 5g). Next, we analyzed enhancer activity on the sorted fractions by MPRA (see *Methods*, Table S3). As expected, pericentral enhancers show higher activity in the pericentral fraction, and vice-versa (Fig 5h, Fig S22b,c). Hnf1a and Hnf4a GOF variants resulted in increased activity in both fractions, with milder effects for Cebpa and Foxa1 variants. Destruction of these motifs reduced enhancer activity compared to their wild-type counterparts (Fig 5h,i, Fig S22d). Tbx3 LOF and Tcf7l1 GOF resulted in an activity increase in the pericentral fraction and Tcf7l1 LOF in the periportal population (Fig 5h,i, Fig S22d, S23).

As a third validation experiment, we analyzed a public human liver scATAC-seq data set^62^, revealing that the predicted Tbx3 and Tcf7l1 repressive sites are conserved between species, with similar accessibility patterns along the porto-central axis (Fig 5j). Human HepG2 cells express Tbx3 and may therefore serve as a model to test the effect of mutating Tbx3 sites in hepatocyte enhancer sequences^63^. Indeed, the predicted Tbx3 binding sites by SCENIC+ are less active in HepG2 compared to *in vivo* (Fig 3e), and Tbx3 LOF variants showed increased activity in HepG2 by bulk MPRA (Fig S19f-g). We therefore tested five periportal enhancers and their Tbx3 LOF variants using luciferase reporter assays in HepG2 cells and included the pericentral Cyp2e1 enhancer as control (Fig 5k). As predicted by DeepLiver, Tbx3 LOF in inactive enhancers did not rescue the enhancers (Hsd17b13 and Ass1). However, the predicted active enhancers (Aspg, Cdh1 and Dlgap1) exhibited increased activity when the Tbx3 binding site was mutated. This indicates that these enhancers are directly repressed by Tbx3 though these sites. In summary, our results suggest that the grammar of hepatocyte enhancers that encodes their zonation pattern includes Tbx3 and Tcf7l1/2 binding sites, while Hnf1a and Hnf4a are the most relevant binding sites regarding activity.

## Discussion

Historically, the definition of a cell type has relied on discrete classifications based on morphology, size, function, location, or interaction with other cell types^6^. Single cell omics methods have revolutionized the definition of cell types, as they allow to profile up to thousands of snapshots of cell states in a tissue. We can now define cell types as a continuum of (reversible) cell states, that are often binarized based on statistical clustering of their transcriptome or epigenome. Yet, the discretization of dynamic populations is not a trivial task and is strongly affected by parameter selection. An alternative approach to characterize cell states is to study its underlying GRNs, and all the regulatory variations on that central theme. GRN inference from single cell data^22–24^ allows identifying specific and reproducible regulatory programs that underlie the different functions and behavior of a cell in a particular context. As such, one would expect that all cell states that compose a cell type share a core GRN, while variations on this program could be modulated by environmental cues. Using the mouse liver as a model system, we aimed at depicting the core identity, and the various cell states, of hepatocytes, alongside their gene regulatory programs.

We employed two complementary computational strategies to tackle this problem. Firstly, our recently developed method SCENIC+ identified a core hepatocyte GRN controlled by Hnf4a, Hnf1a, Cebpa, Foxa1, Nfib and Onecut1. As a subset of this program, we could disentangle novel mechanisms underlying hepatocyte zonation, ruled by the repressor TFs Tcf7l1 and Tbx3. SCENIC+ could identify these candidate repressors because their motif is significantly enriched in regulatory regions that are accessible in hepatocytes where the TF is not expressed, while they are inaccessible in hepatocytes where the TF is expressed. As a potential mechanism, how repressor binding could result in the absence of an ATAC peak, TF footprinting indeed suggests that direct repressor binding may occur within nucleosome-occupied regions, while activator binding is strongly associated with nucleosome depletion^64^. This illustrates the power of single-cell multi-omic profiling, whereby both positive and negative correlations between accessibility and gene expression can be exploited to infer regulatory interactions. Reassuringly, a comparison with the human liver confirmed that these two candidate repressors are expressed in a similar pattern, relative to hepatocyte zonation in human. Furthermore, in the eGRN, we identified a regulatory feedback loop between Tbx3 and Tcf7l1, which are predicted to repress each other. Tcf7l1 and Tbx3 are indeed well-known repressors in development^68, 69^, but had not been directly linked to hepatocyte zonation. Our data suggests a model (Fig 6) in which the Wnt ligand is produced by endothelial cells in the central vein^70^, creating a decreasing gradient of Wnt from the central to the portal vein. High Wnt concentration leads to b-catenin stabilization^71^, Tcf7l1 degradation, and Tcf7l2 activation, resulting in the activation of the pericentral gene program, that includes Tbx3. Tbx3 represses the periportal program (including Tcf7l1) in pericentral hepatocytes. In periportal hepatocytes, with low Wnt concentration, b-catenin is degraded and Tcf7l1 activated. This represses the pericentral program, including Tbx3, and activates the periportal program. Recent work from the Morris lab^72^ has shown that inferred eGRNs can be used to predict the effect of a TF perturbation. We employed a similar strategy within SCENIC+, which revealed that Tbx3 knock-down indeed drives cells to the periportal state, while Tcf7l1 knock-down drives cells to the pericentral state.

**Figure 6.**
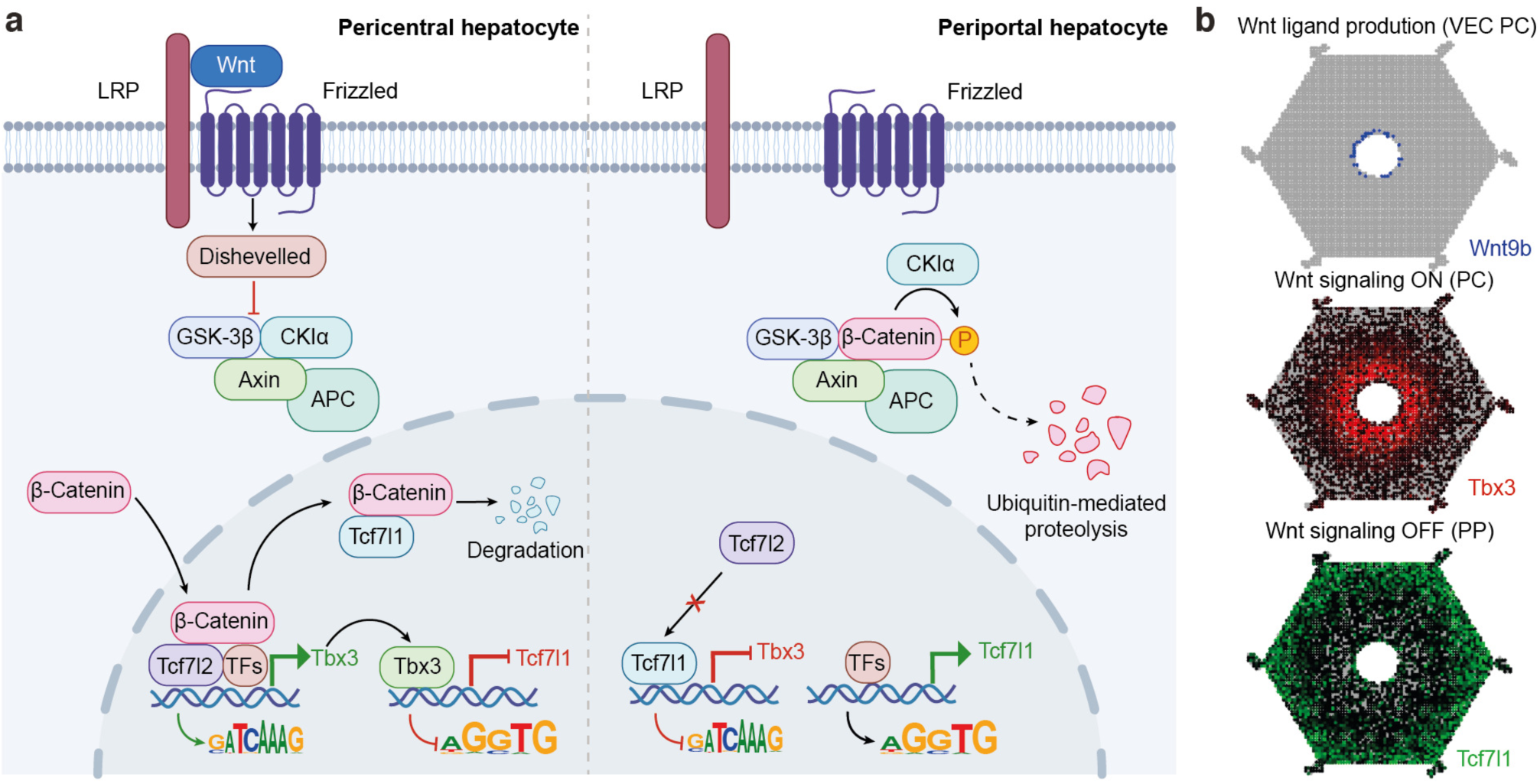
Wnt signaling orchestrates the core GRNs of hepatocyte zonation. **a.** Schematic describing the role of Wnt signaling in hepatocyte zonation. Briefly, Wnt (Wnt9b, Wnt2) is produced by pericentral vascular endothelial cells, creating a decreasing Wnt gradient along the porto-central axis. In pericentral hepatocytes, Wnt signaling is activated resulting in the translocation of b-catenin into the nucleus. b-catenin promotes Tcf7l1 degradation, which allows the binding of general hepatocyte TFs (including Tcf7l2) to the Tcf7l1 repressed regions, opening up chromatin. The binding of these TFs to these regions promotes the transcription of pericentral genes, including Tbx3. Tbx3 will repress its target regions in pericentral hepatocytes, resulting in the silencing of periportal genes. In periportal hepatocytes, reached by lower amounts of Wnt, b-catenin is degraded. Tcf7l1 represses its target regions, resulting in the silencing of pericentral genes, including Tbx3. This allows the binding of hepatocyte TFs to the Tbx3 target regions, opening up chromatin and activating periportal genes. Created with BioRender.com, **b.** ScoMAP liver lobule showing localized gene expression of Wnt9b (Wnt ligand), Tbx3 and Tcf7l1, using RGB encoding.

Topic modelling and spatial transcriptomics suggest the existence of three major zones along the porto-central axis, in agreement with the previously reported metabolic segregation of hepatocytes^73^. Particularly, a pericentral zone is formed by the first 1-2 layers of hepatocytes (with topic 58 and topic 60 accessibility); this is followed by an intermediate-pericentral zone with 6-8 layers of hepatocytes (with topic 60 accessibility) and a periportal zone with 6-10 hepatocyte layers (with topic 66 accessibility). The pericentral topics (topic 58 and topic 60) consist of regions with Tcf7l1/2 motifs while the regions in the periportal topic (topic 66) contain Tbx3 binding sites, suggesting that Tcf7l1 and Tbx3 are directly involved in liver metabolism.

In a second, complementary strategy we trained convolutional neural networks (CNN) to predict, based on the enhancer sequence as input, its ATAC topic membership, or in other words, in which cell type/state the enhancer is accessible. CNN-based enhancer modeling has recently gained traction, due to their capacity for the interpretation of enhancer grammar ^12, 19, 20, 74^ A key limitation of CNN models is that they require large input data sets for training. While training on small data sets may lead to overfitting, transfer learning from sequence models trained with large data sets has been recently shown as a robust alternative^75^, Here, we propose several transfer learning applications, whereby the first (topic-based) model is fine-tuned either to learn cell state (in our case, hepatocyte zonation) or enhancer activity (based on MPRA data). The topic-CNN could recapitulate the core hepatocyte code, with sequence features associated with the same TFs as identified by SCENIC+. The zonation-CNN added Tbx3 and Tcf7l1 motifs as crucial sequence features to the hepatocyte enhancers; while the activity-CNN added ETS and AP-1 sites underlying higher enhancer activity, in agreement with previous MPRA studies in the liver^17^. Importantly, Tbx3 and Tcf7l1 binding sites are located predominantly within hepatocyte enhancers, in close proximity to binding sites of the hepatocyte core TFs. This shows that rather than repressing genes through distinct regulatory regions, these repressor sites form an integral, and likely evolutionary selected, part of the state-specific hepatocyte enhancer logic.

In conclusion, we unraveled the regulatory grammar underlying cellular identity, using hepatocytes as a model cell type. We provide an extensive resource of the adult mouse liver, including a spatial and single-cell multi-omics atlas, eGRNs and enhancer activity, that can be explored in Scope (http://scope.aertslab.org/#/ Bravo_et_al_Liver) and the UCSC genome browser (https://genome.ucsc.edu/s/cbravo/Bravo_et_al_Liver). We envision that our workflow can be used as a roadmap to study other biolgical systems, which will further improve our understanding how cell types and their functional states are encoded in the genome.

## Methods

### Single-cell data generation

#### Mouse liver dissection

All animal experiments were conducted according to the KU Leuven ethical guidelines and approved by the KU Leuven Ethical Committee for Animal Experimentation (approved protocol number ECD P007/2021). Mice were maintained under standard housing conditions, with continuous access to food and water; except for mice 4 and 5, for which food was removed approximately 10 hours before the experiments. Adult mice used in the study were C57BL/6JaxCrl (expect for mouse 3, which was Crl:CD-1). Animals were sacrificed by CO_2_ and the liver was collected for further experiments. For the fresh nuclei isolations, samples were immediately processed. For the frozen nuclei isolation samples were immediately snap frozen in liquid nitrogen and stored at -80°C until processing.

#### Sample and library preparation for 10x snRNA-seq

##### Nuclei isolation

The liver nuclei were isolated following the protocol described by Thrupp et al. (2020)^85^. For the fresh samples, 200 mg fresh big lobe piece of mouse liver tissue was minced and transferred to a Dounce homogenizer cylinder containing 1 mL of ice-cold homogenization buffer (320mM Sucrose, 5mM CaCl_2_, 3mM Magnesium Acetate, 10mM Tris-HCl (ph 7.5), 0.1mM EDTA, 0.1% Igepal, 0.1mM PMSF, 1mM βME and 0.2 U/ul RNasin Plus RNase Inhibitor (Promega). For the frozen samples, a piece of 200 mg liver big lobe was sectioned on dry ice and transferred to a Dounce homogenizer cylinder containing 1 mL of ice-cold homogenization buffer and let to thaw for 5 minutes. From this step onward, both the fresh and frozen tissue were homogenized with 10 strokes of pestle A and 10 strokes of pestle B until a homogeneous nuclei suspension was achieved. The resulting homogenate was filtered through a 70-μm cell strainer (Corning). An addition 1.65 mL of homogenization buffer was topped up and mixed with 2.65 mL of gradient medium (5mM CaCl_2_, 50% Optiprep (Stemcell Technologies), 3mM Magnesium Acetate, 10 mM Tris-HCl (pH 7.5), 0.1mM PMSF, 1mM βME). 4 mL of 29% Iodoxanol cushion was prepared with OptiPrep (Stemcell Technologies) and diluent medium (250 mM sucrose, 150 mM KCl, 30 mM MgCl_2_, 60 mM Tris-HCl (pH 7.5), and added into an ultracentrifuge tube. Next, 5.3 mL of sample in homogenization buffer and gradient medium was gently layered on top of the 29% Iodoxanol cushion. Samples were centrifuged in a SW41Ti rotor (Beckman) at 7700 x g and 4°C for 30 min and the obtained supernatant was gently removed without disturbing the nuclei pellet. Nuclei were resuspended in 200 μL resuspension buffer (1x PBS, 1% BSA and 0.2U μl/1 RNasin Plus RNase Inhibitor (Promega)) and transferred to a 1.5 mL Eppendorf tube tube. 9μL of sample was mixed with 1μL of arginine orange/propidium iodide (AO/PI) stain, loaded onto a LUNA-FL slide and visualized with the LUNA-FL Automated cell counter for nuclei yield, morphology and presence of clumps/debris.

##### Library preparation

Single-nuclei libraries were generated using the 10x Chromium Single-Cell Instrument and Chromium Single Cell 3’ Reagent v3 Kits (10x Genomics) according to the manufacturer’s protocol. In brief, the single mouse liver nuclei suspension was loaded in the Chromium chip for partitioning into nanoliter-scale Gel Beads-in-emulsion (GEMs). After GEMs generation, the obtained emulsion was incubated in a C1000 Touch Thermal Cycler (Bio-Rad) under the following program: 53°C for 45 min, 85°C for 5 min and hold at 4°C. Incubation of the GEMs produced barcoded, full-length cDNA from poly-adenylated mRNA. After incubation, single-cell droplets were dissolved, and full-length cDNA were isolated using Cleanup Mix containing Silane Dynabeads. To generate sufficient mass for library construction, the cDNA was amplified via PCR: 98°C for 3 min; 12 cycles of 98°C for 15 s, 63°C for 20 s, 72°C for 1 min; 72°C for 1 min; and hold at 4°C. Subsequently, the amplified cDNA was fragmented, end-repaired, A-tailed and index adaptor ligated, with SPRIselect cleanup in between steps. The final gene expression library was amplified by PCR: 98°C for 45 s; 10-12 cycles of 98°C for 20 s, 54°C for 30 s, 72°C for 20 s. 72°C for 1 min; and hold at 4°C. The sequencing-ready libraries were cleaned up with SPRIselect beads.

##### Sequencing

Final libraries were quantified using the Qubit dsDNA HS Assay Kit (Life Technologies). The fragment size of every library was analyzed using the Bioanalyzer high-sensitivity chip and were sequenced on HiSeq4000 or NovaSeq6000 instruments with the following sequencing parameters: 28 bp read 1 – 8 bp index 1 (i7) – 0 bp index 2 (i5) – 91 bp read 2.

##### snRNA read mapping

The generated fastq files were processed with cellranger (v1.0.0) count function. Reads were aligned to a pre-mRNA *Mus musculus* reference genome, that listed each gene transcript locus as an exon, and included intronic reads in the counting (10x Genomics, see https:// support.10xgenomics.com/single-cell-gene-expression/software/pipelines/latest/advanced/ references#premrna).

#### Sample and library preparation for 10x scATAC-seq

##### Nuclei isolation

The liver nuclei were isolated using a modified protocol from the Nuclei Isolation for Single Cell ATAC Sequencing (CG000169) Demonstrated Protocol from 10x Genomics. In brief, 200 mg fresh big lobe piece of mouse liver tissue was minced and transferred to a Dounce homogenizer cylinder containing 1 mL of ice-cold homogenization buffer (10 mM NaCl, 10 mM Tris-HCl (pH 7.4), 3 mM MgCl2, 0.1% Tween-20, 0.1% IGEPAL CA-63, 0.01% Digitonin, 1% BSA) and incubated for 5 min on ice. Next the tissue was homogenized with 15 strokes of pestle A and 15 strokes of pestle B until a homogeneous nuclei suspension was achieved. The resulting homogenate was filtered through a 70-μm cell strainer (Corning). The tissue material was spin down at 500 x g for 5 min at 4°C and the supernatant was discarded. The tissue pellet was resuspended in 1 mL wash buffer (20 mM NaCl, 20 mM Tris-HCl (pH 7.4), 6 mM MgCl_2_, 1% BSA). The wash step was repeated one more time and the resulting final pellet was resuspended in 100 µl Diluted Nuclei Buffer (10x Genomics scATAC kit). 9μL of sample was mixed with 1μL of arginine orange/propidium iodide (AO/PI) stain, loaded onto a LUNA-FL slide and visualized with the LUNA-FL Automated cell counter for nuclei yield, morphology, and presence of clumps/debris.

##### Library preparation

Single-nuclei libraries were generated using the 10x Chromium Single-Cell Instrument and Single Cell ATAC v1 kit (10x Genomics) according to the manufacturer’s protocol. In brief, the single mouse liver nuclei were incubated for 60 min at 37°C with a transposase that fragments the DNA in open regions of the chromatin and adds adapter sequences to the ends of the DNA fragments. After generation of nanolitre-scale gel bead-in-emulsions (GEMs), GEMs were incubated in a C1000 Touch Thermal Cycler (Bio-Rad) under the following program: 72°C for 5 min; 98°C for 3 s; 12 cycles of 98°C for 10 s, 59°C for 30 s, 72°C for 1 min; 72°C for 1 min; and hold at 4°C. Incubation of the GEMs produced 10x barcoded DNA from the transposed DNA. Next, single-cell droplets were dissolved, and the transposed DNA was isolated using Cleanup Mix containing Silane Dynabeads. Illumina P7 sequence and a sample index were added to the single-strand DNA during ATAC library construction via PCR: 98°C for 45 s; 9 cycles of 98°C for 20 s, 67°C for 30 s, 72°C for 20 s; 72°C for 1 min; and hold at 4°C. The sequencing-ready ATAC library was cleaned up with SPRIselect beads (Beckman Coulter).

##### Sequencing

Final libraries were quantified using the Qubit dsDNA HS Assay Kit (Life Technologies). The fragment size of every library was analysed using the Bioanalyzer high-sensitivity chip and were sequenced on NextSeq500 instruments (Illumina) with the following sequencing parameters: 70 bp read 1 – 8 bp index 1 (i7) – 16 bp index 2 (i5) - 70 bp read 2.

##### scATAC read mapping

The generated fastq files were processed with cellranger-atac (v1.2.0) count function. Reads were aligned to *Mus musculus* reference genome (refdata-cellranger-atac-mm10-1.2.0).

#### Sample and library preparation for 10x single-cell multiome ATAC and gene expression

##### Sample preparation

For “Multiome-10x_Fresh_Mouse-4” we used a modified protocol from the Nuclei Isolation from Complex Tissues for Single Cell Multiome ATAC + Gene Expression Sequencing Protocol (CG000375) from 10x Genomics. Briefly, 100 mg fresh big lobe piece of mouse liver tissue was minced and transferred to a Dounce homogenizer cylinder containing 1 mL of ice-cold homogenization buffer (10 mM NaCl, 10 mM Tris-HCl (pH 7.4), 3 mM MgCl_2_, 0,1% IGEPAL CA-63, 1 mM DTT, 1 U/µl of Protector RNase inhibitor (Sigma)). The tissue was homogenized with 5 strokes of pestle A and 10 strokes of pestle B until a homogeneous nuclei suspension was achieved. The resulting homogenate was filtered through a 70-μm cell strainer (Corning). The tissue material was spin down at 500 x g for 5 min at 4°C and the supernatant was discarded. The tissue pellet was resuspended in wash buffer (1% BSA in PBS + 1 U/µl of Protector RNase inhibitor (Sigma)). Nuclei were stained with 7AAD (Thermo Fisher Scientific) and viability sorted on a BD FACS Fusion into 5 mL low bind Eppendorf tube containing BSA with RNase inhibitor. The sorted nuclei were spin down at 500 x g for 5 min at 4°C and the supernatant was discarded. Next, the nuclei were permeabilized by resuspending the pellet in 0.1x lysis buffer (10 mM NaCl, 10 mM Tris-HCl (pH 7.4), 3 mM MgCl_2_, 0,1% IGEPAL CA-63, 0.01% Digitonin, 1% BSA, 1 mM DTT, 1 U/µl of Protector RNase inhibitor (Sigma)) and incubated on ice for 2 min. 1 mL Wash Buffer (10 mM NaCl, 10 mM Tris-HCl (pH 7.4), 3 mM MgCl_2_, 0.1% Tween-20, 1% BSA, 1 mM DTT, 1 U/µl of Protector RNase inhibitor (Sigma)) was added. The nuclei were spin down at 500 x g for 5 min at 4°C and the supernatant was discarded. The nuclei pellet was resuspended in diluted nuclei buffer (1x Nuclei buffer Multiome kit (10x Genomics)), 1 mM DTT, 1 U/µl of Protector RNase inhibitor (Sigma)). For the “Multiome-NST_Fresh_Mouse-5” sample nuclei isolation we used a modified protocol from the Slyper et al. (2020)^86^. Briefly, 100 mg fresh big lobe piece of mouse liver tissue was chopped and transferred to a Dounce homogenizer cylinder containing 1 mL of ice-cold homogenization buffer (Salt-tris solution - 146 mM NaCl, 10 mM Tris 7.5, 1 mM CaCl_2_, 21 mM MgCl_2_, 0.2% IGEPAL CA-63, 0.01% BSA, 0.2 U/µl of RNasin Plus RNase Inhibitor (Promega)). The tissue was homogenized with 5 strokes of pestle A and 10 strokes of pestle B until a homogeneous nuclei suspension was achieved. The resulting homogenate was filtered through a 70-μm cell strainer (Corning). The homogenizer and the filter were rinsed with an additional 1 mL of homogenization buffer and 3 mL salt-tris solution buffer (146 mM NaCl, 10 mM Tris 7.5, 1 mM CaCl_2_, 21 mM MgCl_2_). The tissue material was spin down at 500 x g for 5 min at 4°C. The obtained pellet, after supernatant removal, was resuspended in 1.5 mL salt-tris solution buffer supplemented with 0.2 U/ul RNasin Plus RNase Inhibitor (Promega). The tissue material was spin down at 500 x g for 5 min at 4°C. The obtained pellet, after supernatant removal, was resuspended in 1.5 mL wash buffer (1x PBS, 1% BSA and 0.2U μl/1 RNasin Plus RNase Inhibitor (Promega)). The wash step was repeated one more time. The final pellet was resuspended in 500 µl wash buffer, filtered, stained with DAPI (Thermo Fisher Scientific) and viability sorted on a BD FACS Fusion into 5 mL low bind Eppendorf tube containing BSA with RNase inhibitor. The sorted nuclei were spin down at 500 x g for 5 min at 4°C and the supernatant was discarded. Nuclei were resuspended in 50 μL of resuspension buffer The nuclei pellet was resuspend in diluted nuclei buffer (1x Nuclei buffer Multiome kit (10x Genomics)), 1 mM DTT, 1 U/µl RNasin Plus RNase Inhibitor (Promega)). 9μL of sample was mixed with 1μL of arginine orange/propidium iodide (AO/PI) stain, loaded onto a LUNA-FL slide and visualized with the LUNA-FL Automated cell counter for nuclei yield, morphology, and presence of clumps/debris.

##### Library preparation

Single-nuclei libraries were generated using the 10x Chromium Single-Cell Instrument and NextGEM Single Cell Multiome ATAC + Gene Expression kit (10x Genomics) according to the manufacturer’s protocol. In brief, the single mouse liver nuclei were incubated for 60 min at 37°C with a transposase that fragments the DNA in open regions of the chromatin and adds adapter sequences to the ends of the DNA fragments. After generation of nanolitre-scale gel bead-in-emulsions (GEMs), GEMs were incubated in a C1000 Touch Thermal Cycler (Bio-Rad) under the following program: 37°C for 45 min, 25°C for 30 min and hold at 4°C. Incubation of the GEMs produced 10x barcoded DNA from the transposed DNA (for ATAC) and 10x barcoded, full-length cDNA from poly-adenylated mRNA (for GEX). Next quenching reagent (Multiome 10x kit) was used to stop the reaction. After quenching, single-cell droplets were dissolved and the transposed DNA and full-length cDNA were isolated using Cleanup Mix containing Silane Dynabeads. To fill gaps and generate sufficient mass for library construction, the transposed DNA and cDNA were amplified via PCR: 72°C for 5 min; 98°C for 3 min; 7 cycles of 98°C for 20 s, 63°C for 30 s, 72°C for 1 min; 72°C for 1 min; and hold at 4°C. The pre-amplified product was used as input for both ATAC library construction and cDNA amplification for gene expression library construction. Illumina P7 sequence and a sample index were added to the single-strand DNA during ATAC library construction via PCR: 98°C for 45 s; 7-9 cycles of 98°C for 20 s, 67°C for 30 s, 72°C for 20 s; 72°C for 1 min; and hold at 4°C. The sequencing-ready ATAC library was cleaned up with SPRIselect beads (Beckman Coulter). Barcoded, full-length pre-amplified cDNA was further amplified via PCR: 98°C for 3 min; 6-9 cycles of 98°C for 15 s, 63°C for 20 s, 72°C for 1 min; 72°C for 1 min; and hold at 4°C. Subsequently, the amplified cDNA was fragmented, end-repaired, A-tailed and index adaptor ligated, with SPRIselect cleanup in between steps. The final gene expression library was amplified by PCR: 98°C for 45 s; 5-16 cycles of 98°C for 20 s, 54°C for 30 s, 72°C for 20 s. 72°C for 1 min; and hold at 4°C. The sequencing-ready GEX library was cleaned up with SPRIselect beads.

##### Sequencing

Final libraries were quantified using the Qubit dsDNA HS Assay Kit (Life Technologies). The fragment size of every library was analysed using the Bioanalyzer high-sensitivity chip. All 10x Multiome ATAC libraries were sequenced on NovaSeq6000 instruments (Illumina) with the following sequencing parameters: 50 bp read 1 – 8 bp index 1 (i7) – 16 bp index 2 (i5) - 49 bp read 2. All 10x Multiome gene expression libraries were sequenced on NovaSeq6000 instruments with the following sequencing parameters: 28 bp read 1 – 10 bp index 1 (i7) – 10 bp index 2 (i5) – 75 bp read 2.

##### Multiome (scATAC and scRNA) read mapping

The generated fastq files were processed with cellranger-arc (v1.0.0) count function, with include introns =True option. Reads were aligned to *Mus musculus* reference genome (ata-cellranger-arc-mm10-2020-A-2.0.0).

### Single-cell data analysis

#### Transcriptome analysis

10x snRNA-seq and 10x multiome (gene expression) runs were analyzed first independently using VSN-pipelines (v0.27.0)^28^. Briefly, cells with at least 350 genes expressed and a percentage of mitochondrial reads below 10% were kept. Scanpy (v1.8.2)^29^ was run with default parameters, using the number of principal components automatically selected by VSN-Pipelines and using Leiden clustering with resolutions 0.4, 0.6 and 0.8. Hepatocyte clusters with low gene expression and high percentage of mitochondrial reads were removed, as well as doublets called with Scrublet (v0.2.3)^87^. The samples were merged, obtaining 29,798 high-quality cells, and reanalyzed with VSN-Pipelines. To correct for batch effects, we used Harmony on the selected principal components (34), using Leiden clustering with resolution 0.6, resulting in 15 clusters. The VEC and DC subpopulations were identified according to marker genes. This resulted in the identification of 14 cell types.

#### Epigenome analysis

10x scATAC-seq samples were processed with cisTopic (v0.3.0)^79^, using the cells called by cellRanger (5,628 cells) and mm10 SCREEN regions (1,212,823 regions). For topic modelling, we used WarpLDA^88^ with default parameters, using 500 iterations and inferring models with 2, 5, 10 to 30 (by a step of 1), 35, 40, 45 and 50. This resulted in a model with 19 topics. After correcting sample effects with harmony^89^ (v1.0, applied on the scaled topic distributions), we performed Leiden clustering with resolution 0.6, obtaining 11 clusters. Gene activity was calculated by aggregating the probabilities of regions +/-10kb from the TSS (including the gene body). Cluster annotation was done based on motif enrichment, gene activity and label transfer from the annotated transcriptome with Seurat^90^ (v4.0.3, using cisTopic’s gene activity matrix, cca as reduction and the first 10 dimensions). The labelled 10x scATAC-seq and multiome cells (annotated based on the transcriptome labels) and the scATAC-seq fragments were used as input for pycisTopic (v1.0.1.dev75+g3d3b721)^24^. Briefly, we first created pseduobulks per cell type and performed peak calling using MACS2^91^ (v2.2.7.1, with –format BEDPE –keep-dup all –shift 73 –ext_size 146 as parameters, as recommended for single-cell ATAC-seq data). To derive a set of consensus peaks, we used the iterative overlap peak merging procedure describe in Corces et al. (2018)^92^. First, each summit is extended a ‘*peak_half_width*’ (by default, 250bp) in each direction and then we iteratively filter out less significant peaks that overlap with a more significant one. During this procedure peaks are merged and depending on the number of peaks included into them, different processes will happen: 1) 1 peak: The original peak will be kept, 2) 2 peaks: The original peak region with the highest score will be kept and 3) 3 or more peaks: The original region with the most significant score will be taken, and all the original peak regions in this merged peak region that overlap with the significant peak region will be removed. The process is repeated with the next most significant peak (if it was not removed already) until all peaks are processed. This procedure will happen twice, first in each pseudobulk peaks, and after peak score normalization to process all peaks together. This resulted in 486,888 regions. We further filtered the data set based on the scATAC-seq quality as well, keeping cells with at least 1,000 fragments, FRiP > 0.4 and TSS enrichment > 7, resulting in 22,600 high-quality cells. Topic modelling was performed using Mallet (v2.0), using 500 iterations and models with 2 topics and from 5 to 100 by an increase of 5. Additional models between 75 and 85 (by an increase of 1) were added as we observed that the best model should be on that area based on the model selection metrics, and we selected a model with 82 topics. Batch effects between samples were corrected using harmonypy^89^ (v0.0.6) on the scaled topic distributions, and Leiden clustering with resolution 0.6 resulted in 11 clusters, corresponding to 14 cell types based on previous labelling. Drop-out imputation was performed by multiplying the region-topic and topic-cell probabilities. The imputed accessibility matrix was multiplied by 10^6^. Differentially Accessible Regions (DARs) were calculated between all cell populations and specifically within hepatocytes, HSC and LSEC subgroups, using default parameters and topics were binarized using Otsu thresholding. Hepatocyte DARs and shared hepatocyte topics were curated by performing hierarchical clustering on the pseudobulk probabilities, removing a small fraction of lowly and generally accessible regions, and defining non-overlapping groups between the different gradient groups. Gene Onthology analysis was performed using GREAT^93^. We additionally run MACS2 bdgdiff between hepatocytes, LSEC and HSC zonated state using default parameters. The number of shared regions across mice was calculated as the regions in the shared curated topics. PycisTarget (v1.0.1.dev42+gb6707ee) was run using a custom database with the consensus regions, on DARs, binarized topics (with Otsu thresholding), curated DARs and topics and MACS2 bdgdiff, with and without promoters, and using pycisTarget and DEM^24^.

#### Multiome analysis

The gene expression matrix, the imputed accessibility from pycisTopic and the motif enrichment results were used as input for SCENIC+ (v 0.1.dev411+gf4bcae5.d20220810)^24^, using only the multiome cells for eGRN inference. SCENIC+ was run with default parameters, on the complete data set and only using hepatocytes, using http://nov2020.archive.ensembl.org/ as Biomart host. Briefly, a search space of a maximum between either the boundary of the closest gene or 150 kb and a minimum of 1 kb upstream of the TSS or downstream of the end of the gene was considered for calculating region-to-gene relationships using gradient boosting machine regression. TF-to-gene relationships were calculated using gradient boosting machine regression between all TFs and all genes. Final eRegulons were constructed using the GSEA approach in which region-to-gene relationships were binarized based on gradient boosting machine regression importance scores using the 85th, 90th and 95th quantile; the top 5, 10 and 15 regions per gene and using the BASC method for binarization^94^. Regulons between the two runs (with all cells and only hepatocytes) were merged. Gene-based and region-based regulons were scored in the relevant data sets (multiome, all scRNA-seq and scATAC-seq and spatial templates) using AUCell^95^. Regulons with positive region-to-gene relationships, at least 20 target genes and a correlation between gene-based and region-based AUC scores above 0.4 were kept, obtaining 180 high quality regulons.

#### Hi-C and ChIP-seq data analysis

To validate these regulons, we used publicly available Hi-C and ChIP-seq data^50, 51^. Briefly, the Hi-C data was processed using Juicer (v1.9.9), extracting values using KR for normalization by 5kb windows, and keeping only links with score > 10 and involving a bin that overlaps at least one of the consensus peaks and a TSS (+/-1000bp), resulting in 890,488 region-gene links. For the ChIP-seq data processing, reads were mapped to the mm10 genome using Bowtie2 (v2.3.5.1)^96^, peaks were called with MACS2 (v2.2.7.1, with --format BAM --gsize mm --qvalue 0.05 --nomodel --keep-dup all --call- summits –nolambda as options) bigwig files were generated using deepTools^97^ bamCoverage function (v3.5.0, with --normalizeUsing CPM --binSize 1 as parameters). Coverage on the regulon regions was obtained with deepTools computeMatrix.

#### Downstream analyses

Pseudotime order was calculated using the DPT() function of destiny (v3.2.0)^98^ per cell type using as input the harmony corrected PCs and topics from the snRNA-seq and scATAC-seq analyses, respectively. To assess the number of regions and regions affected by zonation, we took the shared regions in hepatocytes, LSEC and HSC (based on topics) and marker genes, respectively, and a Generalized Additive Model (GAM) was fitted based on their accessibility and expression over pseudotime (representing zonation). After filtering for genes fitted with p-adj < 0.01. We identified 220, 275 and 2697 genes and 281, 475 and 8.805 regions that vary along the porto-central axis in HSC, LSEC and hepatocytes, respectively. To rank regulons (or signatures) based on how affected they are by zonation and/or sample, we performed ANOVA over the AUC values along the pseudotime per sample (adjusting the p-values with Bonferroni). We found that PCA dimensionality reduction of the AUC regulon values matrix represented zonation (x-axis) and sample biases (y-axis). This approach was also used to identify pathways affected by zonation and/or sample, derived Halpern et al. (2017)^8^, using the AUC values after scoring the signatures with AUCell (v0.11.2+19.gfaa0216)^95^ on the cells (scRNA-seq or gene activities from scATAC-seq). To obtain the circadian rhythm signatures, we used the single cell RNA-seq data from the mouse liver at different time points of the circadian rythm from Droin et al. (2021)^34^, performing differential expression analysis between the different time points with Seurat (v4.0.03)^90^. Combined genome coverage and gene expression plots were done using Signac^99^.

### Molecular Cartography in the mouse liver

#### Gene panel selection

100 genes were selected based on their gene expression patterns (marker genes for a cell type or group of cell types) on our in-house mouse liver data set and literature (Table S1). In addition, we performed dimensionality reduction only using these 100 genes to ensure that all cell types could be distinguished with this gene panel.

#### Tissue sections

Mouse liver samples were fixed with PAXgene Tissue FIX solution (Resolve Biosciences) for 24 hours at room temperature followed by two hours in PAXgene Tissue Stabilizer (Resolve Biosciences) at room temperature. Samples were cryoprotected in a 30% sucrose solution (w/v) overnight at 4°C and frozen in 2-methylbutane (Sigma-Aldrich 106056) on dry ice. Frozen samples were sectioned with a cryostat (Leica CM3050) and 10µm thick sections were placed within the capture areas of cold Resolve Biosciences slides. Samples were then sent to Resolve BioSciences on dry ice for analysis. Upon arrival, tissue sections were thawed and rehydrated with isopropanol, followed by one min washes in 95% Ethanol and 70% Ethanol at room temperature. The samples were used for Molecular Cartography^TM^ (100-plex combinatorial single molecule fluorescence in-situ hybridization) according to the manufacturer’s instructions (protocol v1.3; available for registered users), starting with the aspiration of ethanol and the addition of buffer DST1 followed by tissue priming and hybridization. Briefly, tissues were primed for 30 minutes at 37°C followed by overnight hybridization of all probes specific for the target genes (see below for probe design details and target list). Samples were washed the next day to remove excess probes and fluorescently tagged in a two-step color development process. Regions of interest were imaged as described below and fluorescent signals removed during decolorization. Color development, imaging and decolorization were repeated for multiple cycles to build a unique combinatorial code for every target gene that was derived from raw images as described below.

#### Probe Design

The probes for the 100 selected genes were designed using Resolve’s proprietary design algorithm. Briefly, the probe-design was performed at the gene-level. For every targeted gene all full-length protein coding transcript sequences from the ENSEMBL database were used as design targets if the isoform had the GENCODE annotation tag ‘basic’^100^. To speed up the process, the calculation of computationally expensive parts, especially the off-target searches, the selection of probe sequences was not performed randomly, but limited to sequences with high success rates. To filter highly repetitive regions, the abundance of k-mers was obtained from the background transcriptome using Jellyfish^101^. Every target sequence was scanned once for all k-mers, and those regions with rare k-mers were preferred as seeds for full probe design. A probe candidate was generated by extending a seed sequence until a certain target stability was reached. A set of simple rules was applied to discard sequences that were found experimentally to cause problems. After these fast screens, every kept probe candidate was mapped to the background transcriptome using ThermonucleotideBLAST^102^ and probes with stable off-target hits were discarded. Specific probes were then scored based on the number of on-target matches (isoforms), which were weighted by their associated APPRIS level^103^, favoring principal isoforms over others. A bonus was added if the binding-site was inside the protein-coding region. From the pool of accepted probes, the final set was composed by greedily picking the highest scoring probes. Gene names and Catalogue numbers for the specific probes designed by Resolve BioSciences are included in Table S1.

#### Imaging

Samples were imaged on a Zeiss Celldiscoverer 7, using the 50x Plan Apochromat water immersion objective with an NA of 1.2 and the 0.5x magnification changer, resulting in a 25x final magnification. Standard CD7 LED excitation light source, filters, and dichroic mirrors were used together with customized emission filters optimized for detecting specific signals. Excitation time per image was 1000 ms for each channel (DAPI was 20 ms). A z-stack was taken at each region with a distance per z-slice according to the Nyquist-Shannon sampling theorem. The custom CD7 CMOS camera (Zeiss Axiocam Mono 712, 3.45 µm pixel size) was used. For each region, a z-stack per fluorescent color (two colors) was imaged per imaging round. A total of 8 imaging rounds were done for each position, resulting in 16 z-stacks per region. The completely automated imaging process per round (including water immersion generation and precise relocation of regions to image in all three dimensions) was realized by a custom python script using the scripting API of the Zeiss ZEN software (Open application development).

#### Spot Segmentation

The algorithms for spot segmentation were written in Java and are based on the ImageJ library functionalities. Only the iterative closest point algorithm is written in C++ based on the libpointmatcher library (https://github.com/ethz-asl/libpointmatcher).

#### Preprocessing

As a first step all images were corrected for background fluorescence. A target value for the allowed number of maxima was determined based upon the area of the slice in µm² multiplied by the factor 0.5. This factor was empirically optimized. The brightest maxima per plane were determined, based upon an empirically optimized threshold. The number and location of the respective maxima was stored. This procedure was done for every image slice independently. Maxima that did not have a neighboring maximum in an adjacent slice (called z-group) were excluded. The resulting maxima list was further filtered in an iterative loop by adjusting the allowed thresholds for (Babs-Bback) and (Bperi-Bback) to reach a feature target value (Babs: absolute brightness, Bback: local background, Bperi: background of periphery within 1 pixel). This feature target values were based upon the volume of the 3D-image. Only maxima still in a zgroup of at least 2 after filtering were passing the filter step. Each z-group was counted as one hit. The members of the z-groups with the highest absolute brightness were used as features and written to a file. They resemble a 3D-point cloud. Final signal segmentation and decoding: To align the raw data images from different imaging rounds, images had to be corrected. To do so the extracted feature point clouds were used to find the transformation matrices. For this purpose, an iterative closest point cloud algorithm was used to minimize the error between two point-clouds. The point clouds of each round were aligned to the point cloud of round one (reference point cloud). The corresponding point clouds were stored for downstream processes. Based upon the transformation matrices the corresponding images were processed by a rigid transformation using trilinear interpolation. The aligned images were used to create a profile for each pixel consisting of 16 values (16 images from two color channels in 8 imaging rounds). The pixel profiles were filtered for variance from zero normalized by total brightness of all pixels in the profile. Matched pixel profiles with the highest score were assigned as an ID to the pixel. Pixels with neighbors having the same ID were grouped. The pixel groups were filtered by group size, number of direct adjacent pixels in group, number of dimensions with size of two pixels. The local 3D-maxima of the groups were determined as potential final transcript locations. Maxima were filtered by number of maxima in the raw data images where a maximum was expected. Remaining maxima were further evaluated by the fit to the corresponding code. The remaining maxima were written to the results file and considered to resemble transcripts of the corresponding gene. The ratio of signals matching to codes used in the experiment and signals matching to codes not used in the experiment were used as estimation for specificity (false positives).

#### Visualization and nuclei segmentation

Final image analysis was performed in ImageJ using the Polylux tool plugin from Resolve BioSciences to examine specific Molecular Cartography^TM^ signals. Nuclei segmentation was performed using QuPATH (v4.2.1)^104^ based on the DAPI signal, setting pixel size to 0.25, minimum area to 10, maximum area to 400, sigma to 1.7 and cell expansion 8. Data was analyzed using Seurat^90^. Using 14 PCs, we performed Leiden clustering, resulting in 19 clusters that corresponded to 11 cell types, that were annotated based on marker gene expression.

#### Single-cell data mapping

The liver lobule representation was used to generate the virtual liver lobule template coordinates. The template was reduced to a size of 100 × 100 pixels and was split into one image per cell type (in red color). Each image was read using the jpeg (v0.1-8) R package, and the background (in white color) was removed using k-means clustering on the RGB pixel values. Cells in the bile duct were labeled as BECs, in the portal vein we included the periportal vascular endothelial cells and fibroblast and we mapped the pericentral vascular endothelial cells around the central vein. The remaining cell types were spread in the lobule randomly based on the proportions of the cell types in the snRNA-seq data. Multiome cells (with gene expression, chromatin accessibility, regulatory topics and eGRNs) were mapped into a template of the liver lobule and the smFISH spatial map using ScoMAP (v0.1.0)^11^. Briefly, zonated cell types (hepatocytes, LSEC and HSC) were first ordered by pseudotime using the DPT() function from the destiny R package (v3.2.0)^98^, using as input the harmony corrected PCs from the snRNA-seq layer. The pseudotime order represents the distance along the porto-central axis. Each cell type was divided into 10 bins based on their pseudotime order. In the liver lobule template, we calculate the distance of each metacell (i.e. pixel) from the central vein, and divide the cells in 10 bins. The zonated cell types were ordered based on pseudotime using PCs calculated by Seurat^90^ (v4.0.3, which represented the distance along the porto-central axis as well) and divided into 10 bins. For each cell type, we assigned a real profile from the matching bin to each virtual cell randomly (e.g., the cells in the first bin of a pseudotime ordered cell type are assigned to the virtual cells in the first bin of that cell type based on the distance to the central vein in the liver lobule template, or to the same bin in the pseudotime order for the smFISH template). For non-spatially located cell types, cells were sampled randomly without binning based on the annotations between the templates and the single-cell data. If there are more real cells than virtual ones, random sampling is done without repetition; if there are more virtual cells than real ones, real profiles are assigned more than once. The gene expression, region accessibility, topic contribution and regulon enrichment values of the virtual cells are those of their matching real cell. These approaches are included in the ScoMAP R package, with detailed tutorials, at https://github.com/aertslab/ScoMAP.

### Massively Parallel Reporter Assays

#### Library design

For the first library (12K library), we selected 10,845 candidate regions based only on accessibility in hepatocytes. This set also includes shared regions (accessible in hepatocytes and at least one other cell type, 4,163, out of which 1,386 are accessible in hepatocytes and BECs); regions specifically accessible across all hepatocytes (4,357); and regions that are only accessible in periportal, intermediate, and pericentral hepatocytes (795, 527 and 656, respectively). In addition, we included 795 shuffled regions as negative control, and 360 positive controls that previously showed activity in HepG2, and for which a mouse orthologous region can be found that is also accessible in the mouse liver. For this latter subset, we included both the human and the corresponding mouse sequences^16–18^. For each region we selected 258 bp sequence centered at the peak summit, to which we added a 12 bp random sequence as barcode selected from https://github.com/hawkjo/freebarcodes, excluding those with repeats (more than the same nucleotide 6 times in a row). The sequences were flanked with the adaptors CCAGTGCAAGTGCAG and GGCCTAACTGGCCGG in 5’ and 3’, respectively, resulting in 300bp sequences. The final library was synthesized by Twist Bioscience as an Oligo Pool. For the second library (455 library), we selected 13 periportal enhancers, 21 pericentral enhancers, 2 positive controls (AldoB and LTV1^14^) and 44 shuffled regions. For each enhancer, we manually introduced mutations affecting activity and/or zonation based on the saturation mutagenesis of DeepLiver. We then selected 259bp windows, based on the information content from DeepExplainer. For enhancers in which the 259bp window did not cover all the relevant nucleotides, we selected more than one window. This resulted in 16 periportal and 25 pericentral windows, with 370 sequence variants in total. For each region, we added an 11 bp random sequence as barcode selected from https://github.com/hawkjo/freebarcodes, excluding those with repeats (more than the same nucleotide 6 times in a row). The sequences were flanked with the adaptors CCAGTGCAAGTGCAG and GGCCTAACTGGCCGG in 5’ and 3’, respectively, resulting in 300bp sequences. The final library was synthesized by Twist Bioscience as an Oligo Pool.

#### Enhancer library cloning

The pSA293-CHEQseq plasmid (Addgene 174669), containing a SCP1 promoter, a chimeric intron and the Venus cDNA, was used as a reporter plasmid for MPRA. Two different versions of that plasmid were used for the cloning of the 12k library: pSA293-CHEQseq-5’BC contains a random 17 bp barcode (BC) upstream of the chimeric intron and pSA293-CHEQseq-3’BC-1 contains a random 17 bp BC between the Venus and the poly(A) tail^56^. The 455 library was cloned in a newly generated pSA293-CHEQseq-3’BC-2 containing an 18 bp BC optimized for ONT sequencing with the following pattern: NNNYRNNNYRNNNYRNNN. The oligonucleotide libraries were resuspended according to the manufacturer’s recommendation and amplified via PCR with the primers ‘CHEQ_liver_For’ and ‘CHEQ_liver_Rev’ (Table S4). To clone the amplified enhancer library upstream of the SCP1 promoter, the vectors were linearized via inverse PCR with primers ‘CHEQ_lin_For’ and ‘CHEQ_lin_Rev’ (Table S4). Amplified libraries and the corresponding linearized vector were combined in an NEBuilder reaction with a vector to insert ratio of 1:5. The NEBuilder reactions were dialyzed against water in a 6 cm Petri dish with a membrane filter MF-Millipore 0.05 µm (Merck, Kenilworth, NJ) for 1 hr. Reactions were recovered from the membrane, and 2.5 µL of the reaction was transformed into 25 µL of Lucigen Endura ElectroCompetent Cells (Biosearch Technologies, Hoddesdon, UK). Before culture for maxiprep, 1:100,000 of the transformed bacteria was plated on an LB-agar dish with carbenicillin to estimate the complexity of the cloned library. A volume of bacteria corresponding to a complexity of 500 BCs per enhancer was put in culture for maxiprep. Maxiprep was performed with the Nucleobond Xtra endotoxin-free maxiprep kit (Macherey-Nagel).

#### Enhancer-barcode assignment

For the liver library cloned in the pSA293-CHEQseq-5’BC plasmid, a PCR amplification (12 cycles) of the enhancer, together with the random BC, was done with the primers ‘Enh_BC_5’_For’ and ‘Enh_BC_5’_Rev’ (Table S4). Illumina sequencing adaptors were added during a second round of PCR with the primers ‘i5_Indexing_For’ and ‘i7_Indexing_Rev’ (Table S4). Before sequencing, the fragment size of every library was analyzed using the Bioanalyzer high-sensitivity chip. All libraries were sequenced on a NextSeq2000 instrument (Illumina) with the following sequencing parameters: 51 bp read 1 – 8 bp index 1 – 8 bp index 2 – 51 bp read 2. Reads were first processed with fastqc (v0.11.8) to assess their quality, and then t r immed using cutadapt (v 1 . 18)^105^ with options - g TGTCCCCAGTGCAAGTGCAG --discard-untrimmed -m 12 -l 12 for read 1 to extract the enhancer barcode and options -g AATTAATTCGGGCCCCGGTCC…GATCGGCGCGCCTGCTCG -j 10 --discard-untrimmed -m 17 -M 17 for read 2 to extract the plasmid barcode. For read 2, we then used seqkit (v0.10.2)^106^, with options seq -r -p, to get the reverse complement sequence. Reads were filtered to keep only those with quality > 30 using fastp (v0.20.0)^107^. This resulted in 8,835,050 enhancer-barcode assignments for the 5’ 12K library, with 78.3% barcodes assigned to a unique enhancer. For the liver libraries cloned in the pSA293-CHEQseq-3’BC plasmids, we performed Nanopore sequencing as follows. A total of 1.5 µg of the library was linearized by digestion with NcoI according to the manufacturer’s protocol. We next processed 200 ng of the cleaned up and linearized plasmid with an Oxford Nanopore Technologies Q20+ ligation sequencing kit early-access SQK-LSK112 following the manufacturer’s protocol (Genomic DNA by Ligation; revision GDE_9141_v112_revC_01Dec2021). A total amount of 10 fmol of the prepared library was loaded onto a MinION R10.4 flow cell and sequenced for at least 72 hr (MinKNOW ≥ v21.11.09). Raw signal fast5 files were re-basecalled using Guppy v6.0.7 using the super-accuracy model (dna_r10.4_e8.1_sup). To process the reads, we first generated a synthetic genome consisting of the plasmid sequence (with 100bp flanks) with all possible enhancers inserted. Reads were mapped using minimap2 (v2.22)^108^ with options -ax map-ont –secondary=no -N 1 -f 500 and a bam file was generated using SAMTools (v1.11)^109^. Next, we used the R package GenomicAlignments (v1.24.0) to extract the sequences overlapping the enhancer sequence and the enhancer barcodes from the reads and calculate the number of mismatches. For the 455 library, we kept only those assignments with no mismatches, as many enhancer sequences differ in few base pairs only, and those assignments in which the plasmid barcode sequence matches with NNNYRNNNYRNNNYRNNN. This resulted in libraries with 723,805 and 71,658 unique enhancer-barcode assignments, with 85.5% and 97.8% barcodes assigned to a unique enhancer, respectively.

#### Culture of HepG2 cells

The HepG2 cell line was purchased from ATCC (HB-8065). HepG2 cells were cultured in Eagle’s Minimum Essential Medium (EMEM, Thermo Fisher Scientific) supplemented with 10% fetal bovine serum (Thermo Fisher Scientific) and 50 µg/mL penicillin/streptomycin (Thermo Fisher Scientific). Cell cultures were kept at 37°C, with 5% CO_2_.

#### Bulk MPRA in vitro

The MPRA libraries were transfected in HepG2 cells by use of the Lipofectamine 3000 reagent (Thermo Fisher Scientific). Briefly, 4 million HepG2 cells were seeded in a 10 cm cell culture dish. The next day, when cells reach 70-90% confluency, a tube A with 500 µL opti-MEM (Thermo Fisher Scientific) and 25 µL Lipofectamine 3000 reagent and a tube B with 500 µL opti-MEM and 15 µL of the liver enhancer library were prepared and incubated for 5 min at RT. Tube B was mixed carefully with tube A and incubated for 15 min at RT. The medium of the HepG2 cells was also changed to opti-MEM medium and finally the mixture was added dropwise to the cells. 48 hours post-transfection, cells were detached from the plate using trypsin (Thermo Fisher Scientific). One-fifth of the cells was used for plasmid DNA extraction (Qiagen). The remaining cells were used for RNA extraction using the innuPREP RNA Mini Kit 2.0 (Analytik Jena), followed by mRNA isolation using the Dynabeads mRNA purification kit (Ambion) and cDNA synthesis using the GoScript RT Kit and oligo dT primer (Promega). To amplify the random 5’ and 3’ BC from the plasmid DNA or cDNA sample, a PCR was performed for 16 cycles with ‘CHEQseq_barcode_5’_For’, ‘CHEQseq_barcode_5’_Rev’ and ‘CHEQseq_barcode_3’_For’, ‘CHEQseq_barcode_3’_Rev’ (Table S4), respectively. To add Illumina sequencing adaptors, all samples were finally amplified by PCR for 6 cycles with the primers ‘i5_Indexing_For’ and ‘i7_Indexing_Rev.

#### Bulk MPRA in vivo

For intrahepatic delivery of the liver MPRA libraries, 8 to 10-week-old mice were secured and hydrodynamically injected with 20 µg of the libraries via the lateral tail vein. All libraries were diluted in sterile filtered 0.9% NaCl, and the total volume was adjusted to 10% (in mL) of the total body weight (in grams). 24 and 48 hours post-injection, for the 12K library and the 455 library, respectively, mice were anesthetized by intraperitoneal injection of sodium pentobarbital (Nembutal, 50 mg/kg) and whole livers were isolated. Liver tissues were homogenized by using M tubes (Miltenyi Biotec) and the GentleMACS Dissociator (Miltenyi Biotec). RNA and plasmid DNA extraction, mRNA purification and cDNA prep, and barcode amplification, were done as described for MPRA *in vitro*.

#### FACS MPRA in vivo

For intrahepatic delivery of the liver MPRA libraries, 8 to 10-week-old mice were secured and hydrodynamically injected with 20 µg of the libraries via the lateral tail vein. All libraries were diluted in sterile filtered 0.9% NaCl, and the total volume was adjusted to 10% (in mL) of the total body weight (in grams). 48 hours post-injection, mice were anesthetized by intraperitoneal injection of sodium pentobarbital (Nembutal, 50 mg/kg). Livers were perfused for 5 minutes with 40 mL of perfusion medium SC-1 (8 g/L Nacl, 400 mg/L KCl, 75.5 mg/L NaH_2_PO_4_, 120.5 mg/L Na_2_HPO_4_, 2,38 g/L HEPES, 350 mg/L NaHCO_3_, 190 mg/L EGTA, 900 mg/L D-(+)-Glucose, 1.2 mL Phenol Red solution) to remove the blood, followed by perfusion with 40 mL of SC-2 medium (8 g/L Nacl, 400 mg/L KCl, 75.5 mg/L NaH_2_PO_4_, 120.5 mg/ L Na_2_HPO_4_, 2,38 g/L HEPES, 350 mg/L NaHCO_3_, 560 mg/L CaCl_2_.2H_2_O, 1.2 mL Phenol Red solution) containing 10 mg of collagenase P (Merck) for 5 minutes. Each lobe was dissected off and minced into small pieces in a beaker containing 39 mL SC-2 supplemented with 1 mL DNase I (Sigma-Aldrich) and 20 mg collagenase P, followed by rotating incubation for 15 min at 37°C. Hepatocytes were centrifuged for 2 min at 50 g, washed with PBS, centrifuged again for 2 min at 50 g, resuspended in 3 ml Hoechst buffer (DMEM + 10% FBS + 10 mM HEPES) and filtered through a 70 µm strainer. The protocol for hepatocyte staining was adapted from Ben-Moshe et al. (2019)^61^. After counting the cells on a LUNA cell counter, the concentration was adjusted to 5 million cells in 1 mL of Hoechst buffer. To determine the ploidy of hepatocytes, DNA was stained with Hoechst (Thermo Fisher Scientific) (15 µg/mL). Reserpine (5 µM) was also supplemented to prevent Hoechst expulsion from the cells. Cells were incubated for 30 min at 37°C. Hepatocytes were centrifuged for 5 min at 1,000 rpm at 4°C and the supernatant was discarded. Cells were resuspended in cold PBS in a concentration of 1 million cells in 100 µL. After spinning down (1,000 rpm for 5 min at 4°C), cells were resuspended in FACS buffer (2 mM EDTA, pH 8, and 0.5% BSA in 1x PBS) at a concentration of 1 million cells in 100 µL. Cells were stained with the following antibodies (BioLegend) at a dilution of 1:300: PE anti-mouse/human CD324 E-cadherin (catalogue no. 147304) and APC anti-mouse CD73 (catalogue no. 127210). FcX blocking solution (BioLegend catalogue no. 101319) was added at a dilution of 1:50. Cells were sorted by FACS-Aria-Fusion (BD Biosciences) using a 100 µm nozzle. FSC-A and SSC-A were used for hepatocytes size selection. Cells containing the library were selected based on GFP. Tetraploid hepatocytes were selected based on Hoechst stain. CD73 and Ecad were used to select hepatocytes bins along the porto-central axis, obtaining 100,000-200,000 cells per bin. RNA extraction, mRNA purification and cDNA prep were done as described for MPRA *in vitro*. To amplify the enhancer barcode on the cDNA small modifications were made. To amplify the random 3’ BC, a PCR with 24 cycles was performed. To add Illumina sequencing adaptors, a PCR with 10 cycles was done.

#### MPRA data analysis

CHEQ-seq barcodes were extracted from the plasmid and cDNA samples (read 2) using cut adapt (v 1 . 1 8) ^105^ w i t h parameters with options - g TTATCATGTCTGCTCGAAGC…GATCGGCGCGCCTGCTCG --discard-untrimmed -m 17 -M 17 for the 12K libraries and g GTATCTTATCATGTCTGCTCGAAGC…GATCGGC -j 10 --discard-untrimmed -m 18 -M 18 for the 455 library and seqkit (v0.10.2)^106^, with options seq -r -p, was used to get the reverse complement sequence. Reads were filtered to keep only those with quality > 30 using fastp (v0.20.0)^107^. Reads were assigned to enhancers based on the corresponding enhancer-barcode assignments, resulting in a count matrix with number of reads per enhancer and sample. Samples were processed using DESeq2 (v1.37.6)^110^, comparing the corresponding cDNA replicates versus their plasmid samples. For the FACS fractions, since we did not extract plasmid DNA from the samples, we used the plasmid replicates from the *in vivo* bulk experiment. To assess enhancer activity, we used the LogFC calculated by DESeq2 (v1.37.6)^110^. To distinguish active and inactive enhancers, a Gaussian fit of the shuffled negative control values was performed with robustbase (v0.93-6), and a p-value and Benjamini–Hochberg adjusted p-value was calculated based on that Gaussian fit for all enhancers. An enhancer is considered active if its adjusted p-value is < 0.1.

#### FACS ATAC-seq

Hepatocytes were isolated and stained as described in the FACS MPRA section, with minor modifications. Cells were additionally stained with Alexa Fluor 488 Zombie Green (BioLegend) to enable the detection of viable cells by FACS. Zombie Green was added at a dilution of 1:500 and cells were kept in a rotator in the dark at room temperature for 15 min. Cells were sorted by FACS-Aria-Fusion (BD Biosciences) using a 100 µm nozzle. FSC-A and SSC-A were used for hepatocytes size selection. Viable cells were selected based on the Zombie Green signal. Tetraploid hepatocytes were selected based on Hoechst stain. CD73 and Ecad were used to select hepatocytes bins along the porto-central axis, obtaining 20,000-50,000 cells per bin. Omni-assay for transposase-accessible chromatin using sequencing (OmniATAC-seq) was performed as described previously^111^. Briefly, 20,000-50,000 cells obtained after FACS were resuspended in 50 µL of cold ATAC-seq resuspension buffer (RSB; 10 mM TrisHCl pH 7.4, 10 mM NaCl, and 3 mM MgCl_2_ in water) containing 0.1% NP40, 0.1% Tween-20, and 0.01% digitonin by pipetting up and down three times. This cell lysis reaction was incubated on ice for 3 min. After lysis, 1 mL of ATAC-seq RSB containing 0.1% Tween-20 was added, and the tubes were inverted to mix. Nuclei were then centrifuged for 10 min at 500 g in a pre-chilled (4°C) fixed-angle centrifuge. Supernatant was removed and nuclei were resuspended in 50 µL of transposition mix (25 µL 2x TD buffer, 2.5 µL transposase (Nextera Tn5 transposase, Illumina), 16.5 µL PBS, 0.5 µL 1% digitonin, 0.5 µL 10% Tween-20, and 5 µL water) by pipetting up and down six times. Transposition reactions were incubated at 37°C for 30 min in a thermoblock. Reactions were cleaned-up by MinElute (Qiagen). Transposed DNA was amplified with primers ’i5_Indexing_For’ and ‘i7_Indexing_Rev’ (Table S4). Number of PCR cycles was based on quantitative real-time PCR as described previously^112^. All libraries were sequenced on a NextSeq2000 instrument (Illumina) with the following sequencing parameters: 51 bp read 1 – 8 bp index 1 – 8 bp index 2 – 51 bp read 2. Adapters we removed with fastq-mcf (ea-utils v1.12) and cleaned reads were mapped to the mm10 genome using Bowtie2 (v2.3.5.1)^96^. A fragment count matrix was generated using the liver scATAC-seq consensus peaks using SubRead (v1.6.3)^113^. AUCell^95^ was used to assess the enrichment of the ‘core’ signatures (general, periportal, pericentral-intermediate and pericentral) in each of the fractions, using default parameters.

### DeepLiver

#### Model training

The top 3,000 regions in each topic (based on the region-topic distributions) were used as input for a DL model, where 500 bp DNA sequences were used to predict the topic set to which the region belongs (Topic-CNN). The model is a hybrid CNN-RNN multiclass classifier and its architecture was adopted from earlier studies^20, 25, 74^, trained with Tensorflow (v1.15) with minor adaptations. Briefly, we used 1,024 filters and a filter size of 24. To initialize the filters, we used 725 PWMs derived from running Differentially Enriched Motifs (DEM) between selected cell-type specific topics (topics 48 (hepatocytes), 66 (periportal hepatocytes), 58 (pericentral hepatocytes), 38 (Kupffer cells), 71 (LSEC), 32 (HSC), 27 (Fibroblast) and 42 (BEC)), with Log2FC > 1.5 and adjusted p-value < 0.0001 as thresholds. The Zonation-CNN was trained using regions derived from the curated shared hepatocytes topics and DARs (classified as general, pericentral and periportal after hierarchical clustering, resulting in 12,122, 4,181 and 1,372 regions, respectively), while the MPRA-CNN was trained using the binarized LogFC distributions from the 12K *in vivo* MPRA (with adjusted p-value < 0.01, resulting in 1,232 and 2,983 high-confidence active and inactive regions, respectively). The weights derived from the Topic-CNN model were used to initialize these two models, an approach known as transfer learning. The Zonation-CNN and the MPRA-CNN, which have the same architecture as the Topic-CNN, were trained with identical parameters, except for the learning rate, which was set to 0.00001 instead of 0.001.

#### Model performance

To assess the performance of the models, we performed 9-fold cross validation. Briefly, the data was divided in 10 groups, and in each iteration we used 8 groups for training (80% of the regions), 1 group as validation set (10% of the regions) and 1 group as test set (10% of the regions). To increase the sample size for the DL model, we augmented the regions by extending them to 700 bp and used a sliding window of 500 bp with a 50 bp stride, increasing the sample size five times. During the training, the validation set was used for early stopping and the 12^th^, 66^th^ and 122^nd^ epochs were chosen to evaluate the performance of cross-validation models for the Topic-CNN, the Zonation-CNN and the MPRA-CNN, respectively. After training, we assessed the performance of the models on the non-augmented test set by scoring the test set regions with the models. Then, using the prediction scores and the labels (topics, zonation class or activity pattern for the Topic-CNN. Zonation-CNN and MPRA-CNN, respectively), we calculated the area under the precision-recall (AUPR) and receiver operating characteristic (AUROC) curves using the average_precision_score and roc_auc_score functions from the scikit-learn package (v0.21.3). To validate DeepLiver predictions, we calculated the correlation between the predictions of DeepLiver and a previously published MPRA data set performed on synthetic sequences *in vivo*^57^. These sequences were designed by adding different number of instances and combinations of motifs corresponding to TFs that are relevant to hepatocytes, including Hnf1a, Hnf4a (COUPTF), Cebpa, Onecut1 (HNF6) and Foxa1 (HNF3), among others.

#### Nucleotide contributions

To find the nucleotides that contribute the most to the topic prediction, we used a DeepExplainer, included in the the SHAP package (v0.37.0)^58^, using default parameters and 500 random sequences for initialization. The importance score obtained from the DeepExplainer analysis was multiplied by the one-hot encoded DNA sequence and visualized as the height of the nucleotide letters as in earlier work^114^. In addition to the nucleotide importance plots, we performed *in silico* saturation mutagenesis in which we calculated the effect of each variant of a region on its model prediction score. The sequences with all possible single mutations were generated and the delta prediction score for each topic was calculated. To validate DeepLiver *in silico* mutagenesis, we calculated the correlation between the effect of the mutations predicted by DeepLiver and experimental saturation mutagenesis data on six enhancers from earlier studies (three enhancers from *in vivo* studies and three enhancers from HepG2)^14, 60^.

#### TF-binding site predictions

High nucleotide importances on DeepExplainer plots represent potential binding sites for TFs. We used TF-Modisco (v0.5.14.1)^59^ to identify the most common patterns along the zonation classes (general, pericentral and periportal, using the Zonation-CNN) and active versus inactive enhancers (using the MPRA-CNN). To run TF-Modisco, we used MEME initialization, a sliding window of 15bp, 10 as flank size, and a FDR threshold of 0.15. We then used TF-Modisco patterns and selected PWMs to score the sequences. Briefly, we trimmed them using trim_by_ic=0.25, and the sum score was calculated by using compute_sum_scores on the nucleotide importance scores. We converted the patterns and PWMs to convolutional filters and calculated pattern activation scores using tf.nn.conv1d function from TensorFlow (v1.15). Global motif instances were calculated for the curated shared hepatocyte regions (general, pericentral and periportal). Optimal thresholds were selected manually. To validate the predicted binding hits, we used the ChIP-seq data for Hnf4a, Onecut1, Cebpa and Foxa1^51^, as described for the SCENIC+ regulons.

### Tbx3 and Tcf7l1 perturbation simulation

T b x 3 a n d Tc f 7 l 1 c o m p u t a t i o n a l p e r t u r b a t i o n s w e r e d o n e u s i n g S C E N I C + (v0.1.dev411+gf4bcae5.d20220810)^24^. Briefly, based on the inferred eGRN, we first trained a GBM model per gene, in which we predict the gene expression using its predicted regulators (i.e. TFs). To simulate knockdowns, we set the expression of the selected TF to 0 across all cells and recalculated the predicted gene expression matrix using the previously trained models. To simulate ovexpression, we set the expression of the TFs to the maximum expression value in the data set on hepatocyte cells. Predictions are updated over several iterations, to account for downstream effects.

### Luciferase reporter assay

Synthetic liver sequences were ordered as gBlocks from IDT (Integrated DNA Technologies). The pGL4.23-GW luciferase reporter vector (Promega) was linearized via inverse PCR with primers ‘LUC_lin_For’ and ‘LUC_lin_Rev’ (Table S4). The synthetic sequences and the linearized vector were combined in an NEBuilder reaction and 2 µL of the reaction was transformed into 25 µL of Stellar chemically competent bacteria. HepG2 cells were seeded in 24-well plates at a density of 100,000 cells/well and transfected with 400 ng pGL4.23-enhancer vector + 40 ng pRL-TK renilla vector (Promega) with Lipofectamine 3000 reagent. One day after transfection, luciferase activity was measured via the Dual-Luciferase Reporter Assay System (Promega) by following the manufacturer’s protocol. Briefly, cells were lysed with 100 µL of Passive Lysis Buffer for 15 min at 500 rpm. 20 µL of the lysate was transferred in duplicate in a well of an OptiPlate-96 HB (PerkinElmer) and 100 µL of Luciferase Assay Reagent II was added in each well. Luciferase-generated luminescence was measured on a Victor X luminometer (PerkinElmer). 100 µL of the Stop & Glo Reagent was added to each well, and the luminescence was measured again to record renilla activity. Luciferase activity was estimated by calculating the ratio luciferase/renilla. This value was normalized by the ratio calculated on blank wells containing only reagents. Three biological replicates were done per condition.

### Human liver data analysis

Human liver data was obtained from Zhang et al (2021)^62^. The authors labels and the scATAC-seq fragments were used as input for pycisTopic (v1.0.1.dev75+g3d3b721)^24^. Briefly, we first inferred consensus peaks as previously described, resulting in a data set with 121,593 regions and 6,366 cells. Topic modelling was performed using Mallet (v2.0), using 500 iterations and models with 2 topics and from 5 to 100 by an increase of 5. Drop-out imputation was performend by multiplying the region-topic and topic-cell probabilities. The imputed accessibility matrix was multiplied by 10^6^. The mouse region-based regulons were transformed to hg38 coordinates using liftOver (https://genome.ucsc.edu/cgi-bin/hgLiftOver). The imputed accessibility matrix and the liftovered signatures were used as input for AUCell to assess regulon enrichment.

## Data availability

Data generated in this manuscript (single cell RNA-seq, single cell ATAC-seq, single cell multiome, MPRAs and bulk FACS ATAC-seq) is available at GEO (GEO218472). Signatures for Ras signalling, Wnt signalling pituitary response and hypoxia were obtained from Halpern *et al.* (2017)^8^. Single cell RNA-seq data of the mouse liver at different time points of the circadian rhythm^34^ was downloaded from GEO (GSE145197). ChIP-seq data for Hnf4a, Cebpa, Foxa1 and Onecut1^51^ was downloaded from ENA (PRJEB1571). Hi-C data^50^ was obtained from GEO (GSE65126). Data for MPRA positive controls was retrieved from ENCODE (ENCFF288HIT, ENCFF032RDN)^18^, GEO (GSE71279)^16^ and Klein et al. (2018)^17^. Saturation mutagenesis data was downloaded from https://mpra.gs.washington.edu/satMutMPRA/ and Patwardhan *et al.* (2012)^14, 60^. Human liver scATAC-seq data^62^ was downloaded from GEO (GSE184462).

## Code availability

VSN-Pipelines as available at https://vsn-pipelines.readthedocs.io/. pycisTopic is available at https:// pycistopic.readthedocs.io/. pycistarget is available at https://pycistarget.readthedocs.io/. SCENIC+ is available at https://scenicplus.readthedocs.io/. ScoMAP is available at https://github.com/aertslab/ScoMAP. Notebooks to reproduce the main figures will be available at https://github.com/aertslab/Bravo_et_al_Liver.

## Acknowledgements

Computing was performed at the Vlaams Supercomputer Center (VSC). This work is funded by the following grants to S.Ae: ERC Consolidator Grant (724226_cis-CONTROL), ERC Proof of Concept (963884), Special Research Fund (BOF) KU Leuven (grant C14/18/092), Foundation Against Cancer (2020-1396), and FWO (grants G0B5619N and G094121N) and a PhD fellowship from the FWO to C.B.G.-B. (11F1519N). The authors also thank members of various groups that make curated position weight matrices publicly available, including T. Hughes (cis-bp), M. Bulyk (Uniprobe), A. Mathelier (Jaspar), V. Makeev (Hocomoco) and many others. We thank Resolve Biosciences, specially Jeroen Aerts, for performing the Molecular Cartography experiments in the mouse liver; Janssen Pharmaceutica, VIB Tech Watch and the VIB single-cell accelerator for their help and funding for generating the mouse liver single-cell data; and the VIB FACS expertise center for their assistance during the FACS-MPRA and FACS-ATAC experiments. Figure 6a was created with BioRender.com (Agreement number: BV24NAUKOX).

## Author contributions

C.B.G.-B., S.A and G.Ha. conceived the study. C.B.G.-B. performed the computational analyses, with assistance of I.I.T. and G.Hu. I.M., V.C., L.S.-G., E.V. and S.P. performed the single cell experiments. H.H., R.V., V.C., L.S.-G., J.D., N.P. and D.M. performed the MPRA experiments. V.C. performed the luciferase experiments. C.B.G.-B. and S.A. wrote the manuscript.

---

**Figure S1.**
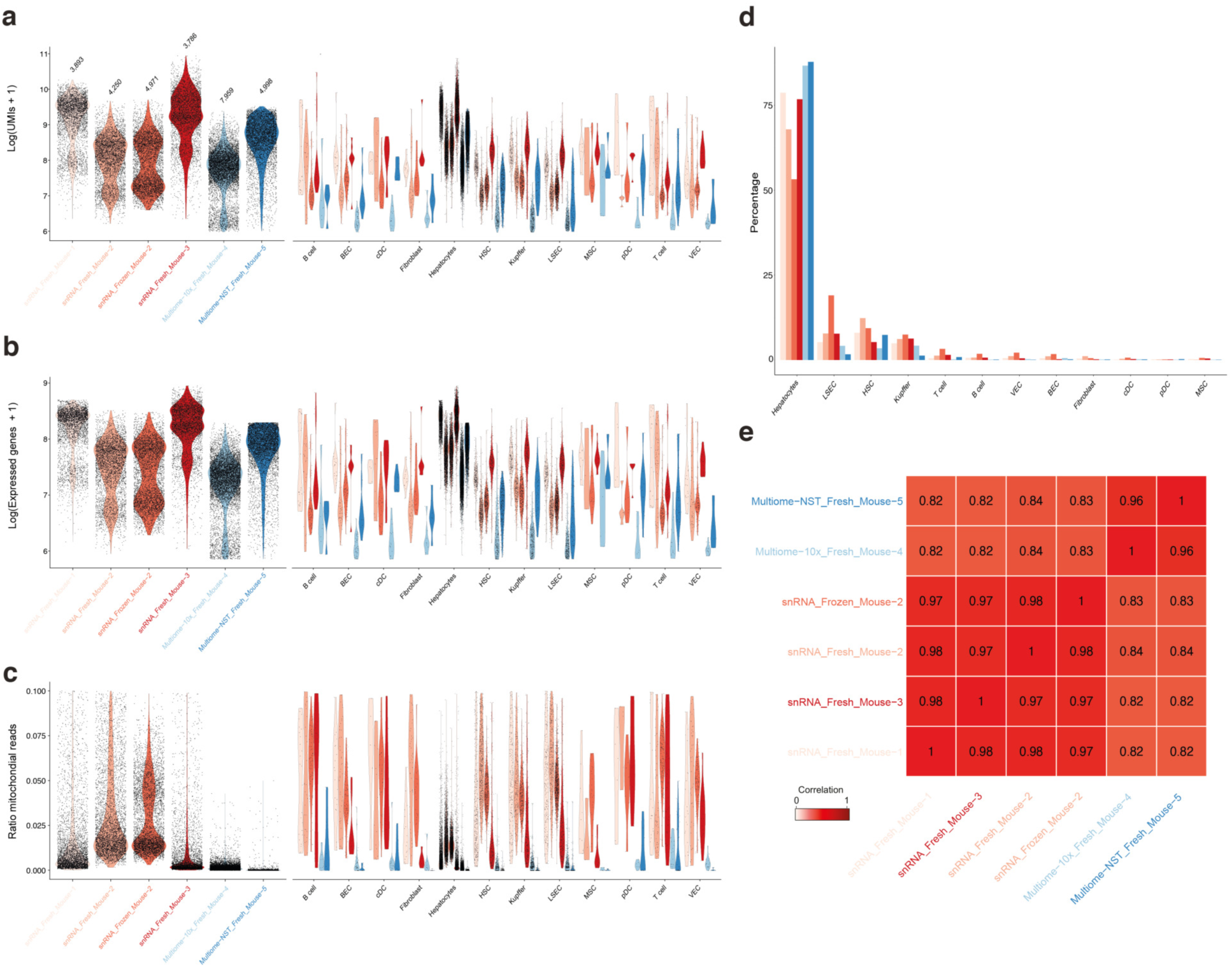
Single nuclei transcriptome quality control. **a.** Violin plot showing the log number of UMIs per sample and cell type. **b.** Violin plot showing the log number of expressed genes per sample and cell type. **c.** Violin plot showing the ratio of mitochondrial reads per sample and cell type. **d.** Barplot showing the percentage of cells corresponding to each cell type across samples. **e.** Correlation between normalized gene expression values (as bulk) across samples.

**Figure S2.**
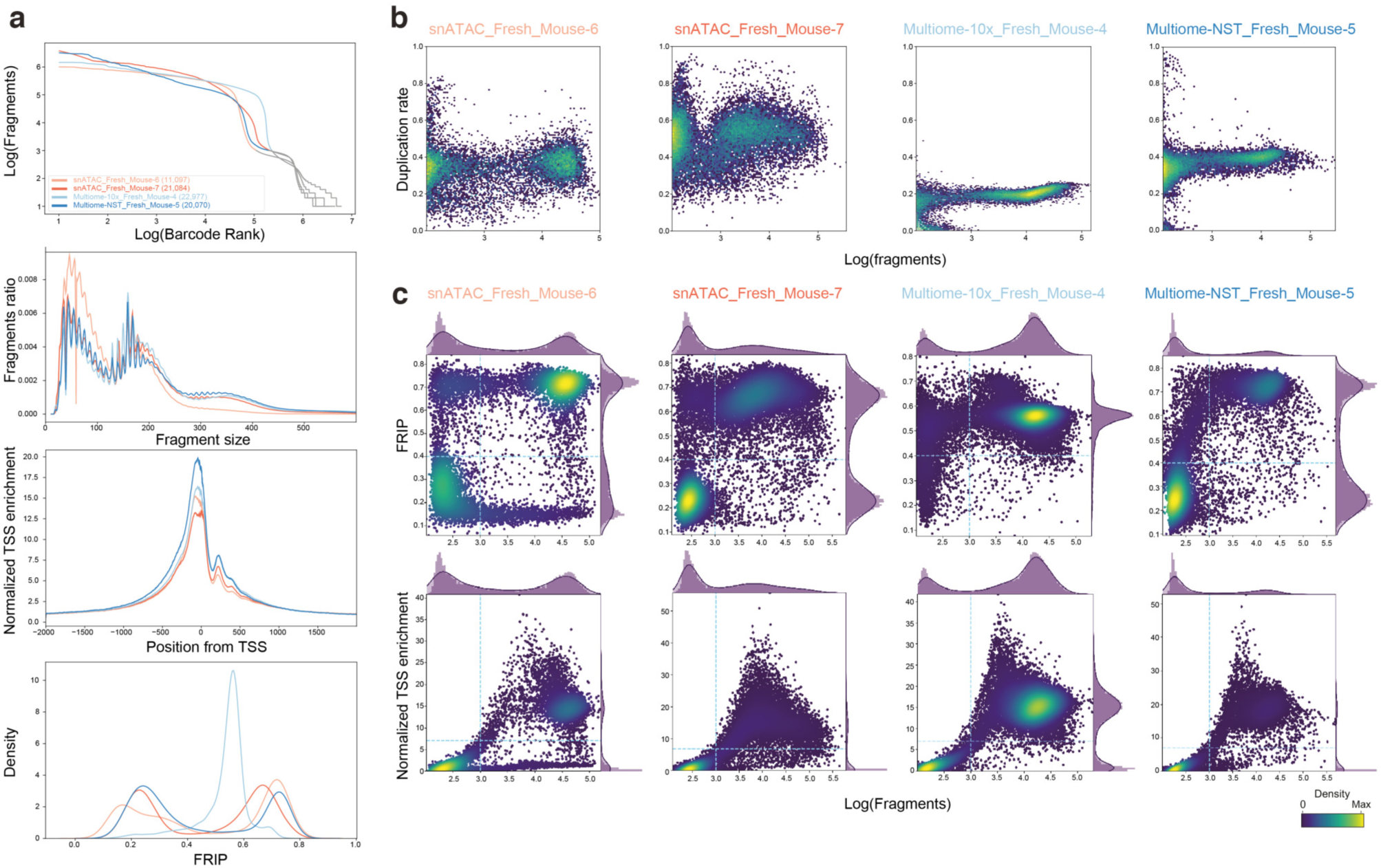
Single nuclei epigenome quality control. **a.** Sample-level epigenome quality control, including (in order top to bottom): barcode rank plot, insert size distribution, TSS enrichment and Fraction of Reads In Peaks (FRIP). **b.** Duplication rate per barcode versus log number of fragments per sample. **c.** Fraction of Reads in Peaks (FRIP, top) and Normalized TSS enrichment (bottom) per barcode per sample. The blue dotted lines indicate the minimum threshold in FRIP, number of fragments and TSS enrichment to select high quality cells.

**Figure S3.**
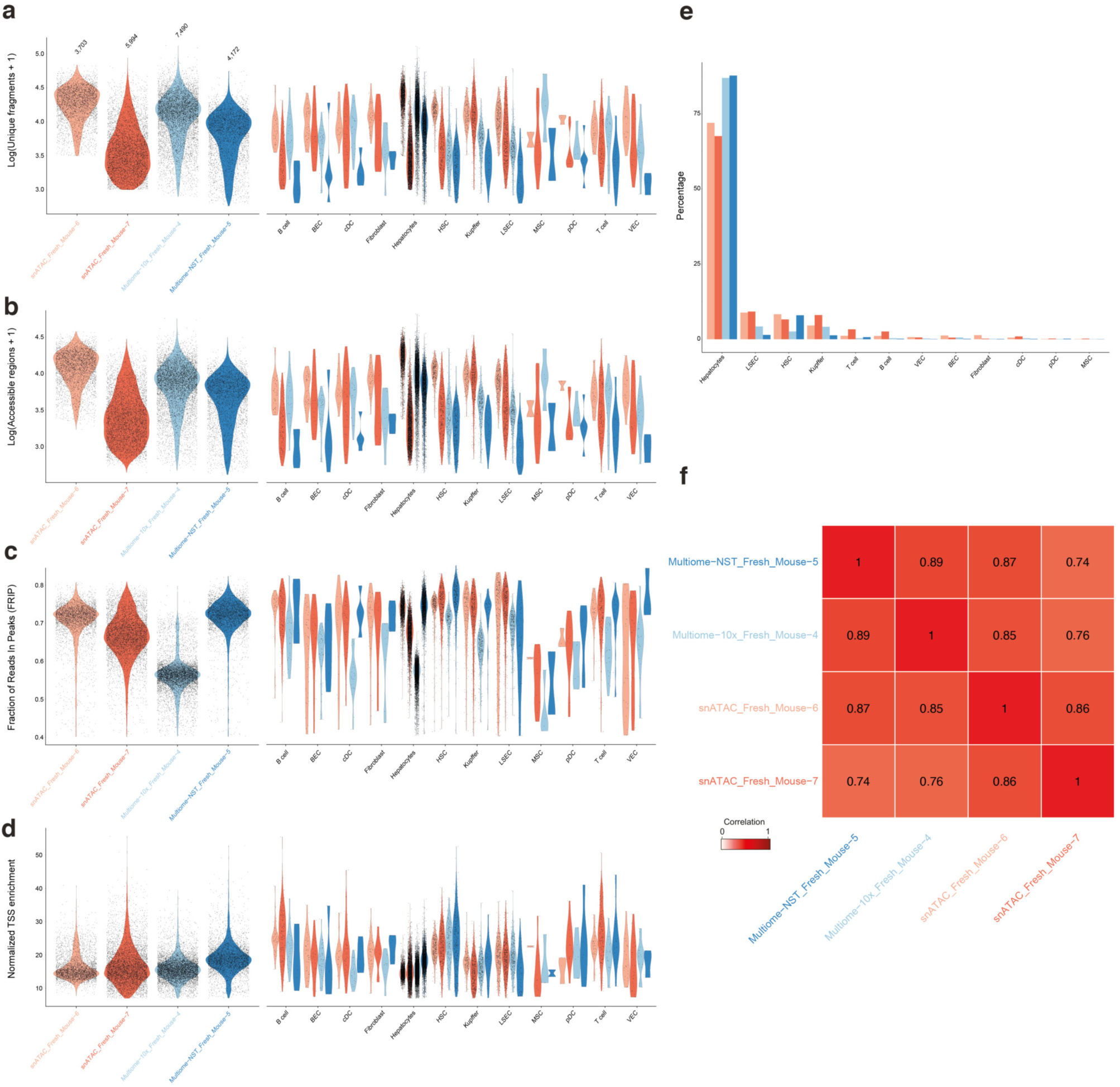
scATAC-seq coverage and quality metrics across samples. **a.** Violin plot showing the log number of fragments per sample and cell type. **b.** Violin plot showing the log number of accessible regions per sample and cell type. **c.** Violin plot showing the Fraction of Reads in Peaks (FRIP) cells per sample and cell type. **d**. Violin plot showing the normalized TSS enrichment across high quality cells per sample and cell type. **e.** Barplot showing the percentage of cells corresponding to each cell type across samples. **f.** Correlation between normalized region accessibility values (as bulk) across samples.

**Figure S4.**
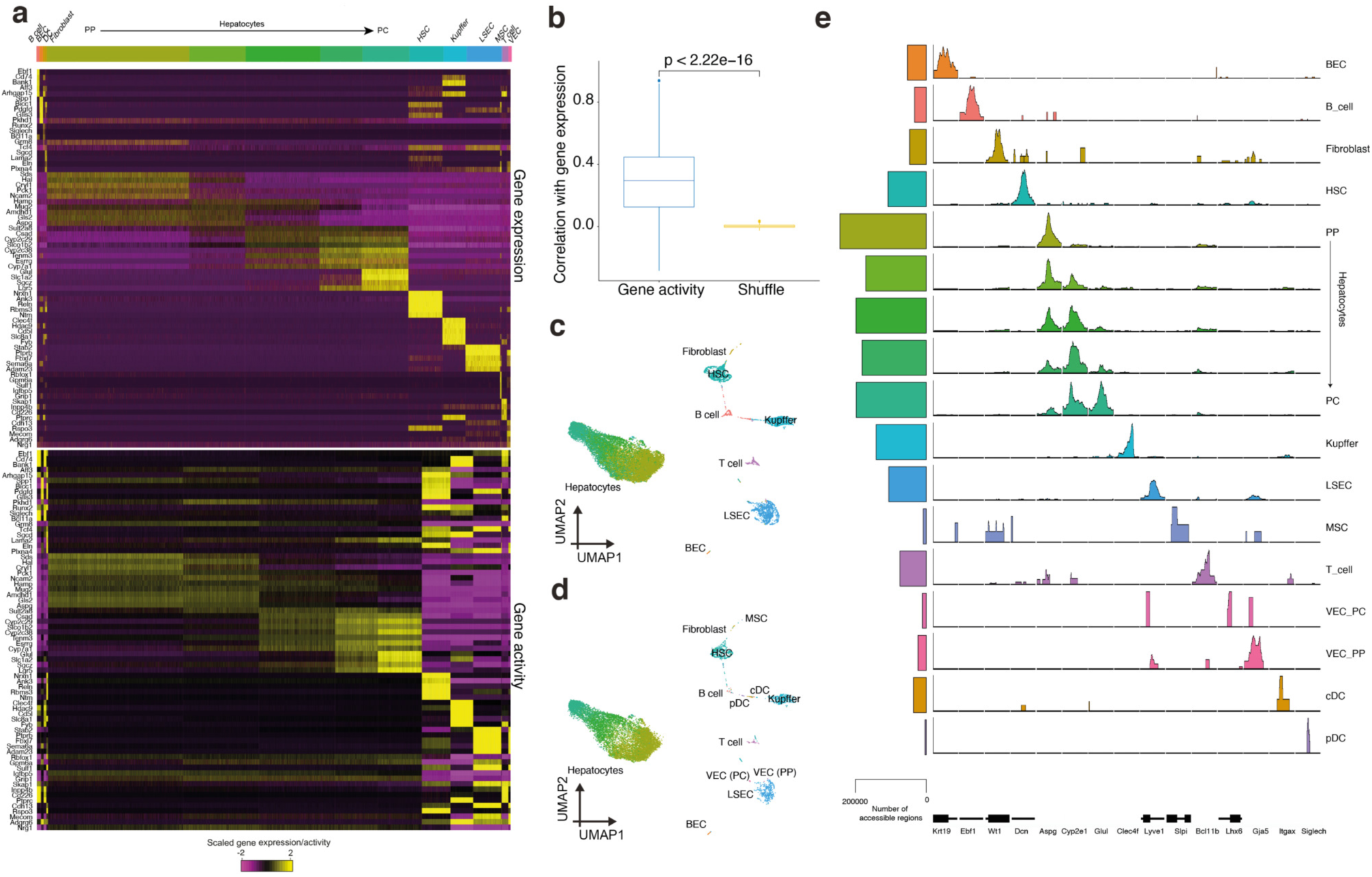
Validation of scATAC-seq topic model with gene activity and label transfer. **a.** Scaled gene expression (top) and scaled gene activity inferred from the scATAC-seq layer (bottom). **b.** Correlation between gene activity and gene expression versus a random control (shuffled gene activity). **c.** scATAC-seq UMAP (22,600 cells) colored by the label transfer cell type annotation (from scRNA-seq cells to scATAC-seq cells). **d.** scATAC-seq UMAP (12,898 cells) colored by the label given to the cell based on the scRNA-seq clustering. **e.** Pseudobulk accessibility profiles on representative Differentially Accessible Regions per cell type. The barplot indicates the number of peaks called by MACS in each pseudobulk.

**Figure S5.**
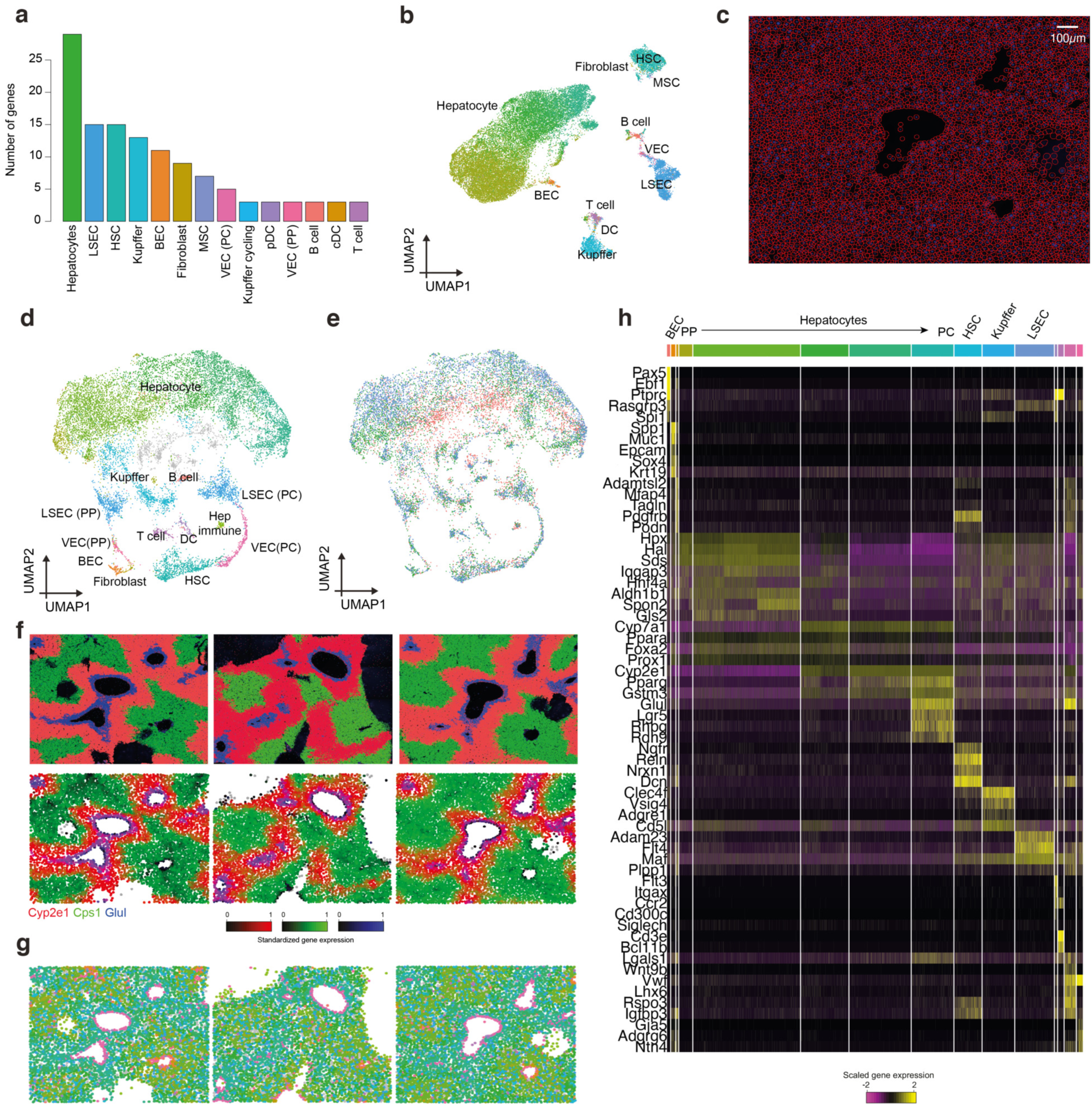
Molecular Cartography of the mouse liver. **a.** Number of selected genes per cell type within the gene panel (100 genes). **b.** scRNA-seq (29,798 cells) UMAP based on the selected 100 genes. **c.** Cell segmentation using the DAPI signal on the sample with QuPath. **d.** UMAP (15,522 spots/cells) of the segmented nuclei based on the number of transcripts measured per gene per spot colored by their assigned cell type. **e.** UMAP (15,522 spots/cells) of the segmented nuclei based on the number of transcripts measured per spot colored by their sample of origin. **f.** Molecular Cartography maps (3 replicates, 15,522 spots) colored by aggregated gene expression using RGB encoding. **g.** Molecular Cartography maps (3 replicates, 15,522 spots) colored by assigned cell type. **h.** Heatmap showing the scaled gene expression across cells (grouped by cell type) of selected genes.

**Figure S6.**
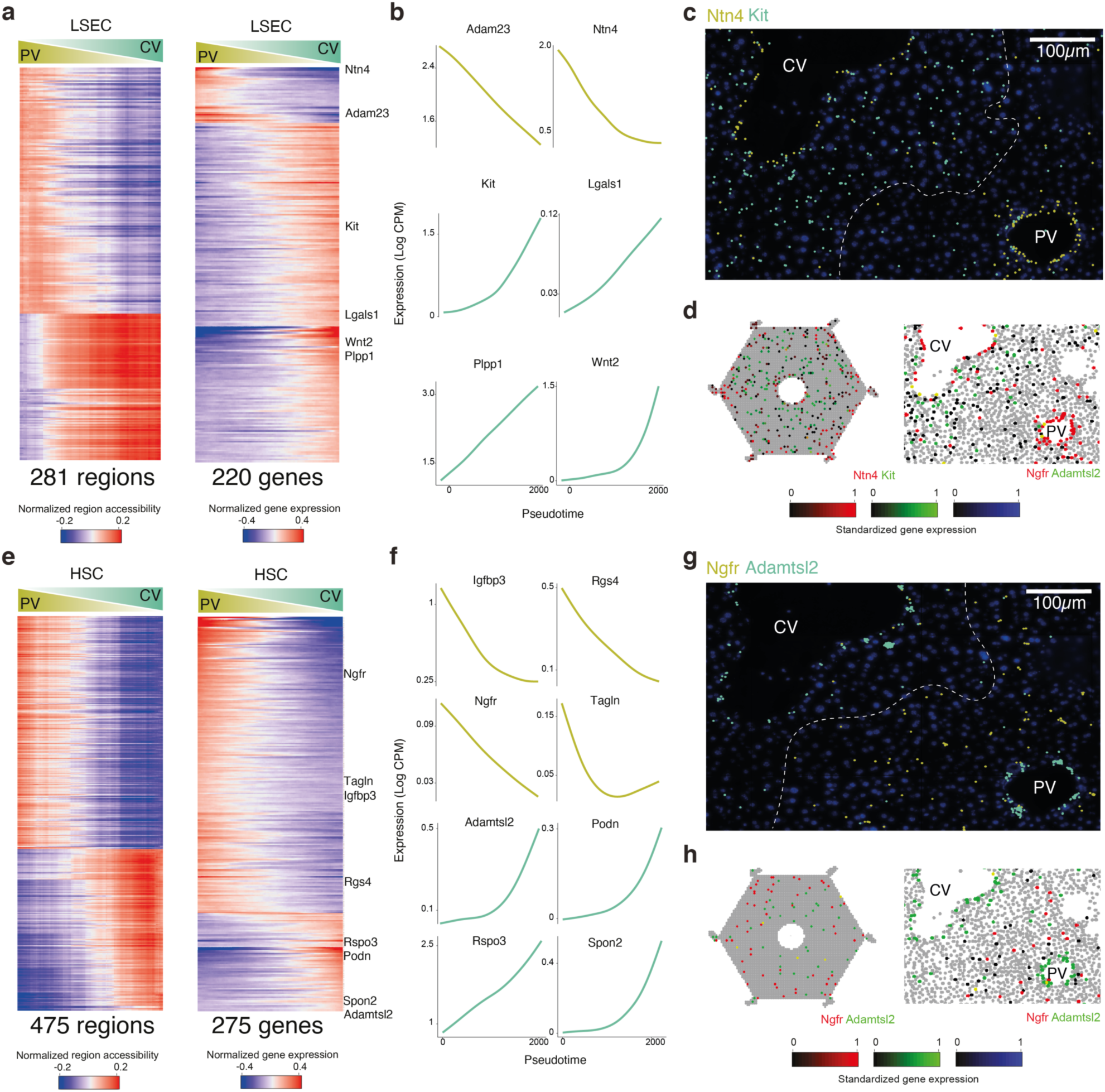
Zonation of Liver Sinusoidal Endothelial Cells (LSEC) and Hepatocellular Stellate Cells (HSC). **a.** Normalized region accessibility and gene expression zonation heatmaps. LSECs are ordered by pseudotime (from periportal to pericentral) and regions and genes affected by zonation are shown (281 regions and 220 genes). **b.** GAM fitted gene expression profiles for selected genes along the zonation pseudotime for LSECs. **c.** Liver section image showing smFISH profiles for Ntn4 (PP LSEC marker) and Kit (PC LSEC marker). **d.** ScoMAP liver lobule and smFISH colored by gene expression using RGB encoding. **e.** Normalized region accessibility and gene expression zonation heatmaps. HSCs are ordered by pseudotime (from periportal to pericentral) and regions and genes affected by zonation are shown (475 regions and 275 genes). **f.** GAM fitted gene expression profiles for selected genes along the zonation pseudotime for HSCs. **g.** Liver section image showing smFISH profiles for Ntgr (HSC PP marker) and Adamtsl2 (PC HSC marker). **d.** ScoMAP liver lobule and smFISH colored by gene expression using RGB encoding.

**Figure S7.**
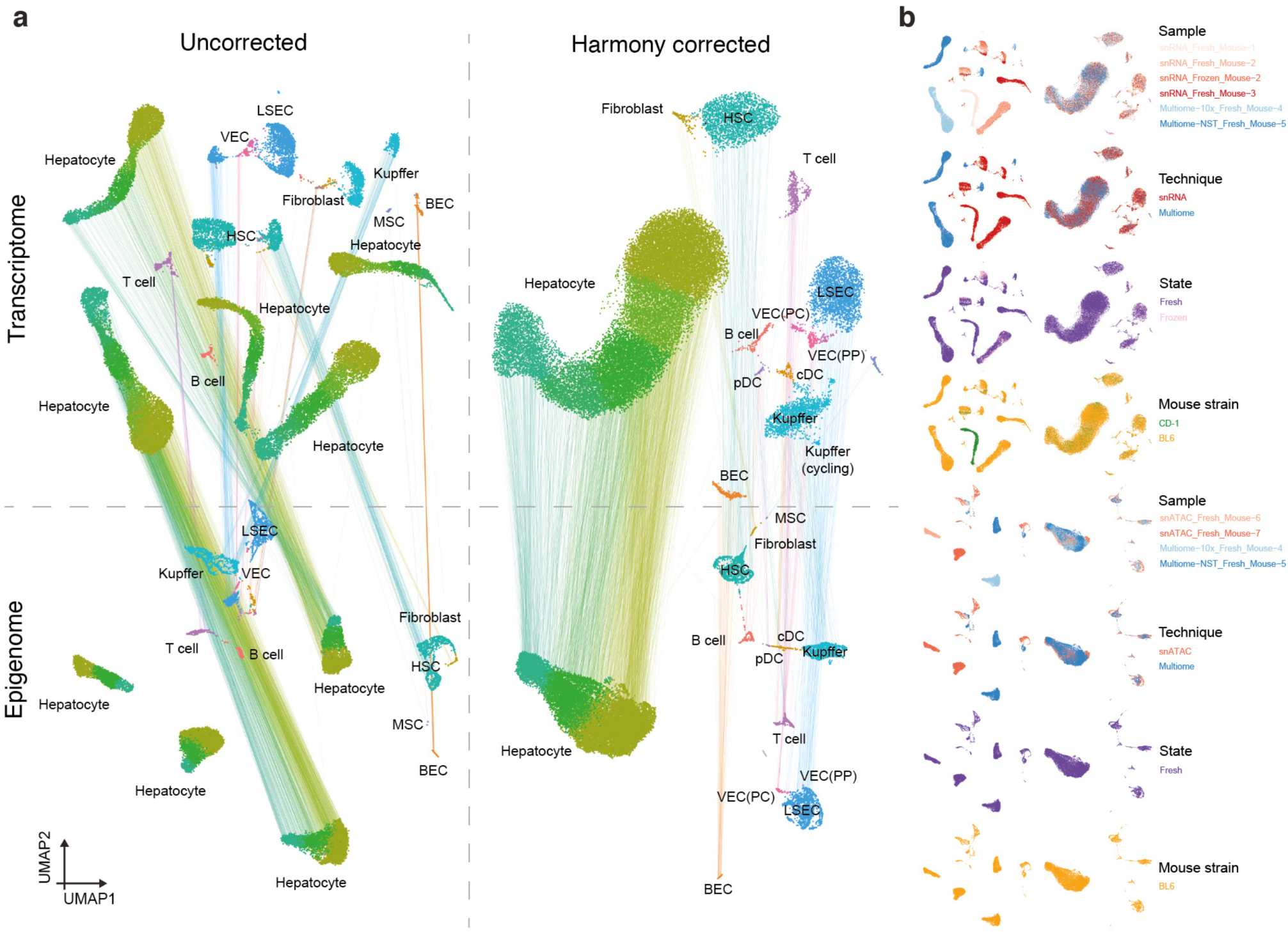
Batch effects in the mouse liver. **a.** Transcriptome and epigenome based UMAPs (29,798 and 22,600 cells, respectively) before and after batch correction with harmony. Lines linking the UMAPs map the transcriptome and the epigenome UMAP positions from the same cell (profiled by single-cell multiomics). **b.** Uncorrected and harmony corrected transcriptome and epigenome UMAP colored by (from top to bottom) sample of origin, technique (multiome or independent snRNA-seq/scATAC-seq), sample condition (fresh/ frozen) and mouse strain (CD-1, BL6).

**Figure S8.**
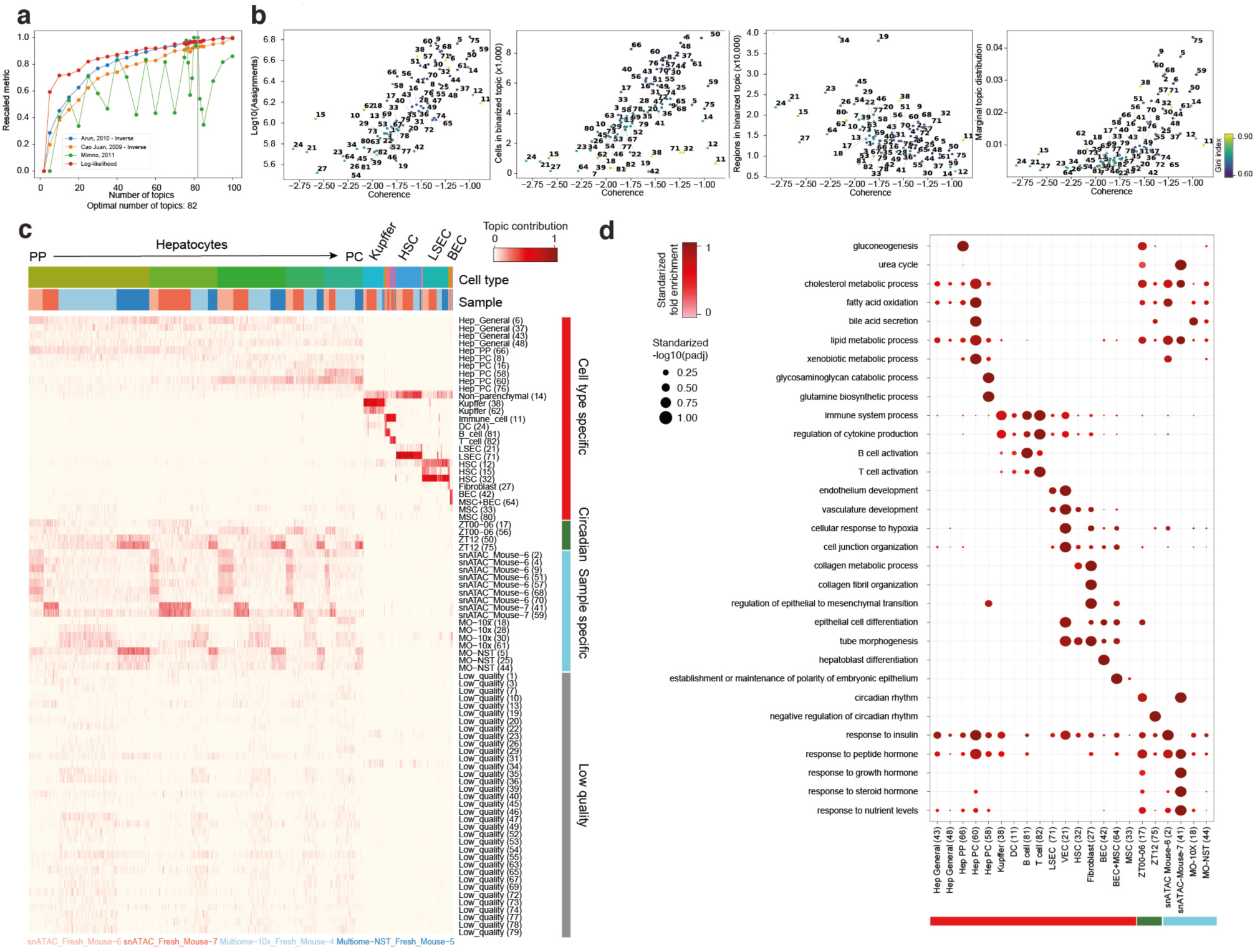
Overview of regulatory topics inferred with pycisTopic from the epigenome layer. **a.** Standardized pycisTopic topic model selection metrics, including loglikelihood, coherence (Minmo et al. 2011), reversed Cao Juan et al., 2009 and reversed Arun, 2010. The selected topic model (82 topics) is indicated with a vertical grey line. **b.** pycisTopic topic quality metrics including the number of assignments, number of regions/cells in the binarized topics and marginal topic distribution, colored by gini index value. **c.** Cell-topic heatmap (22,600 cells) showing topic contributions across cells with annotated topics. **d.** Dotplot showing relevant GO terms found by GREAT in the topics, using the -log10(padj) as size and the standardized fold enrichment as color.

**Figure S9.**
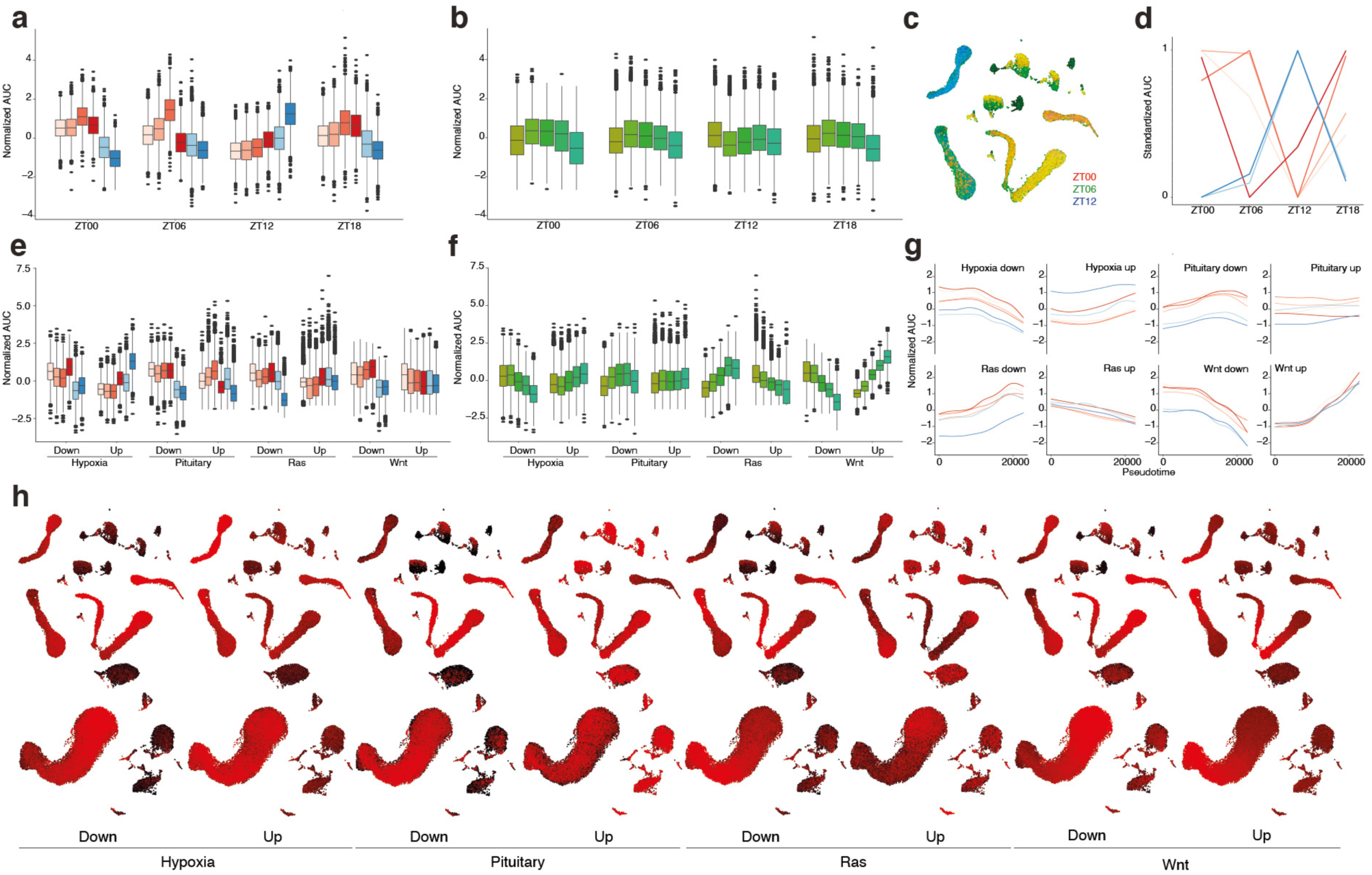
Enrichment of relevant signatures in the mouse liver on the transcriptome layer. **a.** Boxplot showing normalized AUC values across samples for circadian rhythm signatures, derived from Droin et al. 2021. **b.** Boxplot showing normalized AUC values across hepatocyte subclasses (driven by zonation) for circadian rhythm signatures. **c.** Uncorrected scRNA-seq UMAP (29,798 cells) colored by AUC values for circadian rhythm signatures using RGB encoding. **d.** Standardized AUC values across samples for signatures on different circadian rhythm time points. **e.** Boxplot showing normalized AUC values across samples for signatures from different signaling pathways affected by liver zonation, derived from Halpern et al. 2017. **f.** Boxplot showing normalized AUC values across hepatocyte subclasses (driven by zonation) for signaling pathways signatures. **g.** GAM fitted eGRN AUC profiles per sample for signaling pathway signatures along the zonation pseudotime. **h.** Uncorrected (top) and corrected (bottom) UMAP colored by AUC scores for signaling pathway signatures.

**Figure S10.**
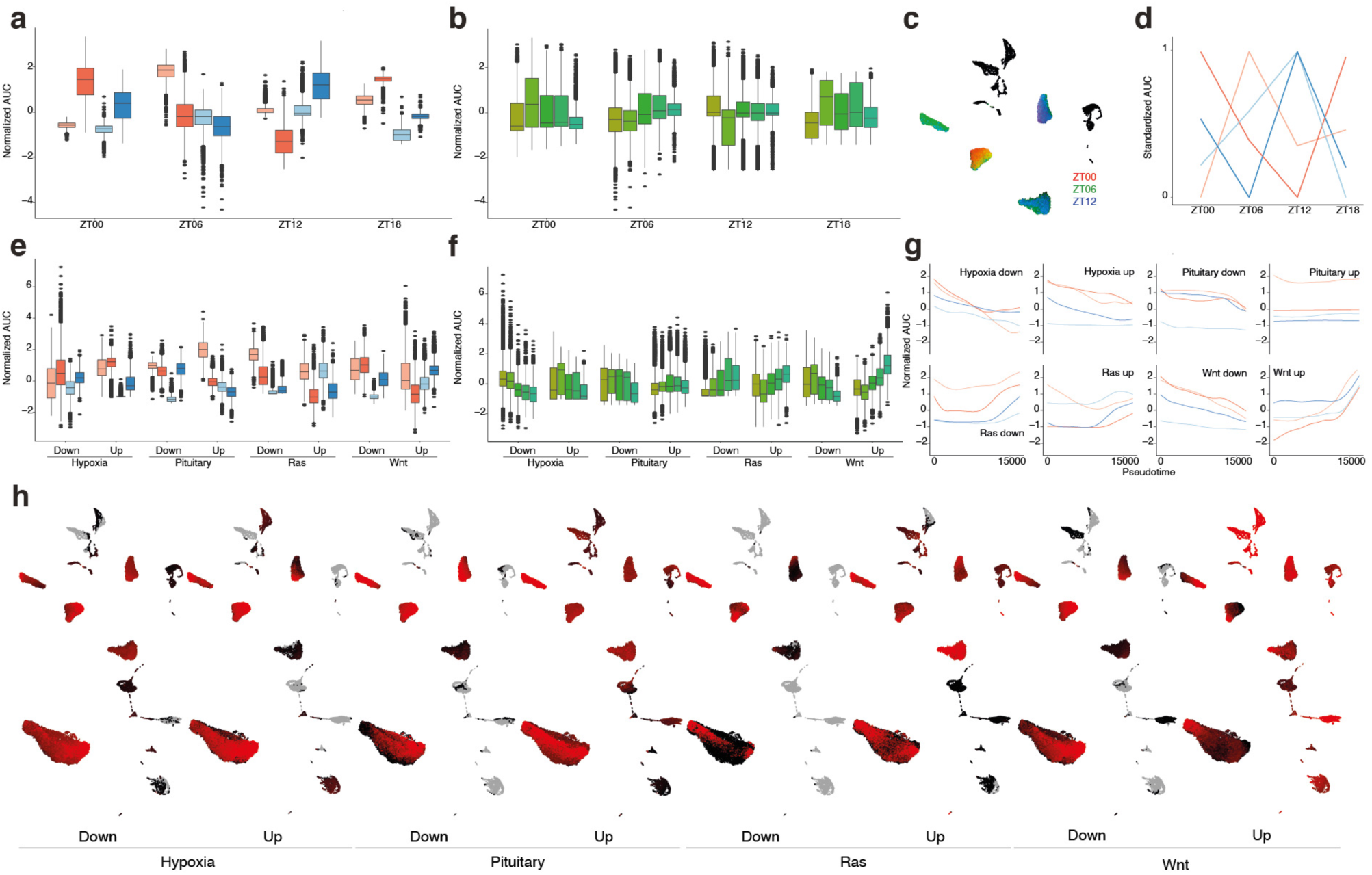
Enrichment of relevant signatures in the mouse liver on the epigenome layer, relying on gene activity. **a.** Boxplot showing normalized AUC values across samples for circadian rhythm signatures, derived from Droin et al. 2021. **b.** Boxplot showing normalized AUC values across hepatocyte subclasses (driven by zonation) for circadian rhythm signatures. **c.** Uncorrected scATAC-seq UMAP (22,600 cells) colored by AUC values for circadian rhythm signatures using RGB encoding. **d.** Standardized AUC values across samples for signatures on different circadian rhythm time points. **e.** Boxplot showing normalized AUC values across samples for signatures from different signaling pathways affected by liver zonation, derived from Halpern et al. 2017. **f.** Boxplot showing normalized AUC values across hepatocyte subclasses (driven by zonation) for signaling pathways signatures. **g.** GAM fitted eGRN AUC profiles per sample for signaling pathway signatures along the zonation pseudotime. **h.** Uncorrected (top) and corrected (bottom) UMAP colored by AUC scores for signaling pathway signatures.

**Figure S11.**
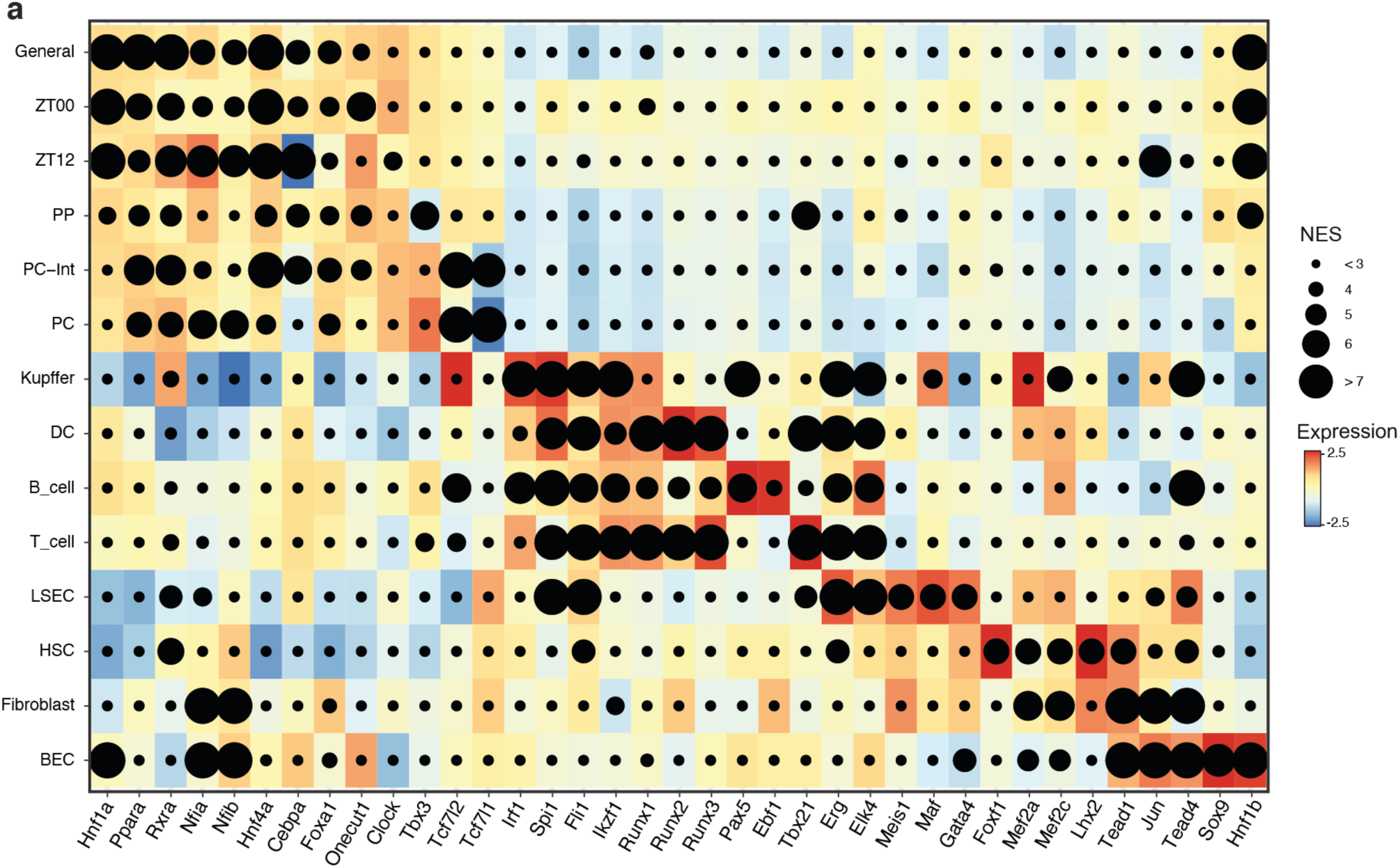
Motif enrichment on DARs/Topics on selected populations. **a.** Dotplot showing enriched motifs (linked to TFs) in different cell populations, using the maximum Normalized Enrichment Score (NES) as dot size and TF expression as color.

**Figure S12.**
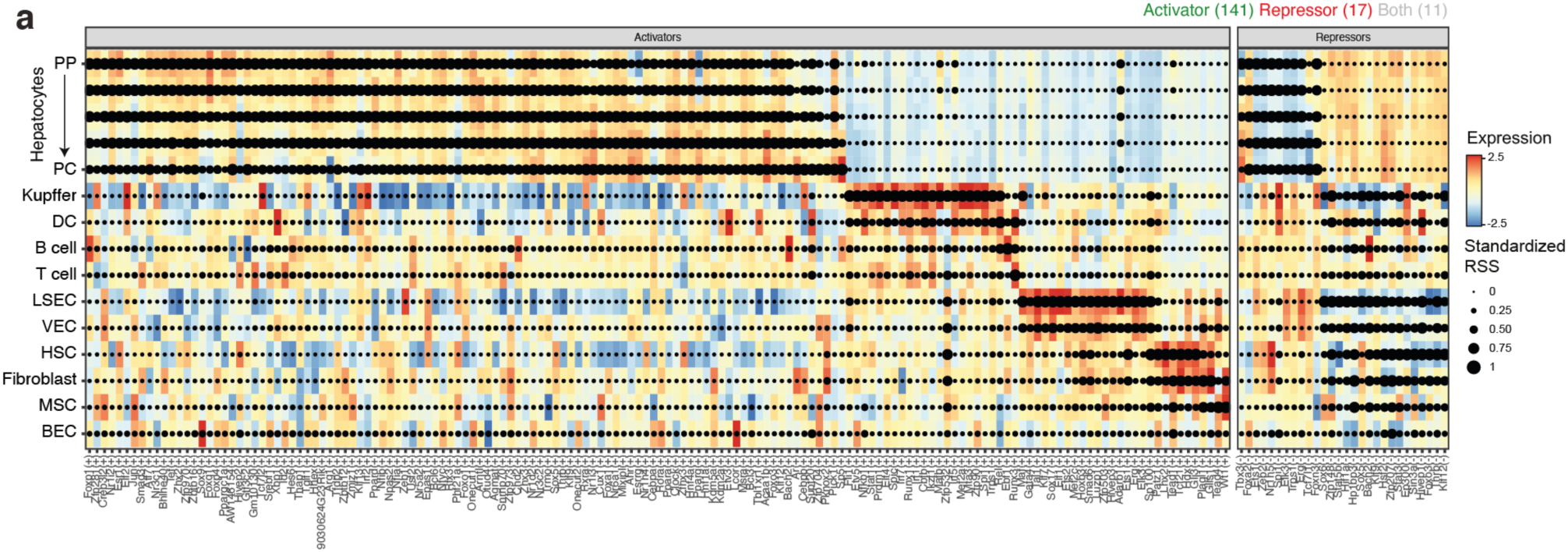
SCENIC+ regulon dotplot. **a.** Dotplot showing enriched regulons across different cell types using the maximum Regulon Specificity Score (RSS) as dot size and TF expression as color.

**Figure S13.**
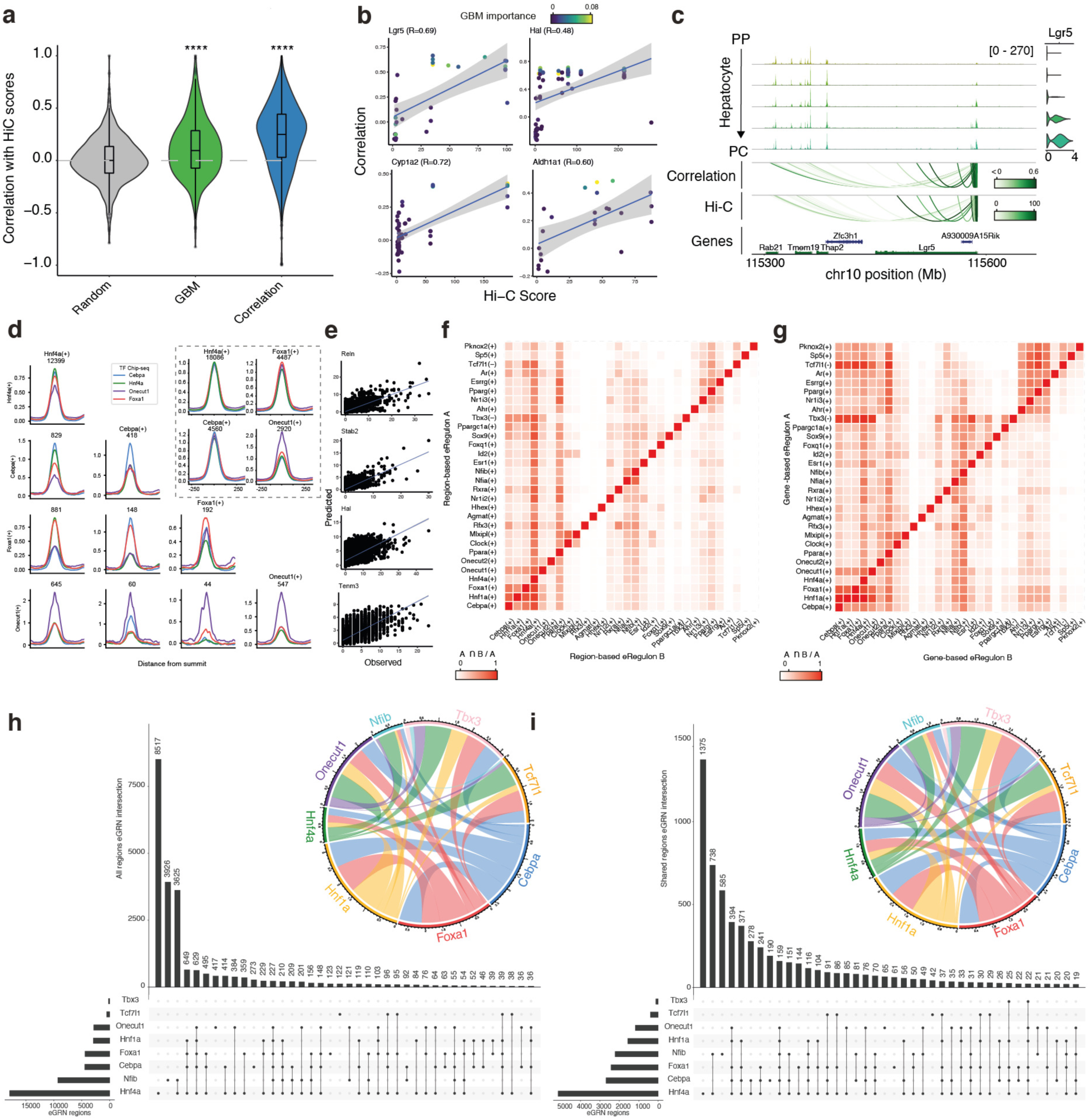
Validation of SCENIC+ regulons. **a.** Violin plot showing the correlation between SCENIC+ predicted region to gene links per gene (by the GBM and correlation methods) with Hi-C scores (between the same regions and the TSS of the linked gene). The random control distribution consists of shuffled correlation values. **b.** Examples showing the correlation between SCENIC+ region to gene links correlation scores and Hi-C scores, colored by their GBM importance. **c.** Example on the Lgr5 locus depicting chromatin accessibility profiles and gene expression across hepatocyte subpopulations and the region to gene correlation and Hi-C scores. **d.** ChIP-seq coverage profiles for Hnf4a, Cebpa, Foxa1 and Onecut1 on their unique and shared predicted regulon regions. **e.** Example showing the correlation between observed gene expression values (on left-out-data) and gene expression predicted using a GBM model (per gene) trained using the expression of the predicted TF regulators as features. **f.** Heatmap showing the overlap between selected gene-based regulons. **g.** Heatmap showing the overlap between selected region-based regulons. **h.** Overlap between all regions included in selected regulons. **i.** Overlap between core regions (i.e. accessible across all mice) included in selected regulons.

**Figure S14.**
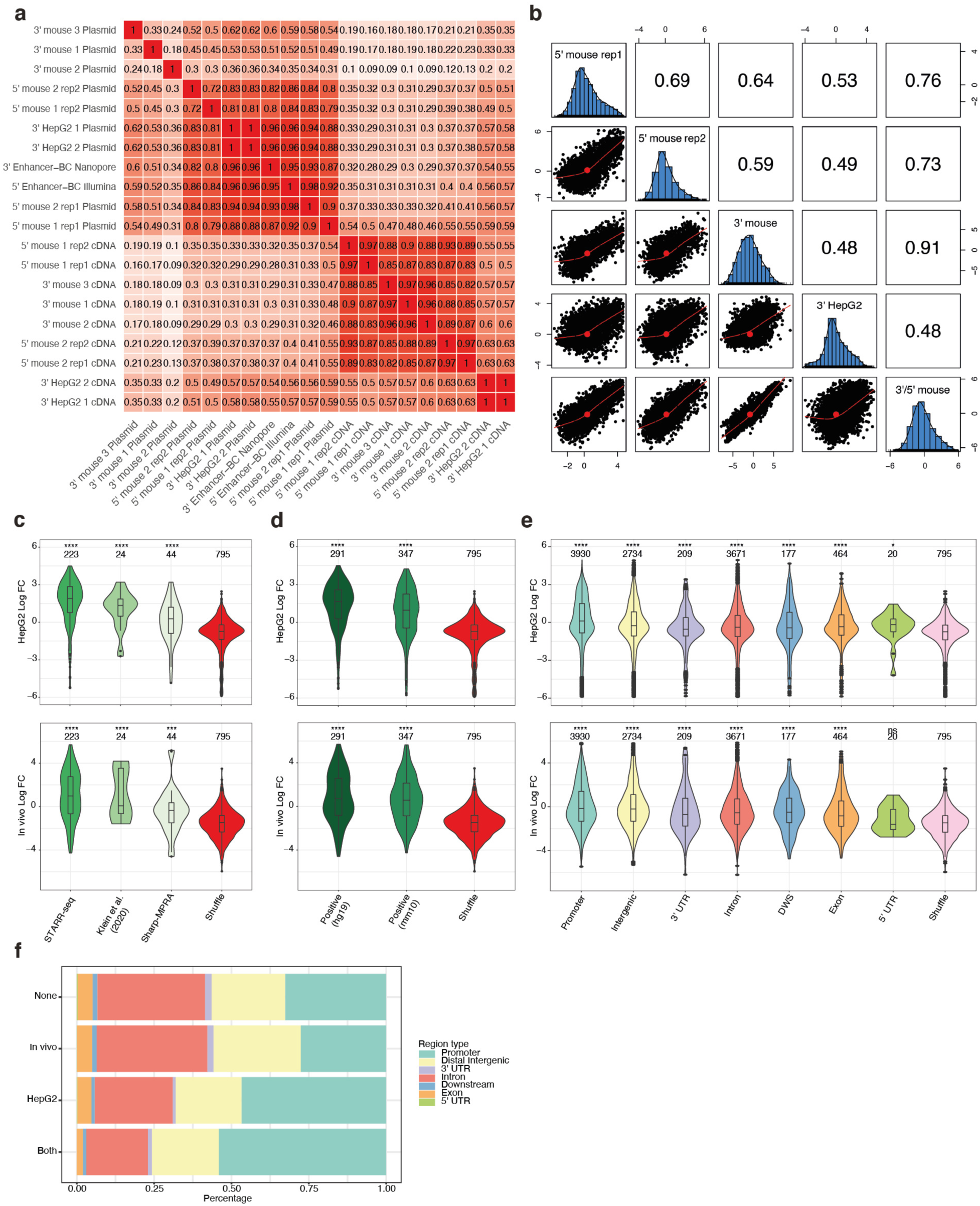
CHEQ-seq in the mouse liver and HepG2. **a.** Correlation between plasmid and cDNA measurements across MPRA experiments in the mouse liver and HepG2. **b.** Correlation of the DESeq LogFC values per enhancer across experiments. **c.** CHEQ-seq Log Fold-Change violin plots per positive control class. **d.** CHEQ-seq Log Fold-Change violin plots per positive control sequence class (whether it is hg19 or mm10 based). **e.** CHEQ-seq Log Fold-Change violin plots per sequence genomic annotation. For **c-e**, the asterisks indicate the significance compared to Shuffle (****: p-value <= 0.0001, ***: p-value <= 0.001, *: p-value <= 0.05, ns: p-value > 0.05). **f.** Proportion of enhancer classes based on genomic annotation per high confidence activity class. None: Not active (4,285), In vivo: Active only *in vivo* (806), HepG2: Active only in HepG2 (921), Both: Active in HepG2 and *in vivo* (1,186).

**Figure S15.**
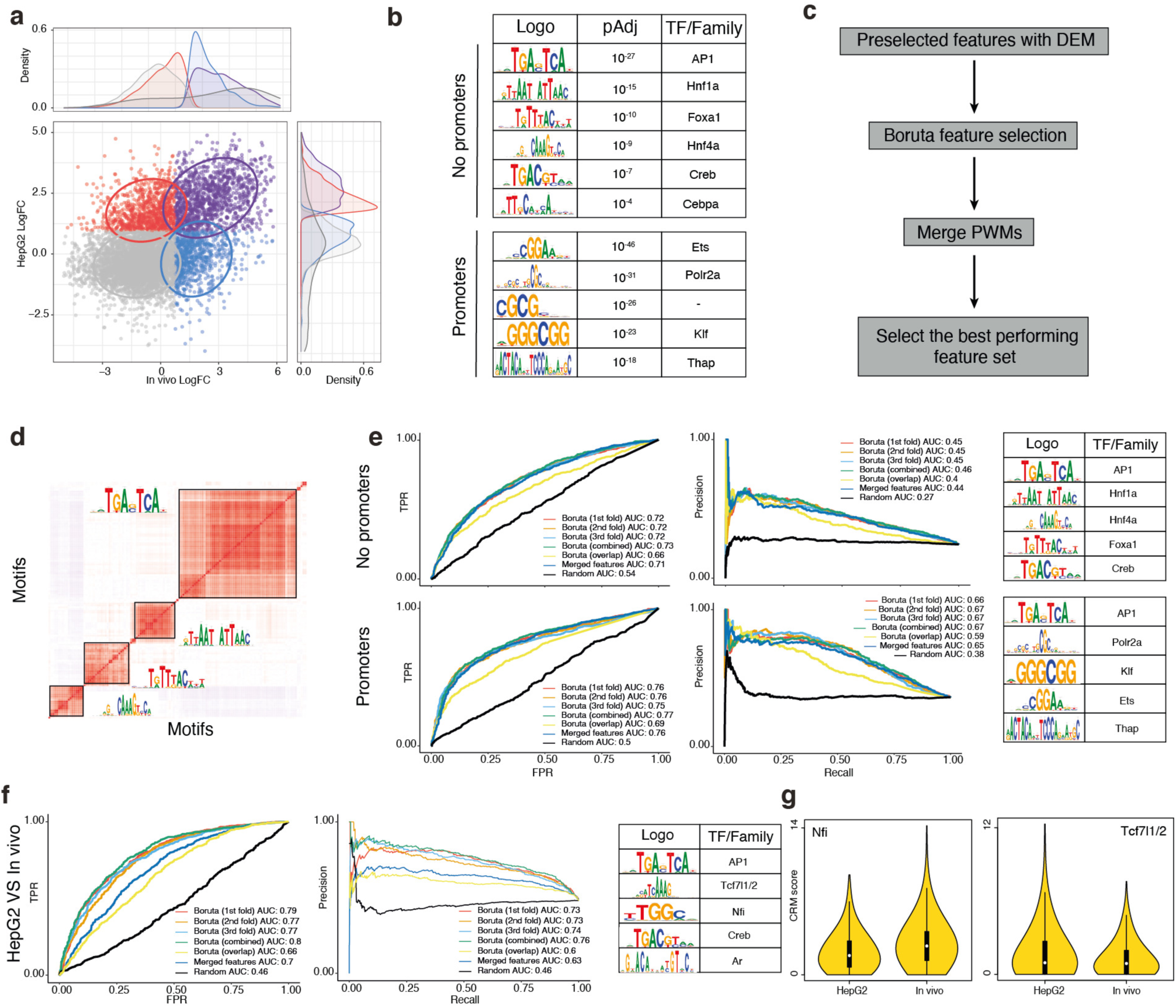
Random Forest models allow to identify sequence features driving enhancer activity. **a.** Correlation between Log Fold-Changes for high confidence enhancers (7,198) in Hepg2 and *in vivo* colored by enhancer type, with data ellipses per activity group: Not active (grey), active in HepG2 (red), active *in vivo* (blue), active both *in vivo* and HepG2 (purple). **b.** Features identified by DEM when comparing active enhancer in vivo versus not active. The analysis was performed with all regions and removing promoters (regions within 1kb from a TSS). **c.** Overview of the random forest-based feature selection workflow. **d.** Correlation between CRM scores of features selected with Boruta (merged from 3 folds). Correlated features represent sets of similar PWM, of which one is shown as a representative example. **e.** ROC and precision-recall curves for the trained activity models (with and without promoters). Boruta using 3-fold validation, and models were trained per fold, using all features found in at least one fold or only overlapping features. Merged features are derived by using all features found in at least one fold and merged based on their CRM score correlation and motif similarity. Top selected features per model (ordered by importance) are shown on the table on the right. **f.** ROC and precision-recall curves for the activity models trained to distinguish differential features between regions only active in HepG2 or only active *in vivo*. Top selected features per model (ordered by importance) are shown on the table on the right. **g.** Violin plots showing Nfia/b and Tcf7l1/2 motifs CRM score distributions on HepG2 and *in vivo*.

**Figure S16.**
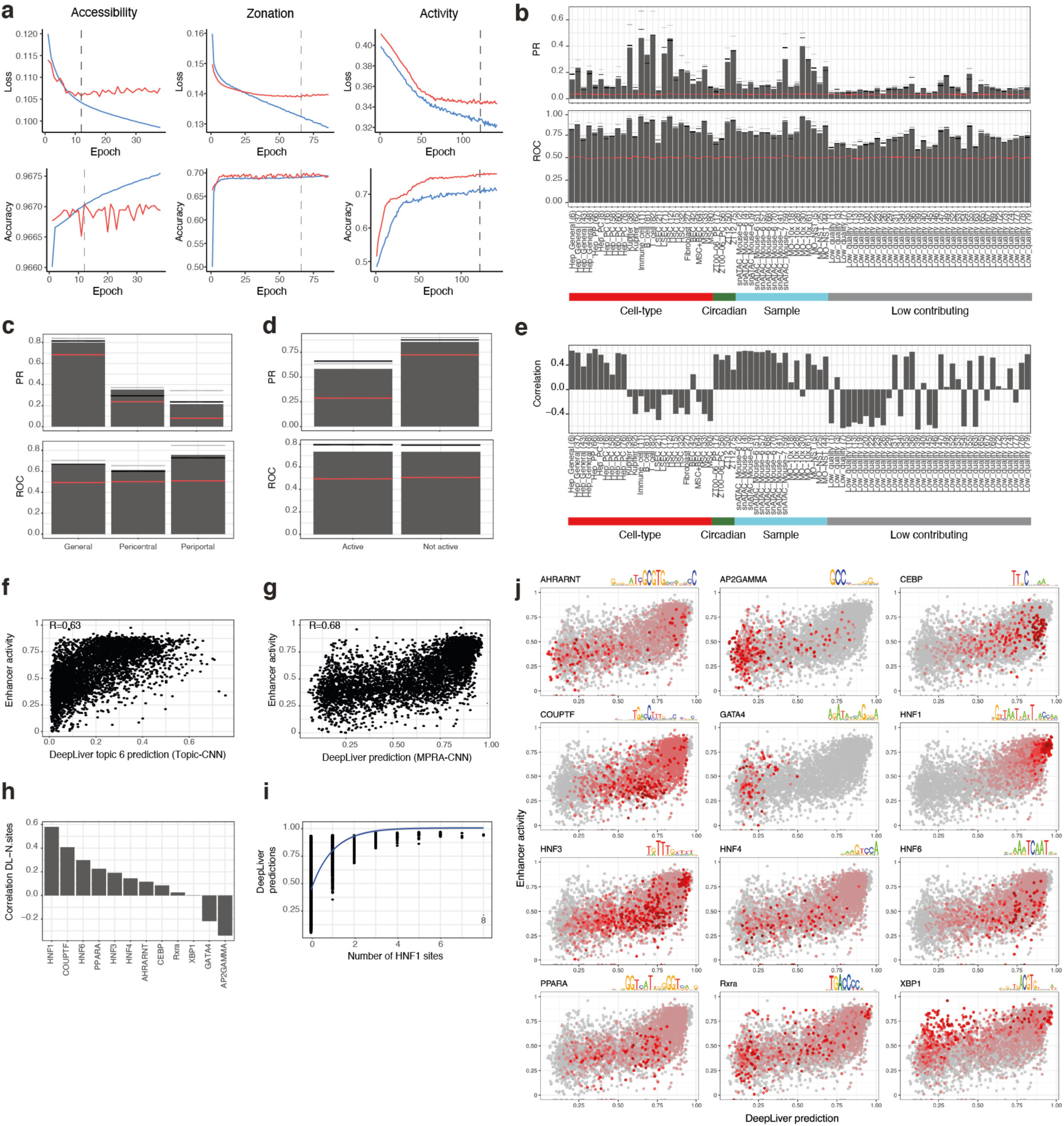
Overview of DeepLiver models and their predictability on synthetic enhancers from Smith et al. 2013. **a.** Loss and accuracy curves for the DeepLiver models. The grey dashed lines indicate the selected epochs per model. **b.** ROC and PR values on test data per topic for the DeepLiver accessibility model. The red line shows the values for a random classifier; the grey line, on the training data; and the black line on the validation data. **c.** ROC and PR values per topic for the DeepLiver zonation model. The red line shows the values for a random classifier; the grey line, on the training data; and the black line on the validation data. **d.** ROC and PR values per topic for the DeepLiver activity model. The red line shows the values for a random classifier; the grey line, on the training data; and the black line on the validation data**. e.** Correlation between DeepLiver Topic-CNN predictions and Smith et al. 2013 enhancer activity. **f.** Correlation plot between Smith et al. 2013 enhancer activity and DeepLiver accessibility predictions (Topic-CNN). **g.** Correlation plot between Smith et al. 2013 enhancer activity and DeepLiver activity predictions (MPRA-CNN). **h.** Barplot showing the correlation between DeepLiver predicted activity scores and the number of sites for each motif. **i.** Plot showing DeepLiver activity predictions versus the number of Hnf1a sites. The blue line depicts a saturation curve. **j.** Correlation plot between Smith et al. 2013 enhancer activity and DeepLiver activity predictions colored by the number of motif instances in the sequences (red scale).

**Figure S17.**
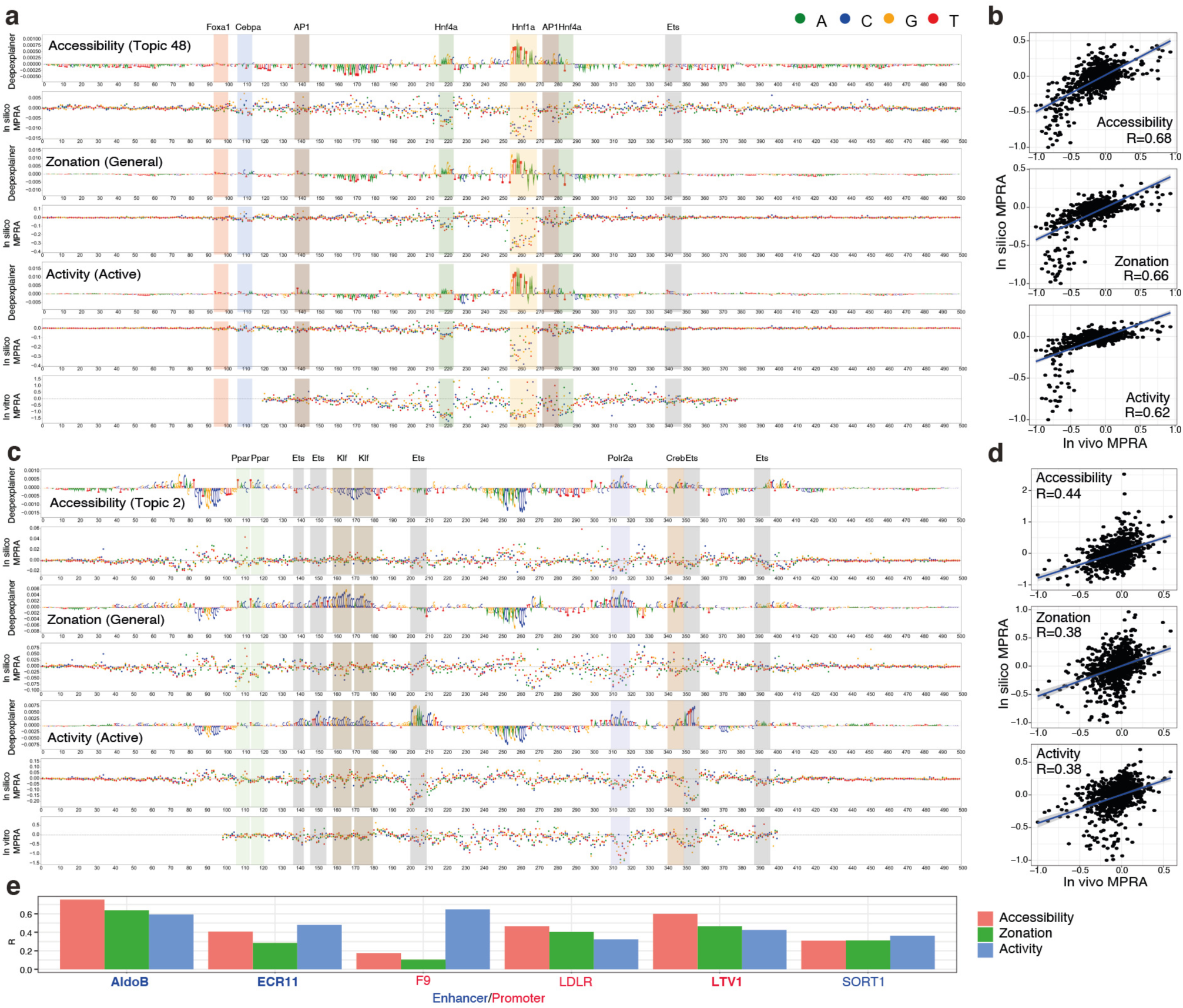
DeepLiver *in silico* saturation mutagenesis MPRA predictions compared to experimental results. **a.** DeepExplainer and saturation mutagenesis plots for the accessibility, zonation and activity models on an AldoB enhancer (hg19: chr9:104195449-104195449), with motifs highlighted. Saturation mutagenesis, shown below, was performed in this enhancer by Patwardhan et al., 2012. **b.** Correlation between DeepLiver in silico mutagenesis and experimental saturation mutagenesis in the AldoB enhancer. **c.** DeepExplainer and saturation mutagenesis plots for the accessibility, zonation and activity models on the LTV1 promoter (mm9: chr7:29161343-29161843), with motifs highlighted. Saturation mutagenesis, shown below, was performed in this enhancer by Patwardhan et al., 2012. **d.** Correlation between DeepLiver *in silico* mutagenesis and experimental saturation mutagenesis in the LTV1 promoter. **e.** Correlation between *in silico* and experimental saturation mutagenesis for different sequences tested *in vivo* by Patwardhan et al., 2012 (AldoB, ECR11 and LTV1) and in HepG2 by Kircher et al. 2019 (F9, LDLR and SORT1).

**Figure S18.**
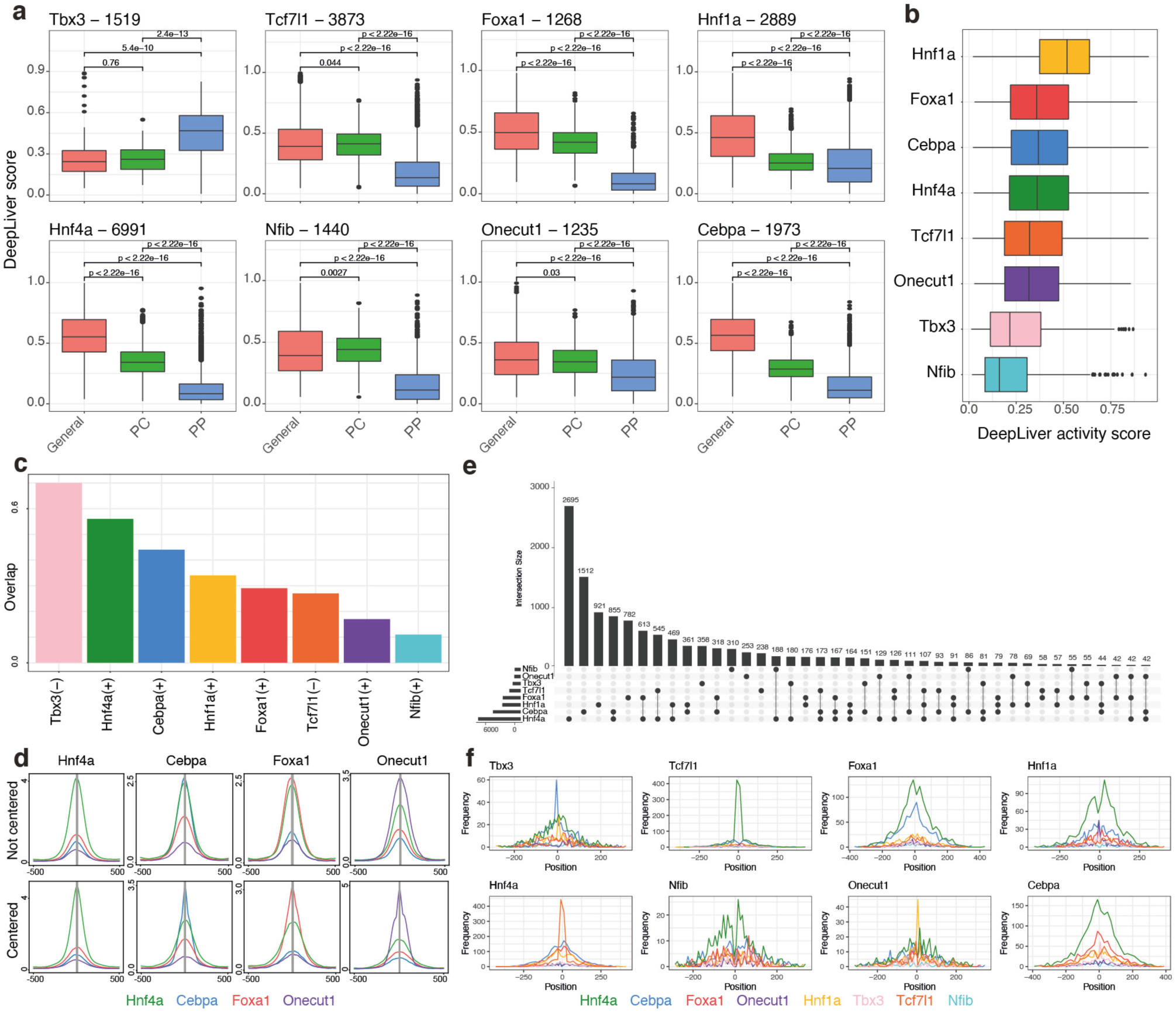
DeepLiver predictions in SCENIC+ regulons and validation of predicted target sites. **a.** DeepLiver zonation predictions on DeepLiver predicted target regions for different transcription factors. **b.** DeepLiver activity predictions on DeepLiver predicted target regions for different transcription factors. **c.** Percentage of TF target regions predicted by SCENIC+ found by DeepLiver. **d.** ChIP-seq coverage on TF target regions predicted by DeepLiver, without centering (i.e. ATAC peak coordinates) or centering on the predicted binding site. **e.** Overlap between target regions predicted by DeepLiver for different transcription factors. **f.** Distances between binding sites for a TF and binding sites of other TFs in overlapping regions.

**Figure S19.**
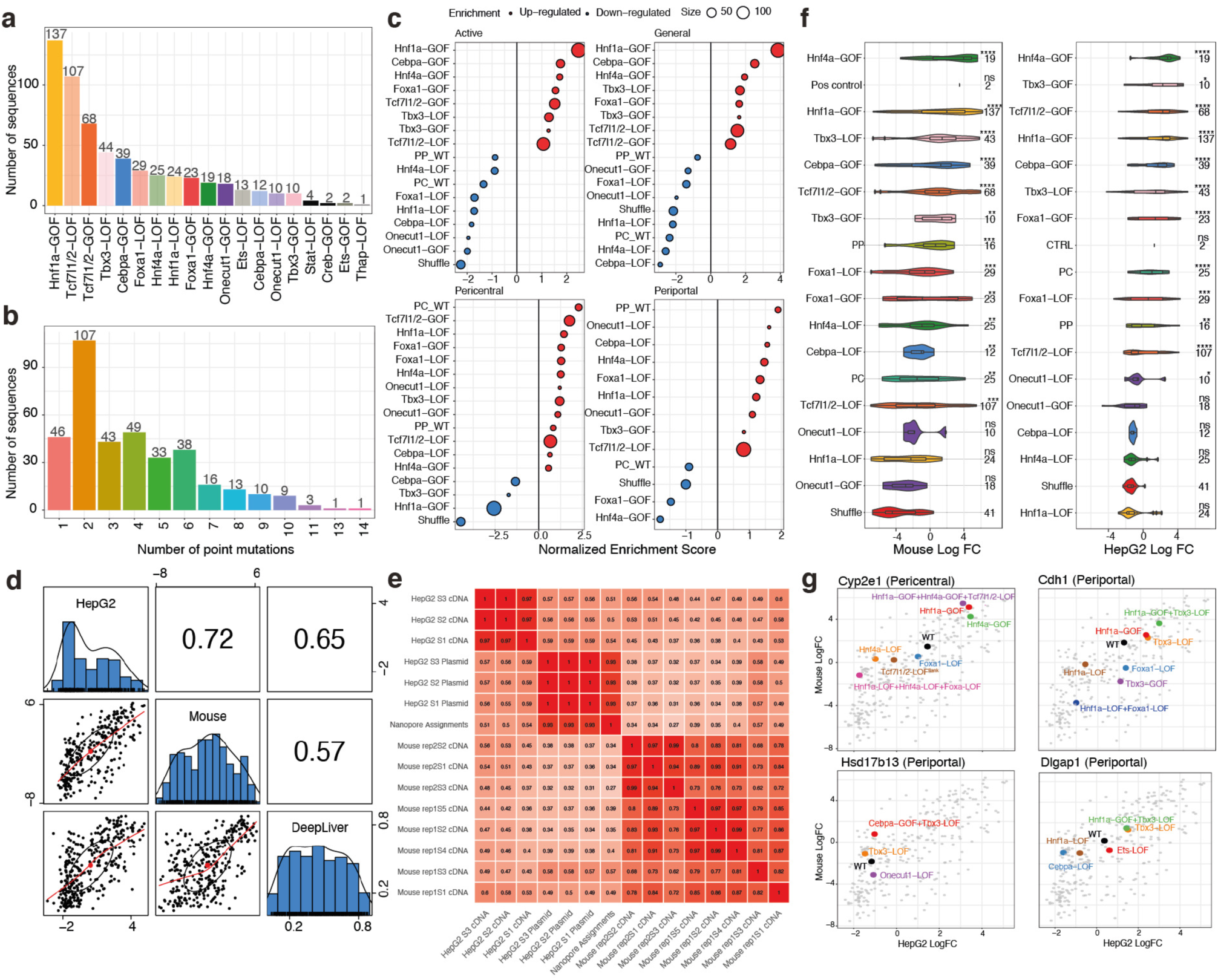
Library design and validation on wild-type zonated sequences and their activity and zonation variants. **a.** Number of sequences containing each variant type. **b.** Number of point mutations across library sequences. **c.** NES scores for variant types along the predicted DeepLiver scores for activity and zonation (general, pericentral and periportal). **d.** Correlation between CHEQ-seq Log Fold-Changes *in vivo* and in HepG2 and DeepLiver activity predictions. **e.** Correlation between number of counts per enhancer across samples and replicates. **f.** CHEQ-seq Log Fold-Changes per variant type *in vivo* and in HepG2. **g.** *In vivo* CHEQ-seq Log Fold change versus DeepLiver activity score with highlighted sequence variants for each enhancer.

**Figure S20.**
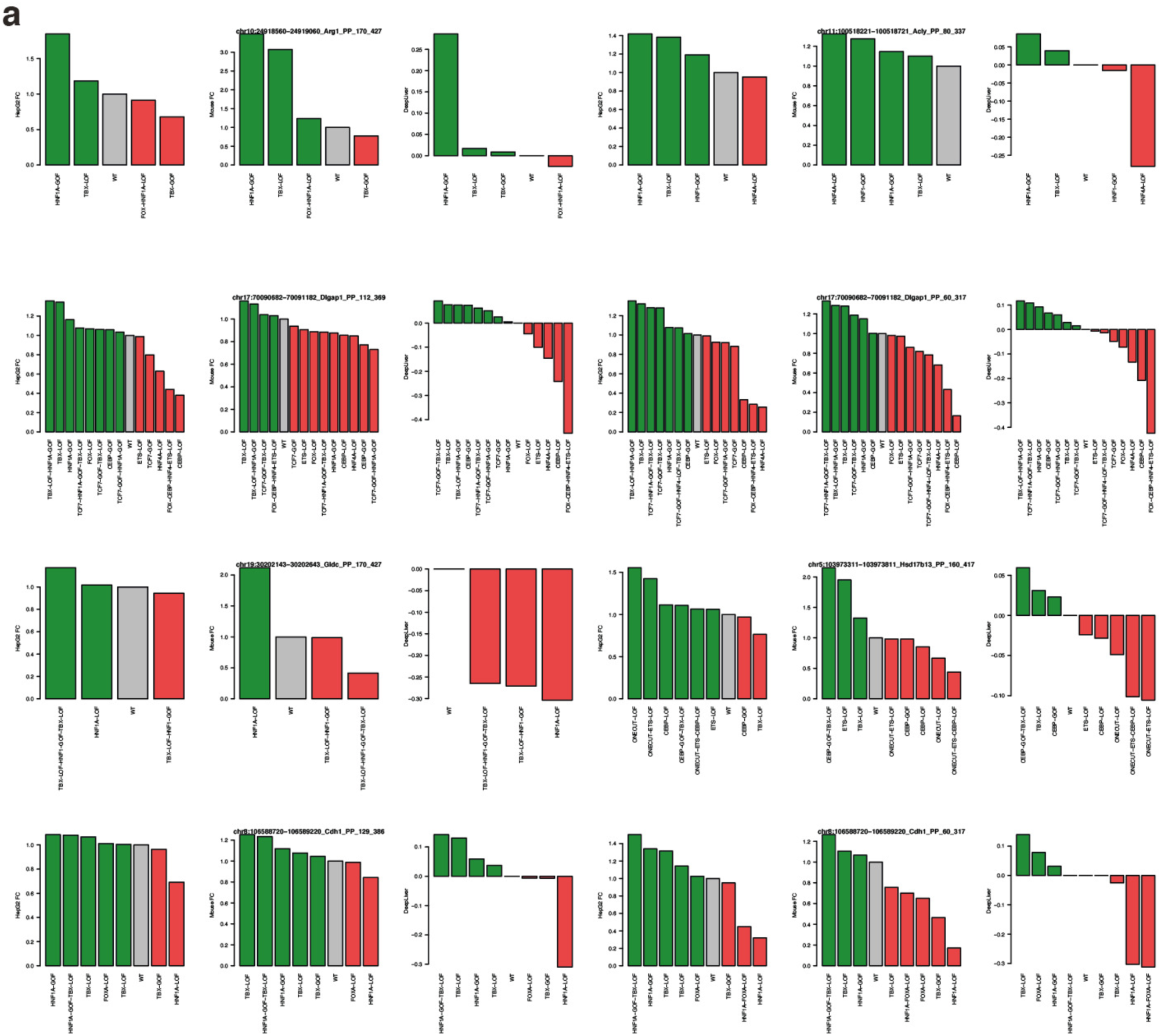
Activity shift between periportal enhancers and their variants. **a.** Comparison between activity shift between a periportal enhancer and its variants based on MPRA measurements *in vivo* and in HepG2 (Fold Change) and DeepLiver predictions. Green indicates that the variant is more active than the wild type sequences (ratio > 1 or DeepLiver shift > 0) and red indicates that the variant is less active than the wild type sequences (ratio < 1 or DeepLiver shift < 0).

**Figure S21.**
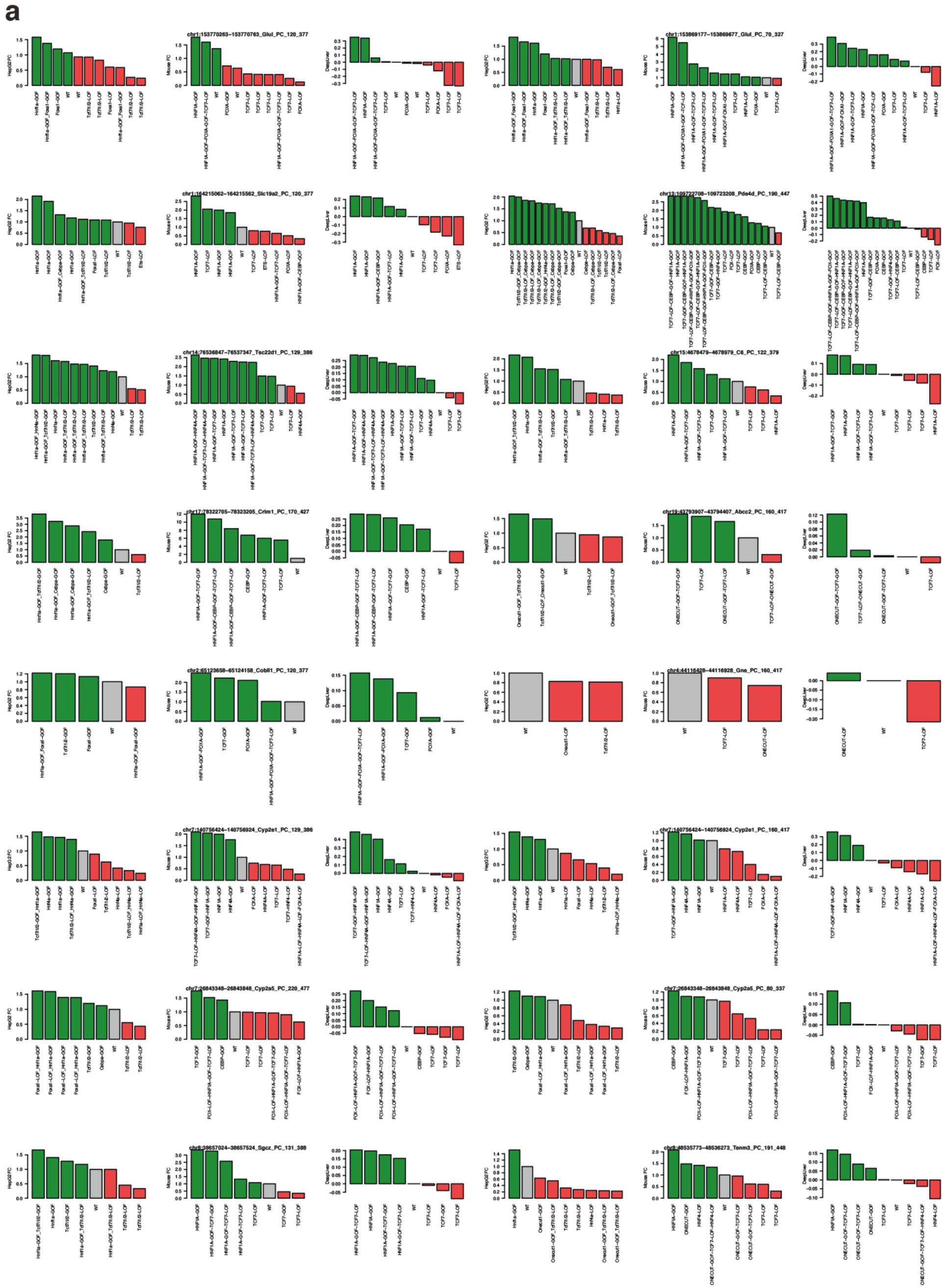
Activity shift between pericentral enhancers and their variants. **a.** Comparison between activity shift between a pericentral enhancer and its variants based on MPRA measurements *in vivo* and in HepG2 (Fold Change) and DeepLiver predictions. Green indicates that the variant is more active than the wild type sequences (ratio > 1 or DeepLiver shift > 0) and red indicates that the variant is less active than the wild type sequences (ratio < 1 or DeepLiver shift < 0).

**Figure S22.**
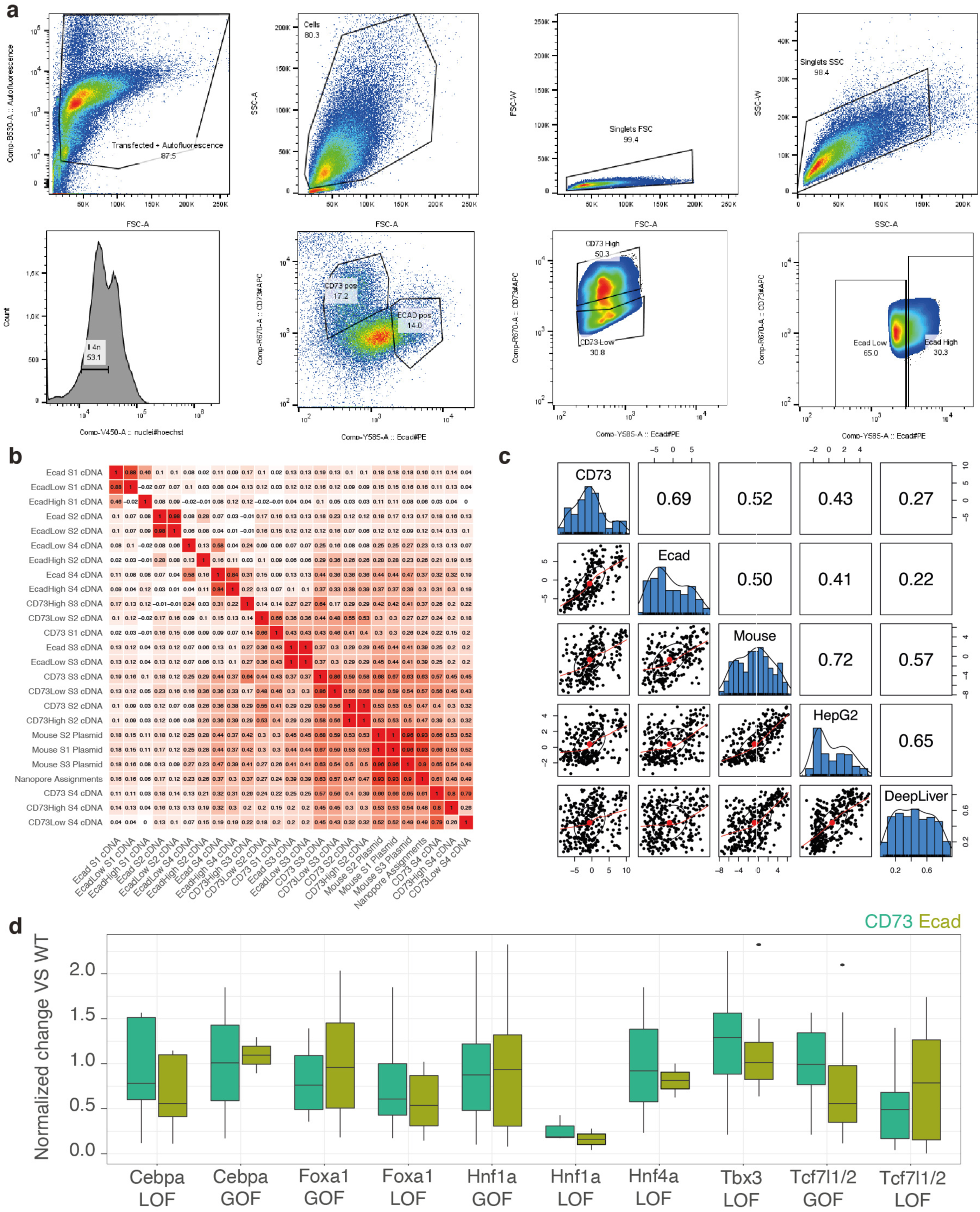
FACS MPRA reveals activity differences between pericentral and periportal enhancers. **a.** FACS gating strategy. FSC-A and SSC-A were used for hepatocytes size selection. For the FACS MPRA experiment, cells containing the library were selected based on GFP. For the FACS ATAC experiment, viable cells were selected using the Zombie Green Viability kit. Tetraploid hepatocytes were selected based on Hoechst stain. CD73 and Ecad were used to select hepatocytes bins along the porto-central axis. **b.** Correlation between number of counts per enhancer across samples and replicates. **c.** Correlation between CHEQ-seq Log Fold-Changes in the CD73^+^ and Ecad^+^ fractions, *in vivo* bulk and in HepG2 and DeepLiver activity predictions. **d.** Normalized change between enhancers (grouped by variant type) in the CD73^+^ and Ecad^+^ fractions.

**Figure S23.**
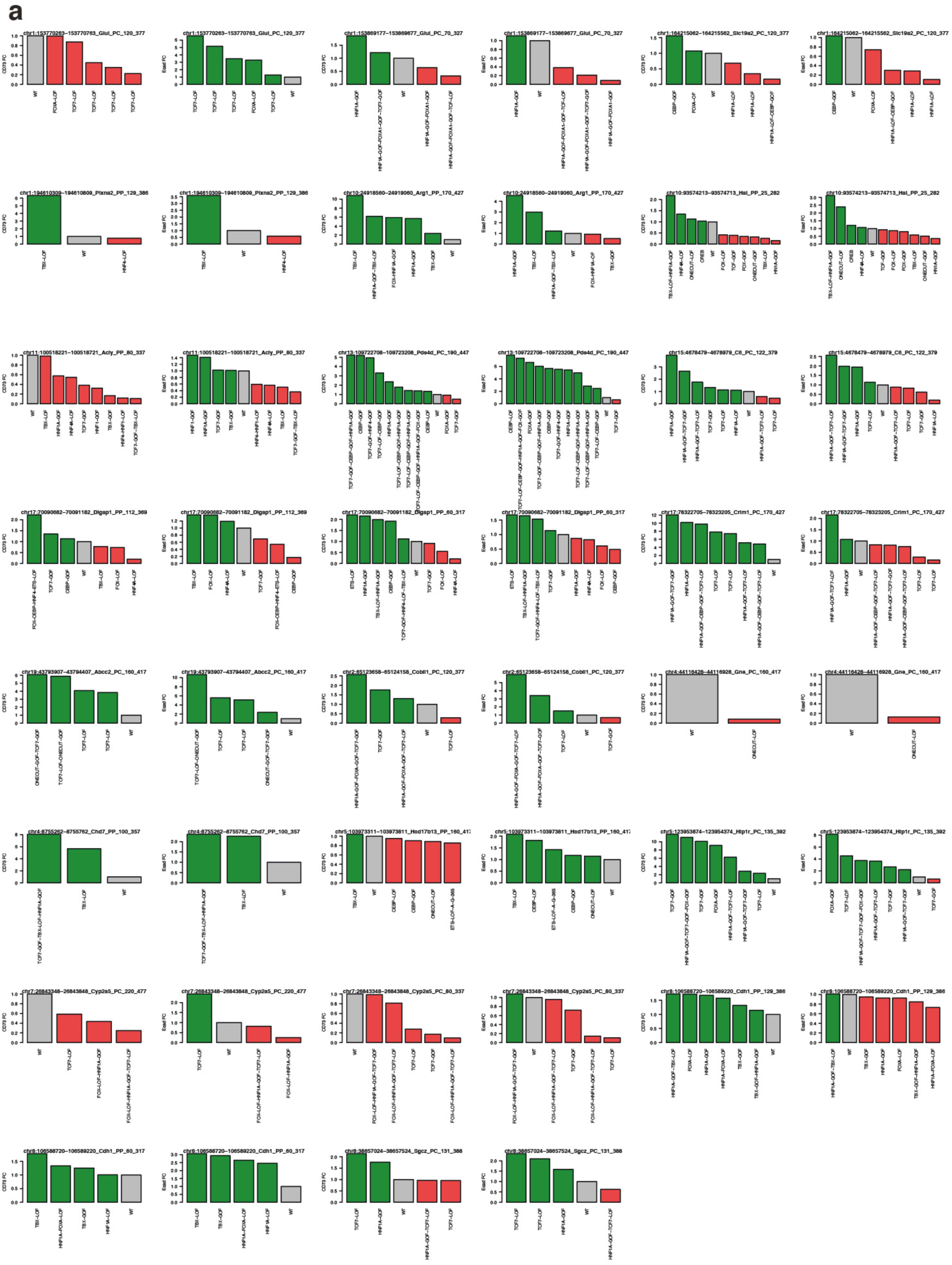
Activity shift between enhancers and their variants. **a.** Comparison between activity shift between an enhancer and its variants based on MPRA measurements in the CD73^+^ and Ecad^+^ fractions (fold change). Green indicates that the variant is more active than the wild type sequences (ratio > 1 or DeepLiver shift > 0) and red indicates that the variant is less active than the wild type sequences (ratio < 1 or DeepLiver shift < 0).

**Supplementary Note 1: DeepExplainer profiles on zonated enhancers**

In this note we include DeepExplainer plots for each of the wild-type zonated enhancers selected for the library design in Figure 5. The DeepExplainer plots correspond to the accessibility model (Topic 43), zonation model (pericentral/periportal) and activity model (active). The grey horizontal lines indicate the 259bp windows that were included in the library. Relevant motifs are highlighted. The ScoMAP liver lobule shows the region accessibility (red) and target gene expression (green). The uncorrected scATAC-seq UMAP is shown to confirm that the region is shared. The UCSC plot shows the hepatocytes pseudobulk profiles (from periportal to pericentral).

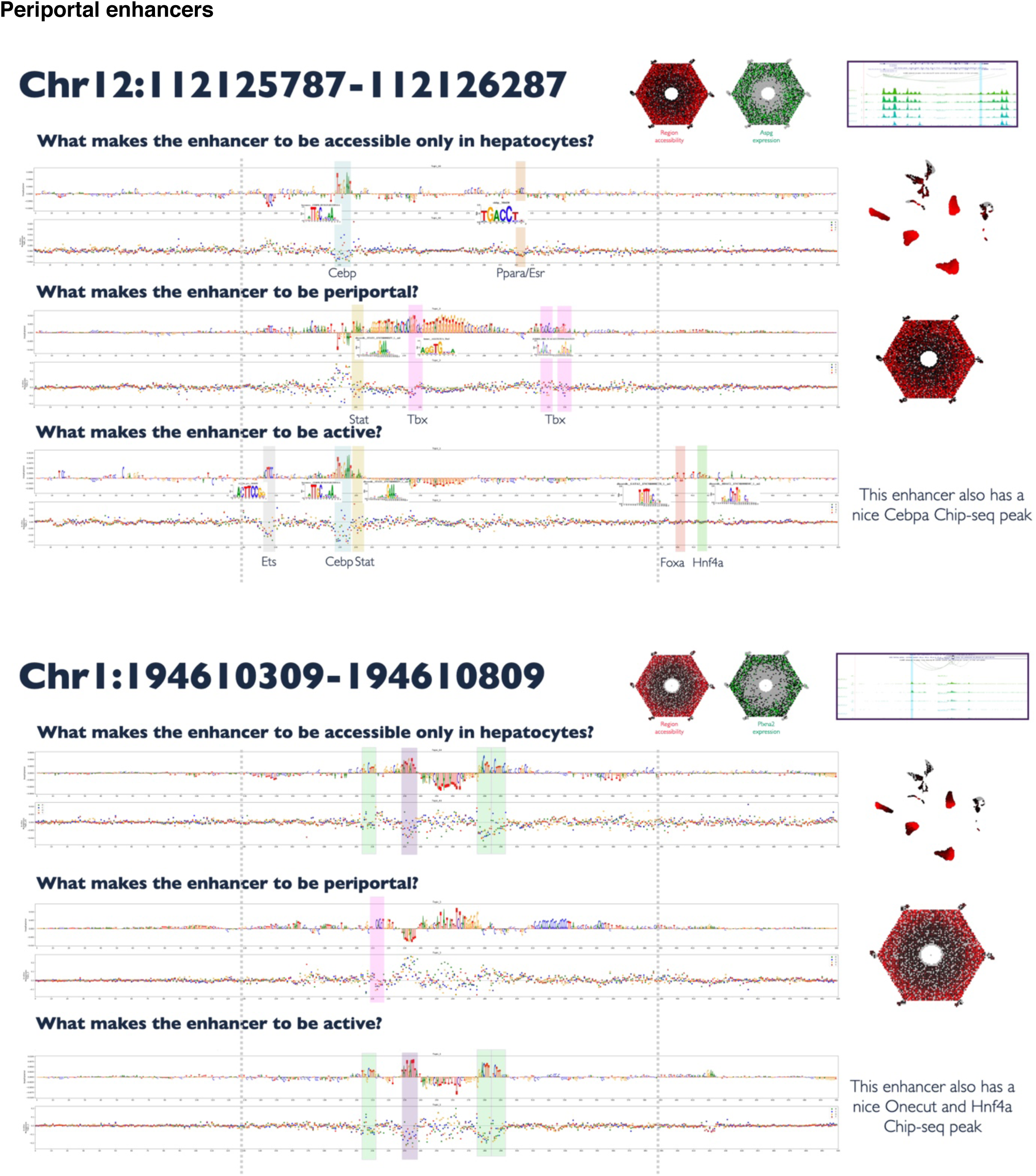
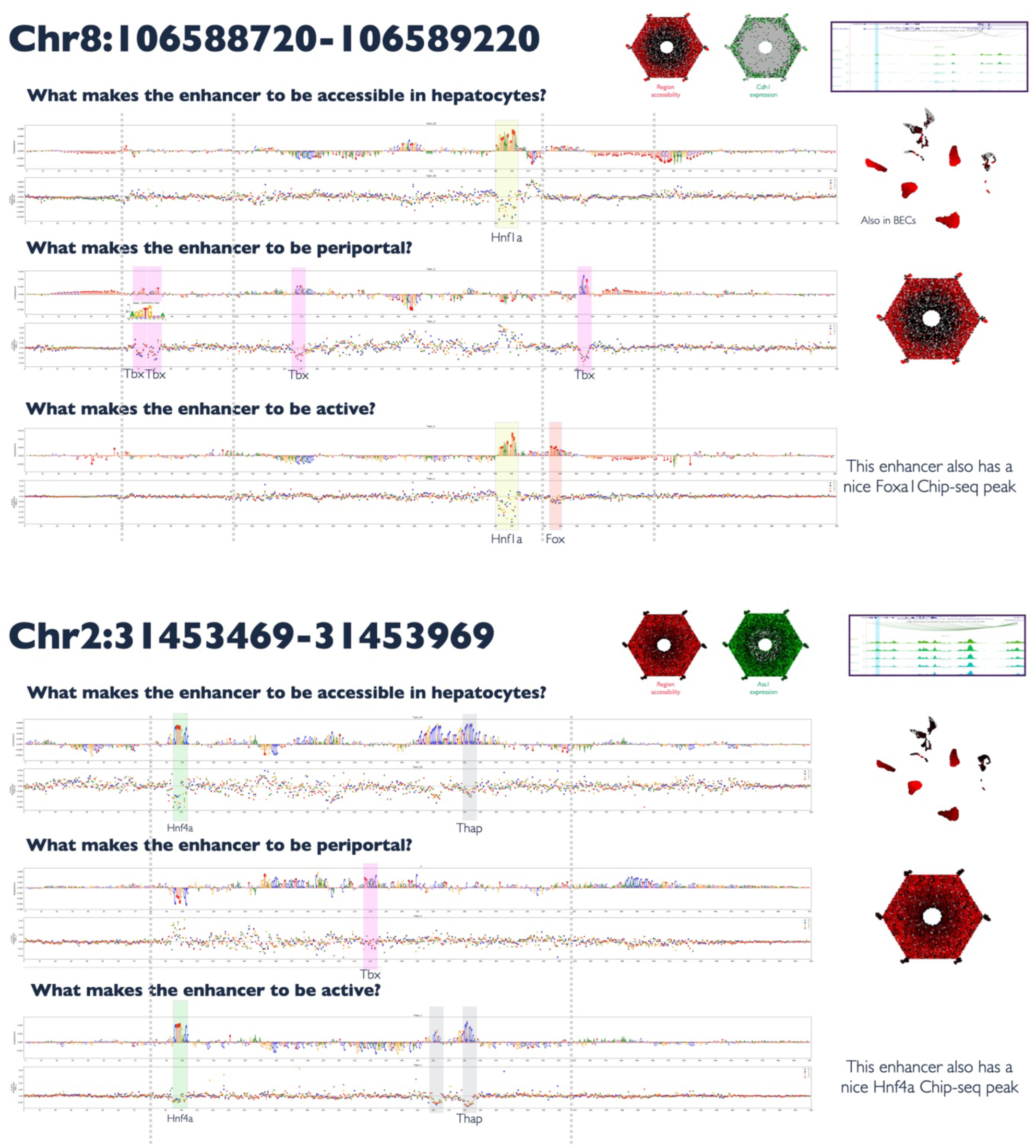
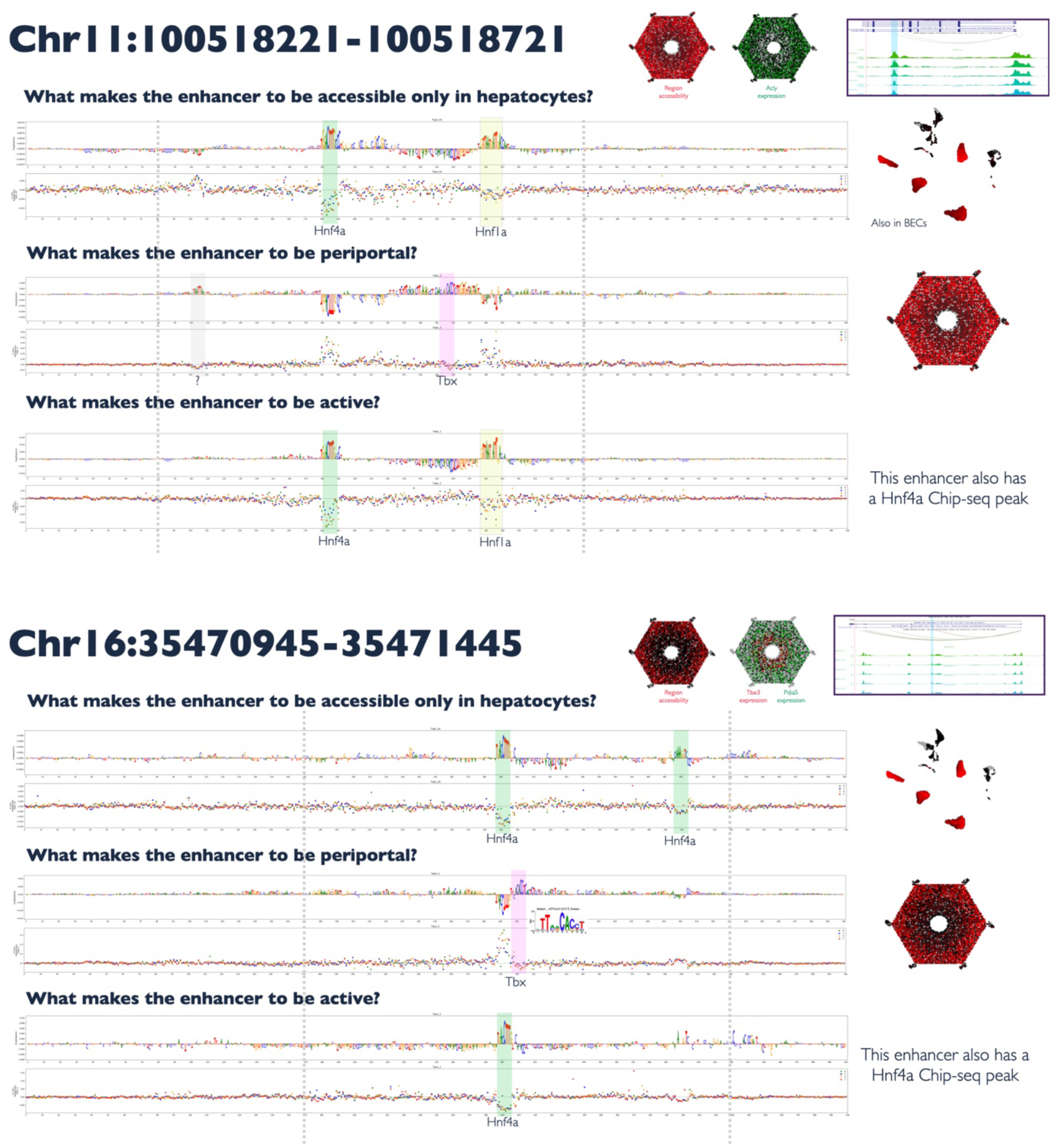
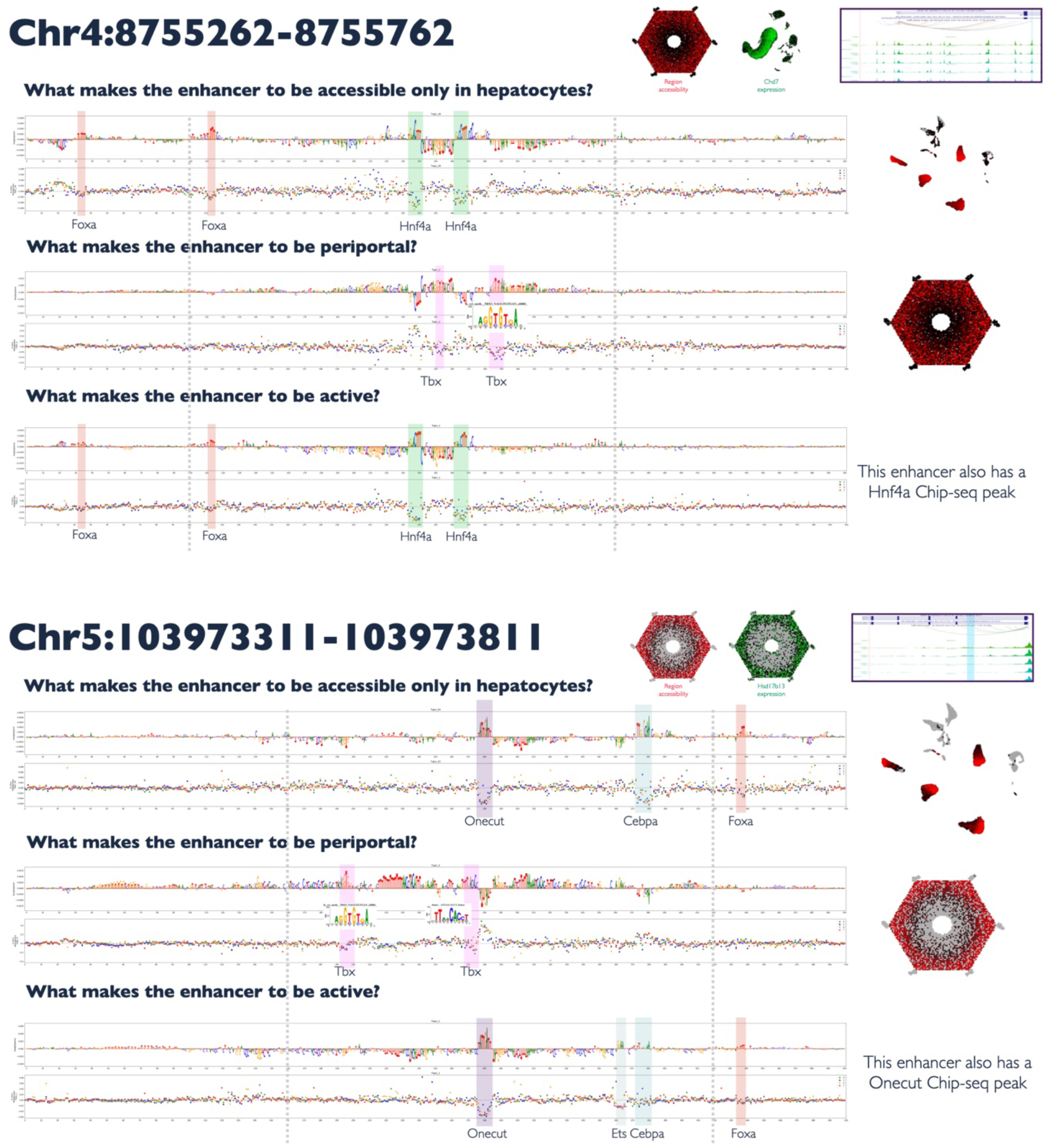
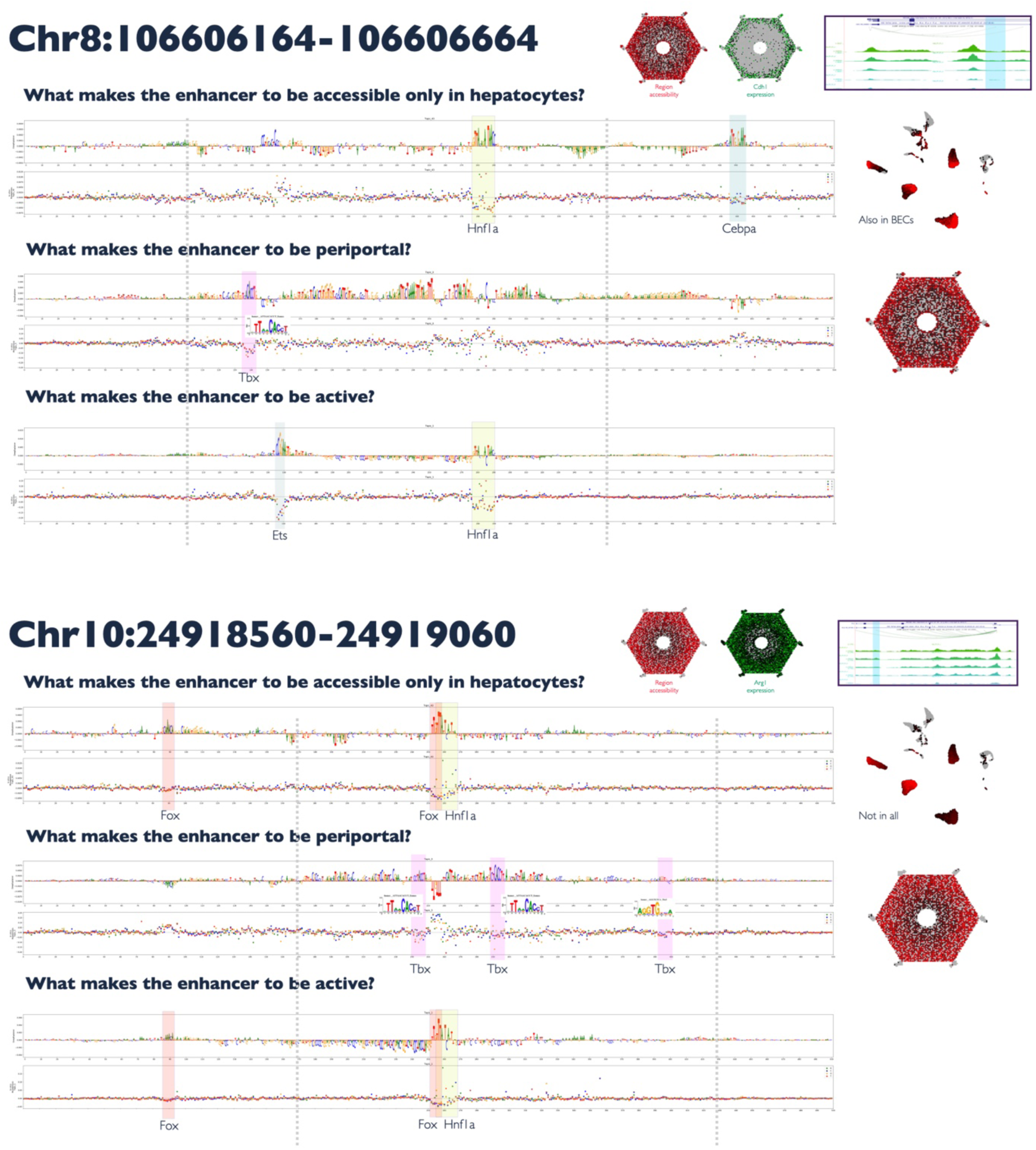
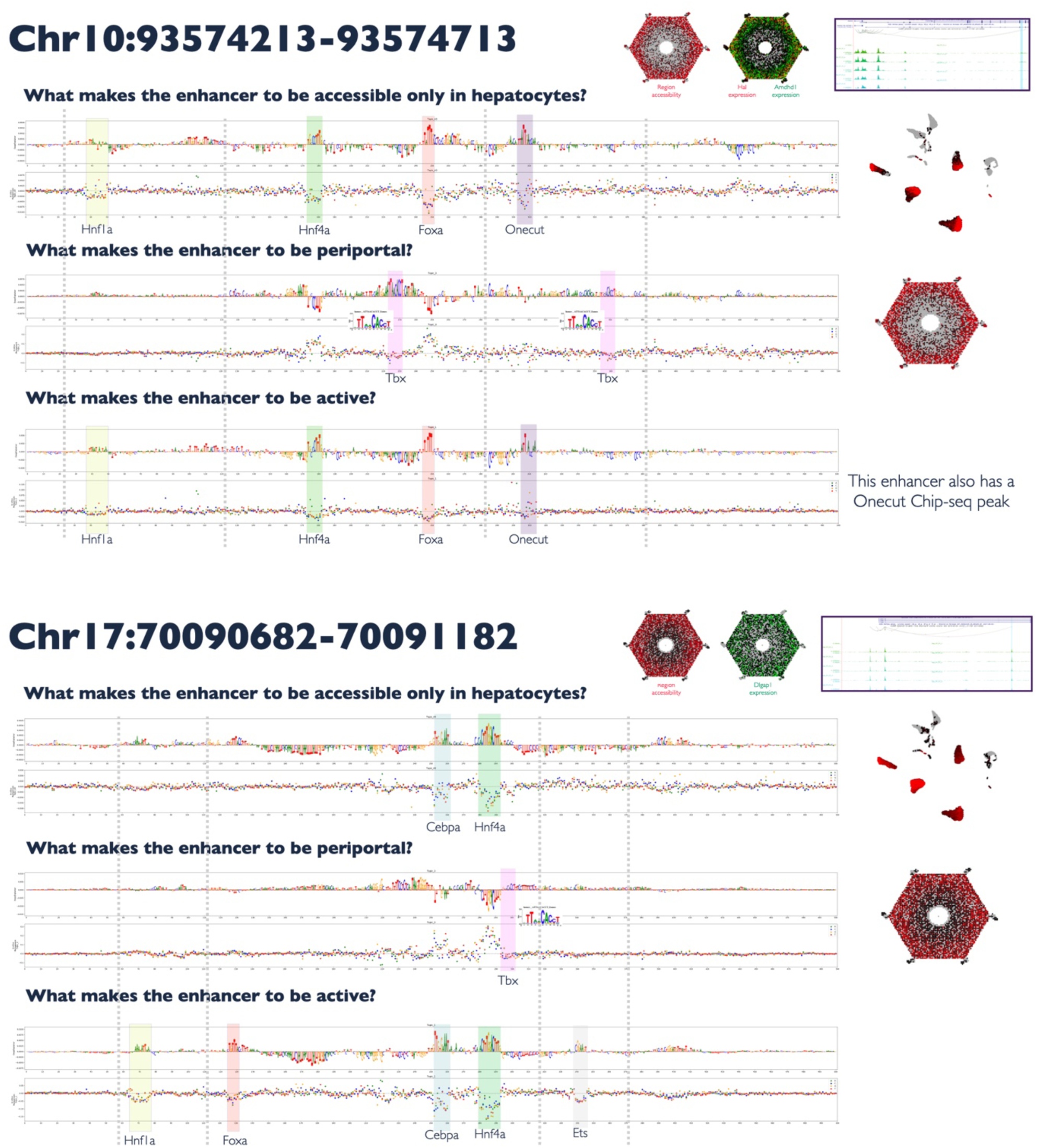
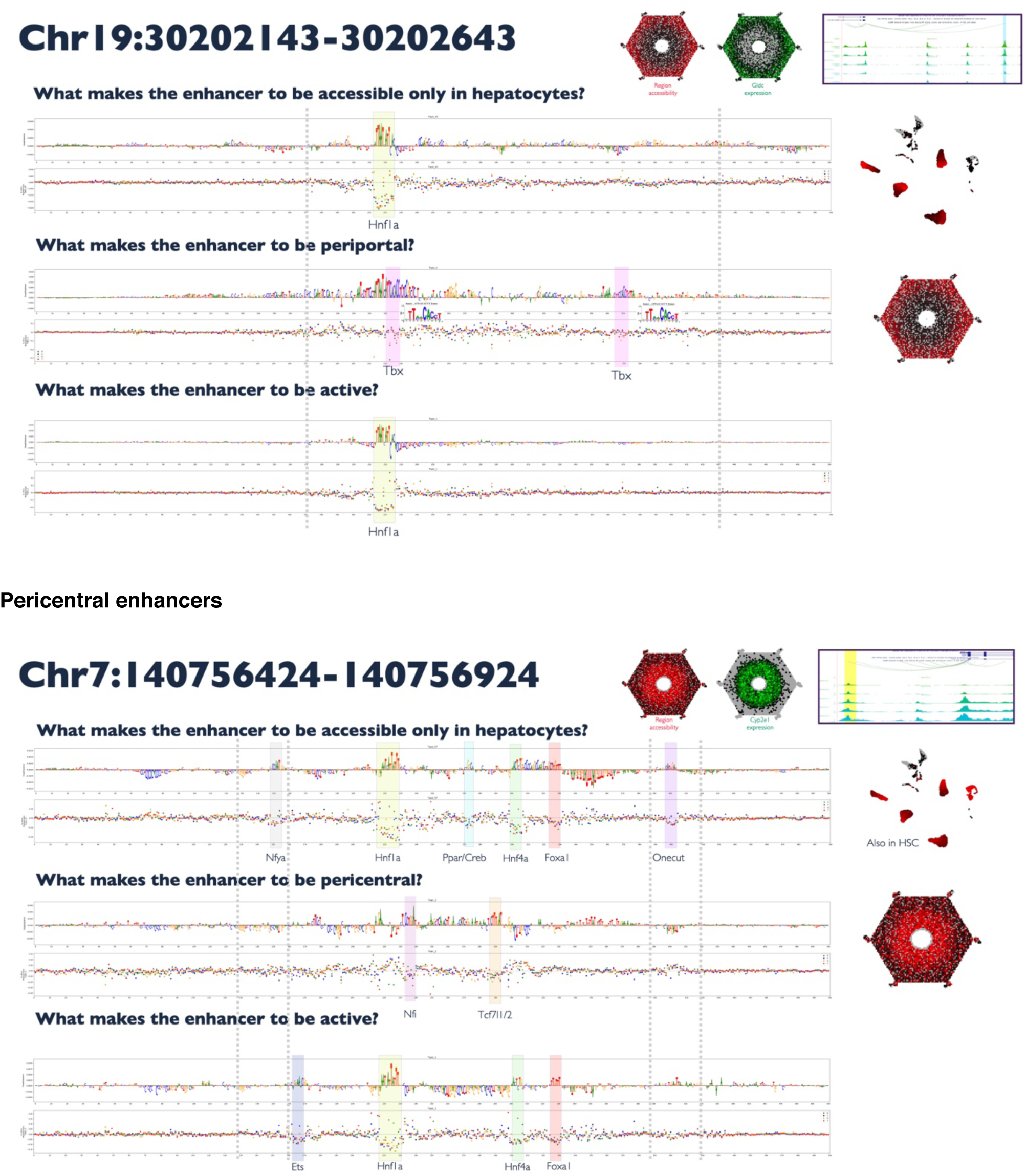
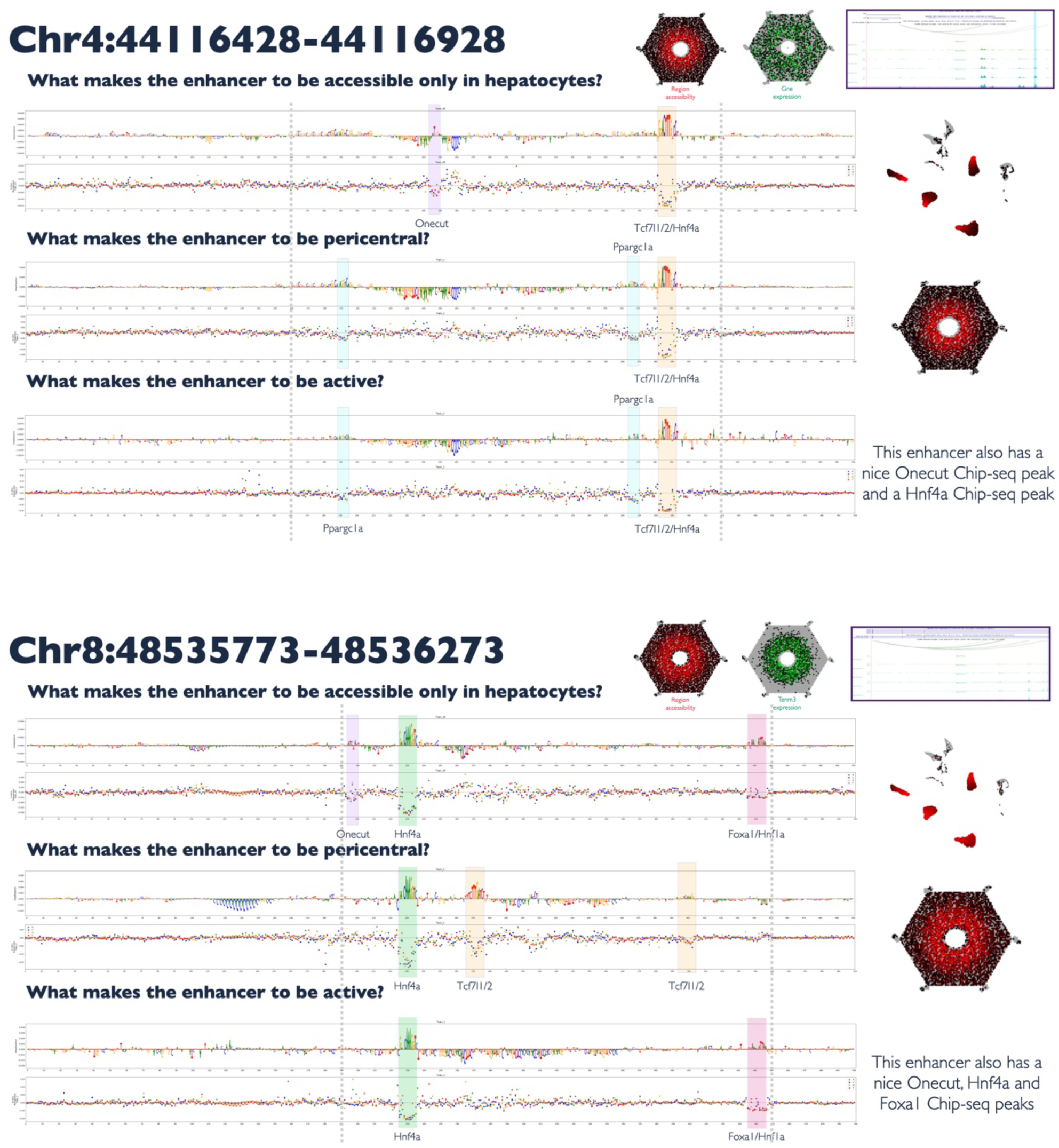
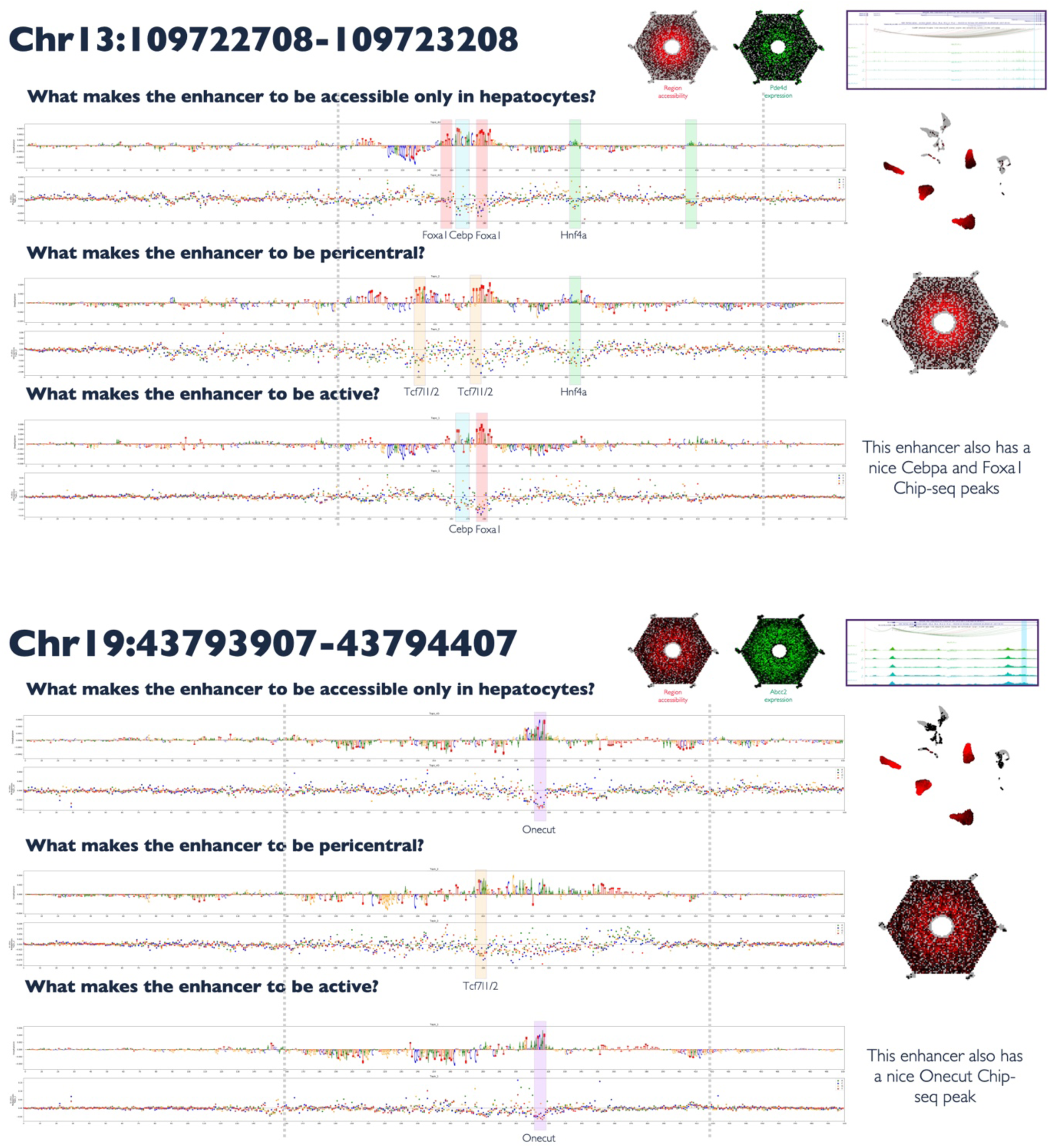
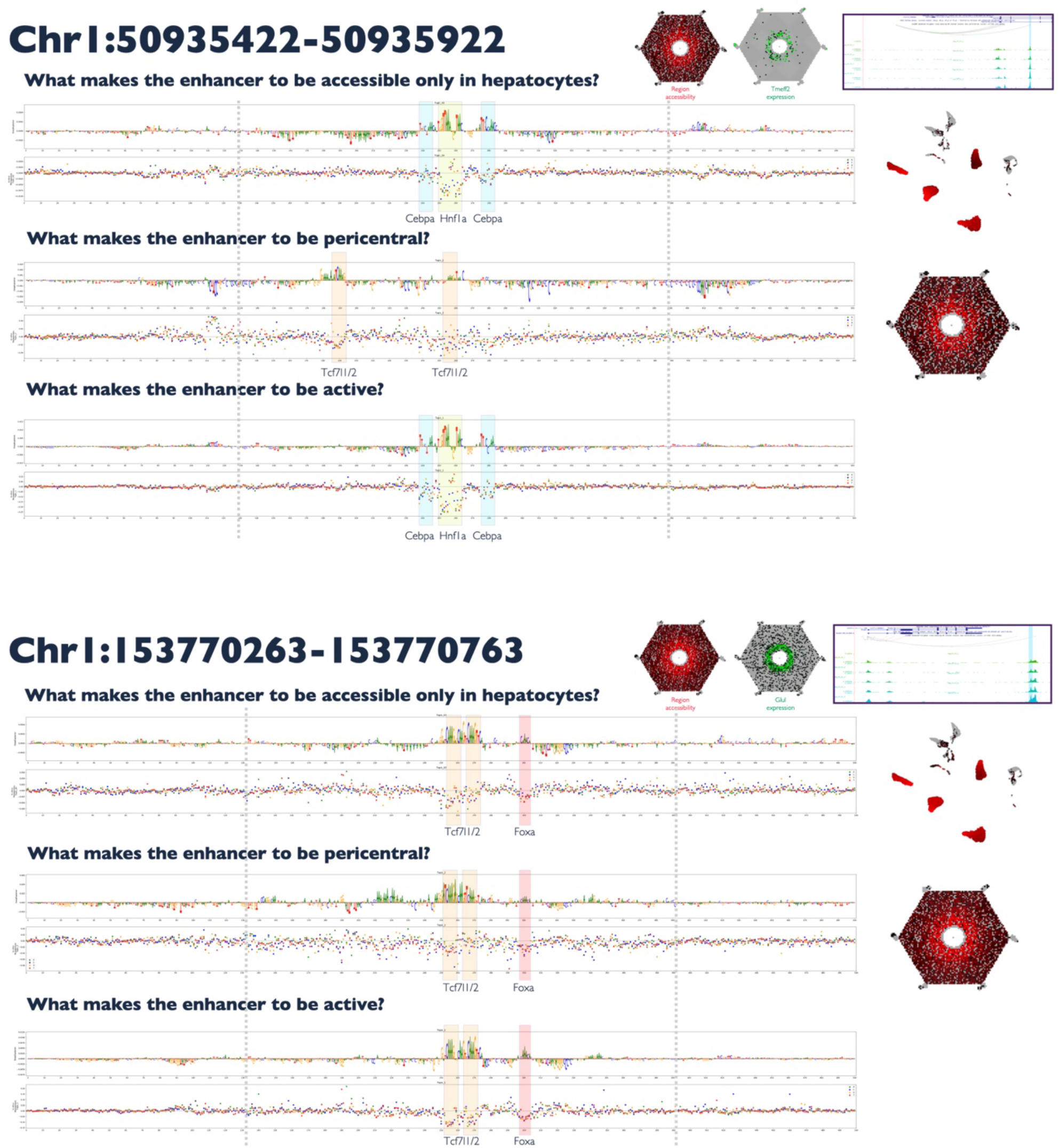
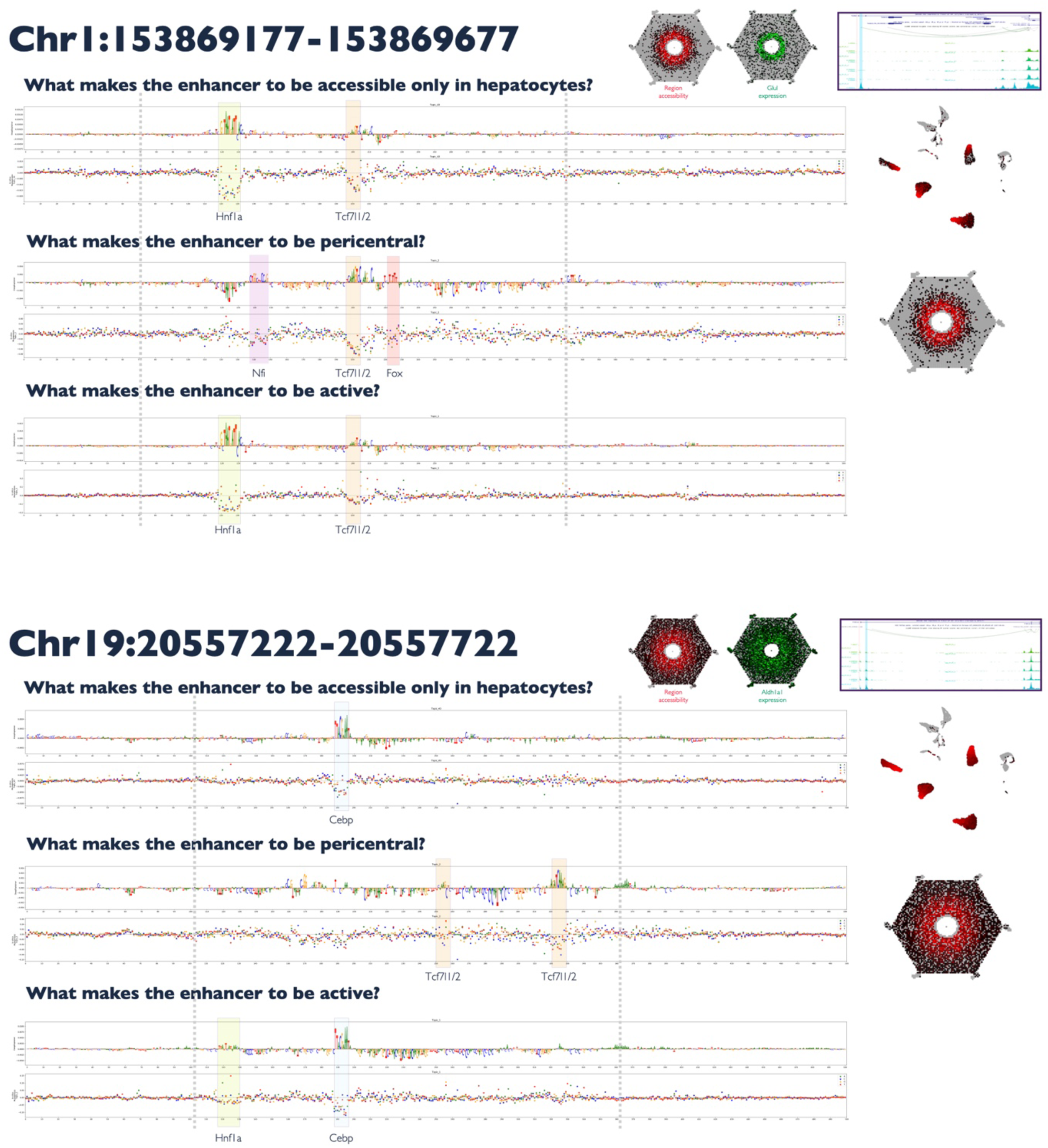
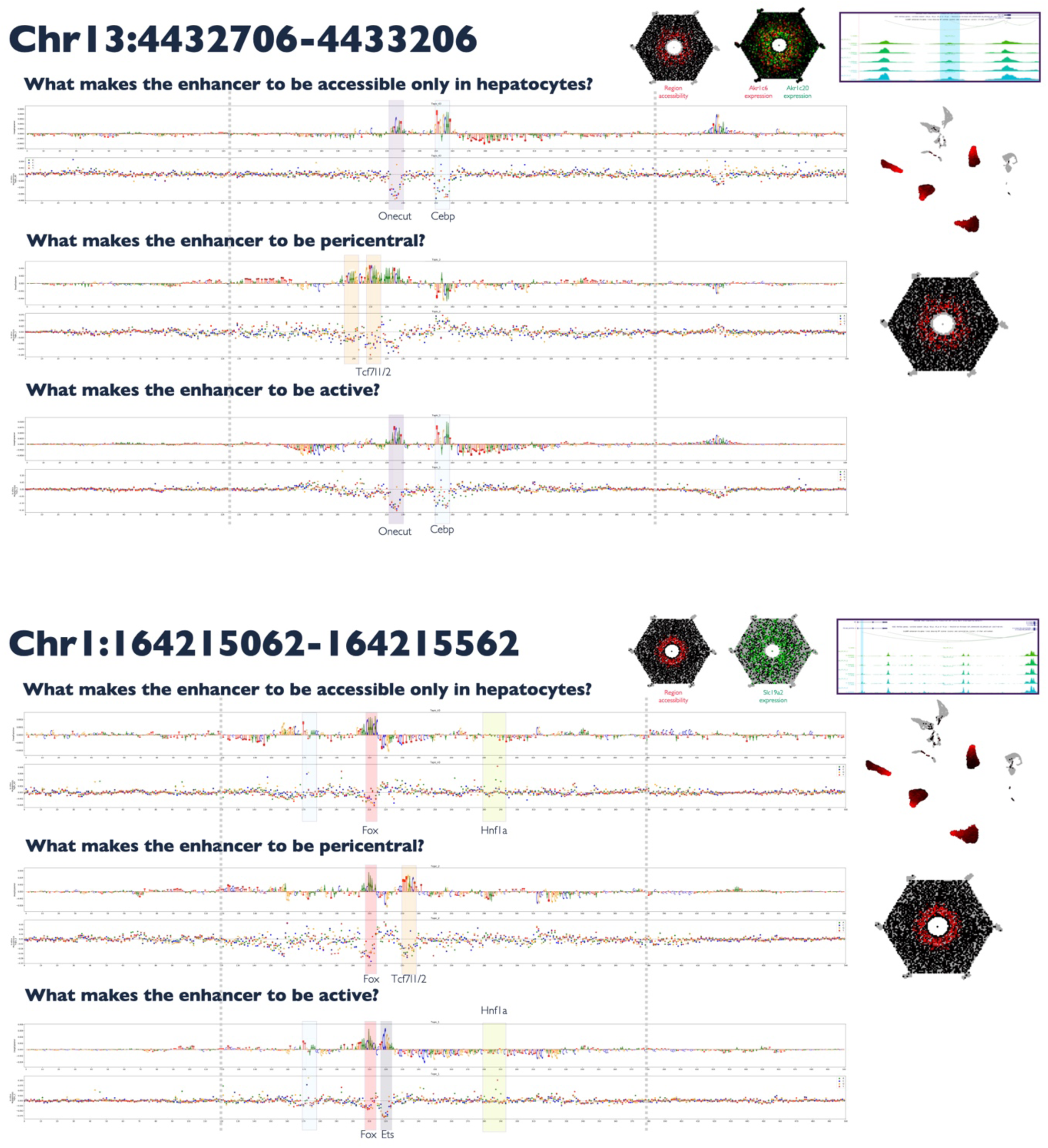
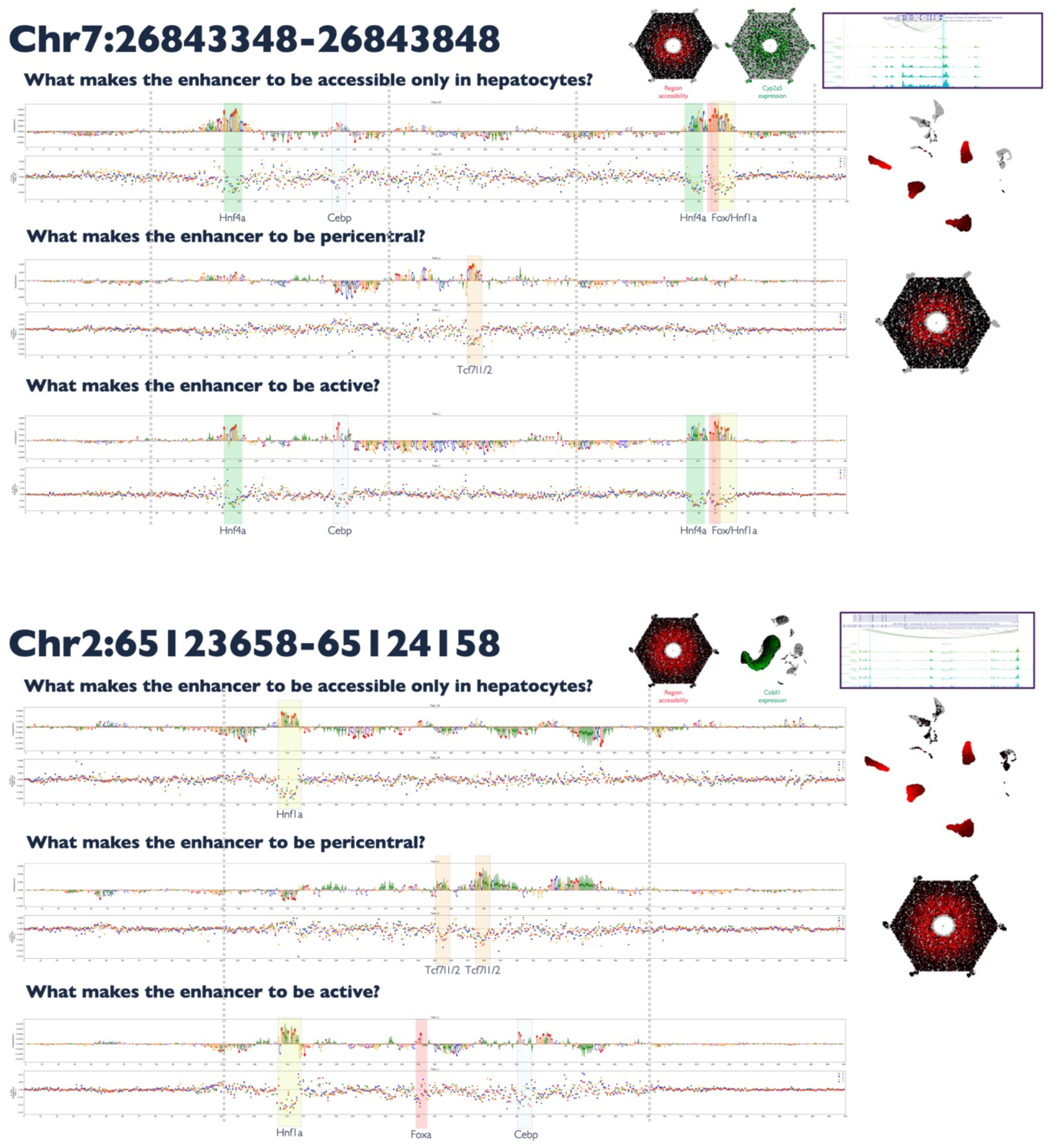
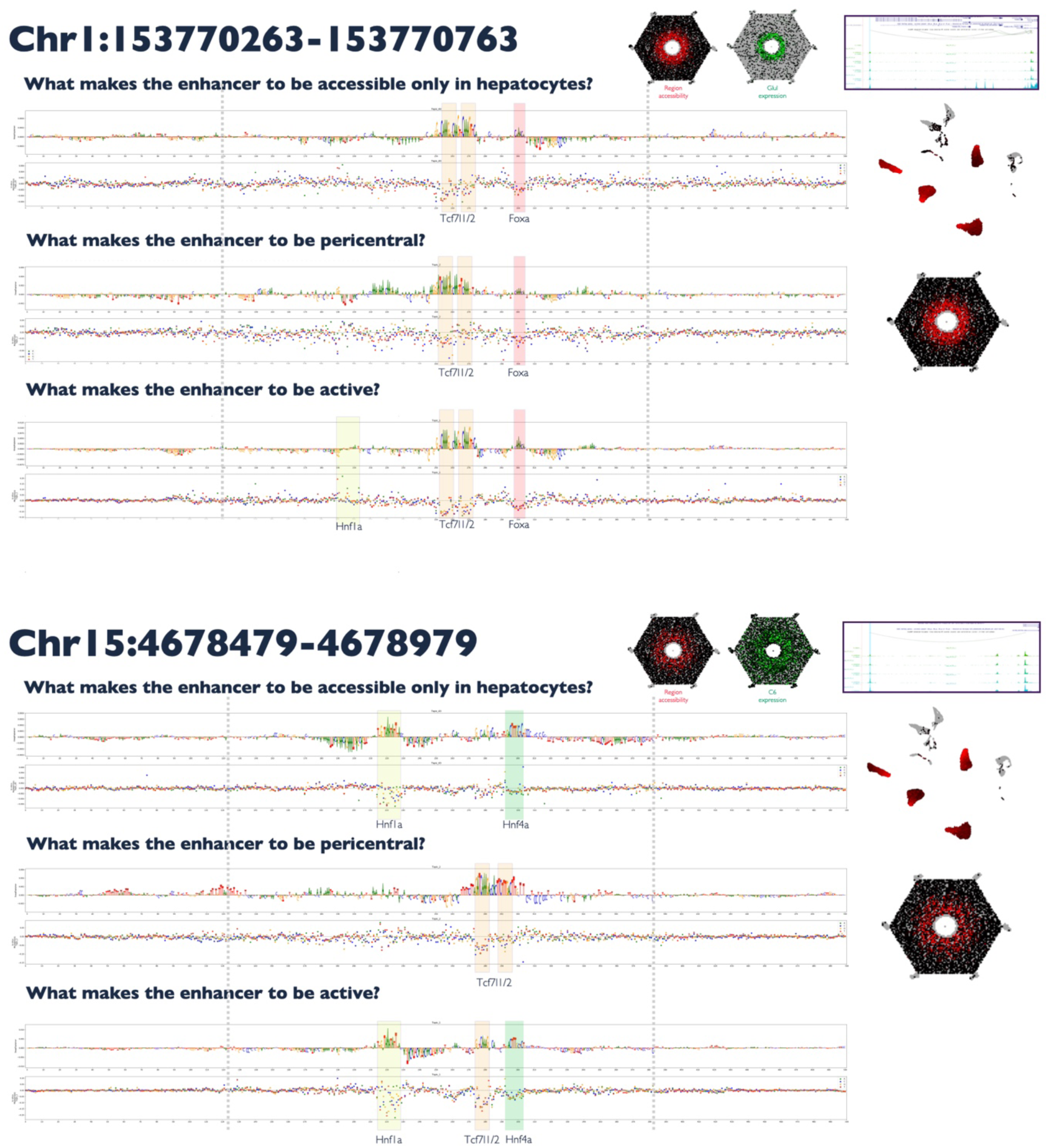
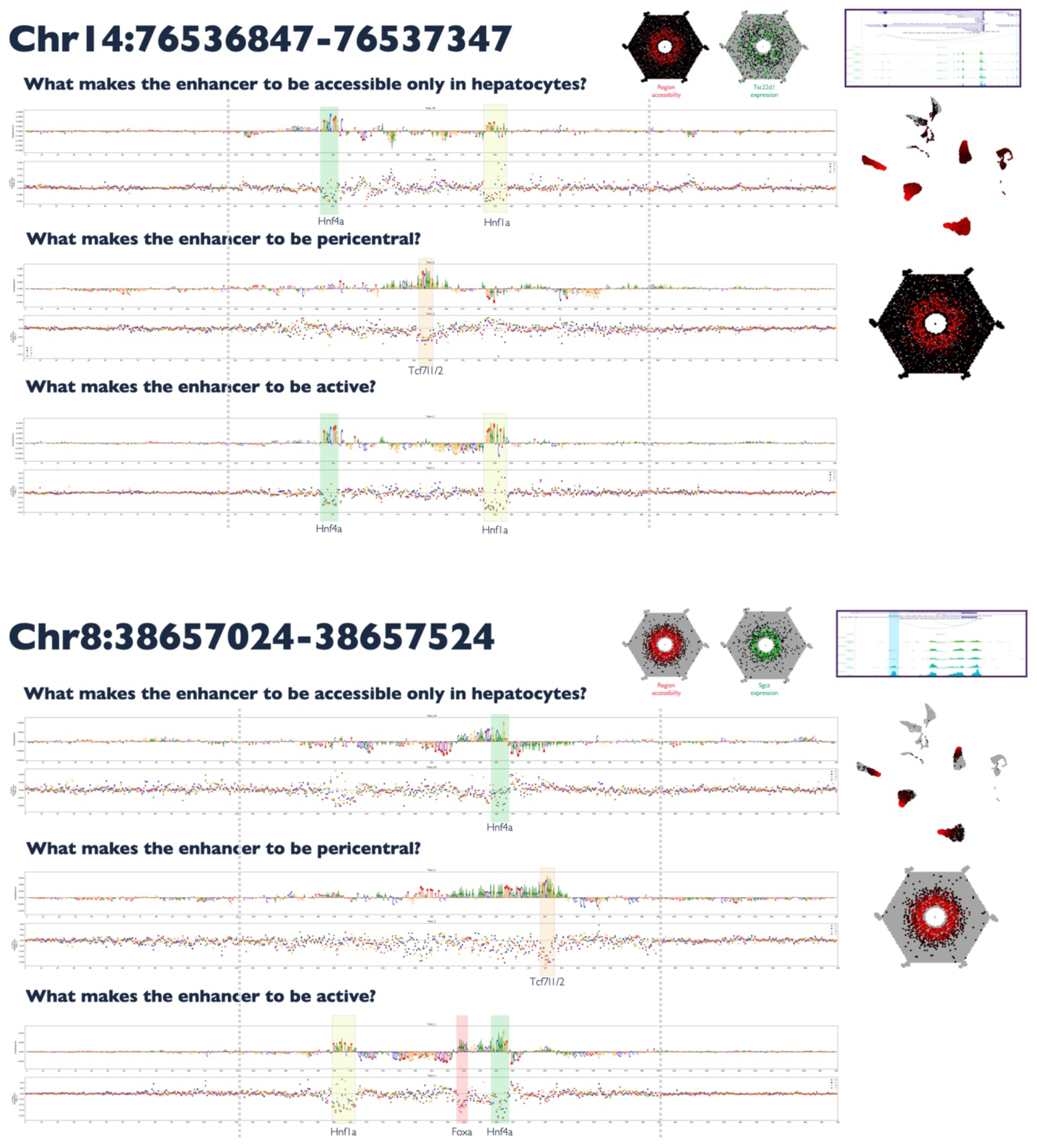
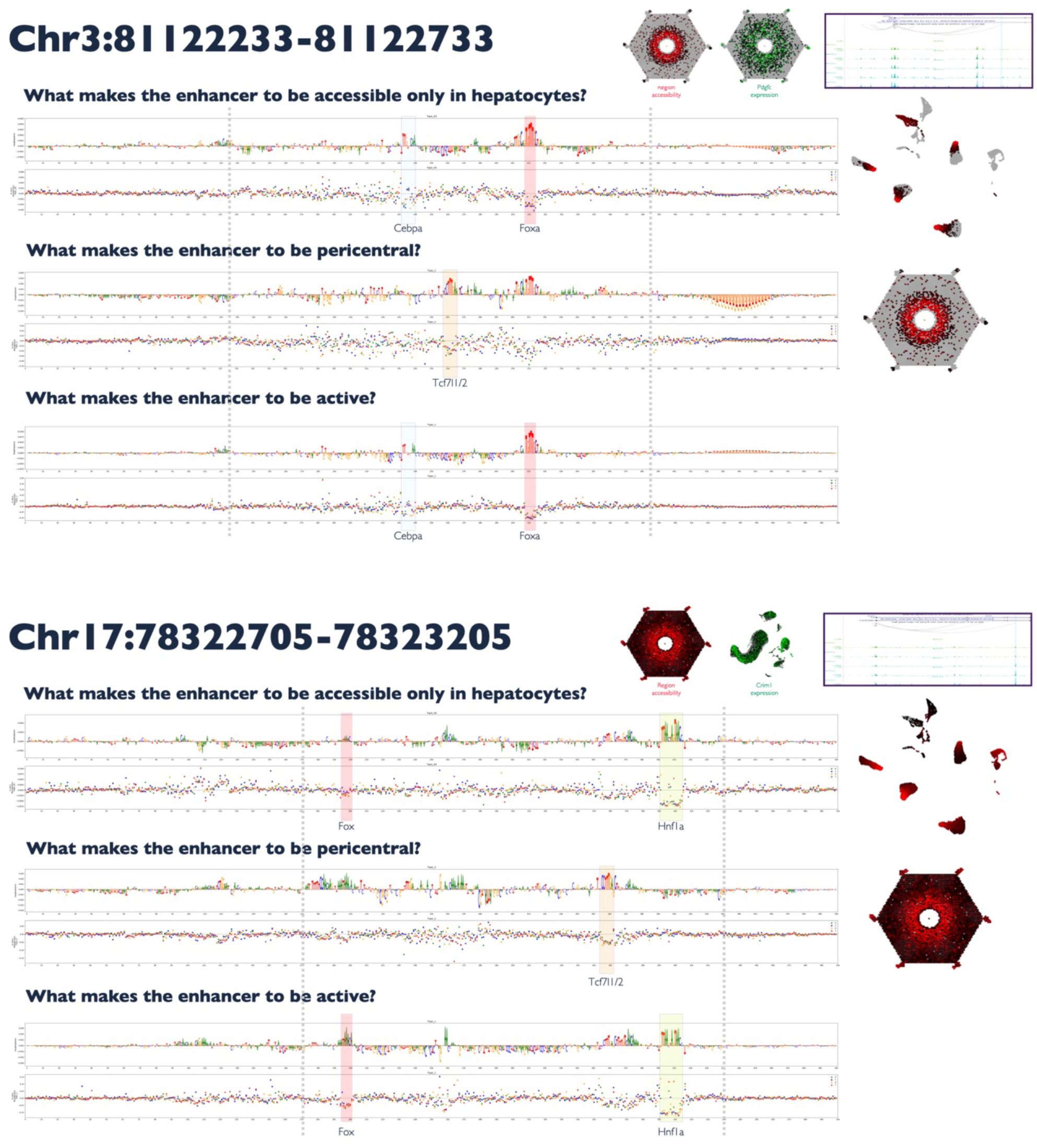
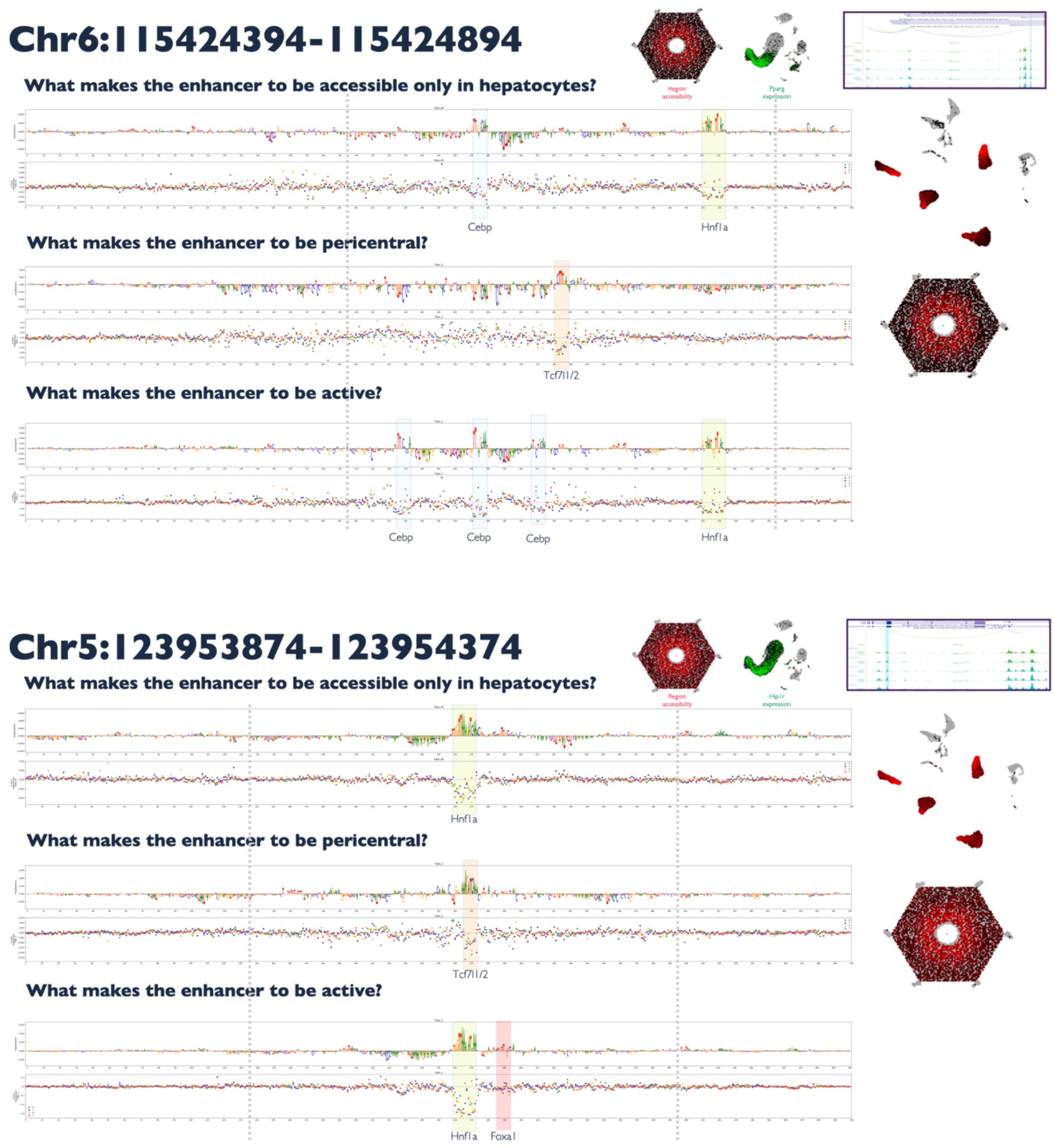

